# sciCNV: High-throughput paired profiling of transcriptomes and DNA copy number variations at single cell resolution

**DOI:** 10.1101/2020.02.10.942607

**Authors:** Ali Mahdipour-Shirayeh, Natalie Erdmann, Chungyee Leung-Hagesteijn, Rodger E. Tiedemann

## Abstract

Chromosome copy number variations (CNVs) are a near-universal feature of cancer however their effects on cellular function are incompletely understood. Single cell RNA sequencing (scRNA-seq) can reveal cellular gene expression however cannot directly link this to CNVs. Here we report new normalization methods (RTAM1 and −2) for scRNA-seq that improve gene expression alignment between cells, enhancing gene expression comparisons and the application of scRNA-seq to CNV detection. We also report sciCNV, a pipeline for inferring CNVs from RTAM-normalized data. Together, these tools provide dual profiling of transcriptomes and CNVs at single-cell resolution, enabling exploration of the effects of cancer CNVs on cellular programs. We apply these tools to multiple myeloma (MM) and examine the cellular effects of cancer CNVs +8q. Consistent with prior reports, MM cells with +8q22-24 upregulate MYC, MYC-target genes, mRNA processing and protein synthesis, verifying the approach. Overall, we provide new tools for scRNA-seq that enable matched profiling of the CNV landscape and transcriptome of single cells, facilitate deconstruction of the effects of cancer CNVs on cellular reprogramming within single samples.

## INTRODUCTION

Aneuploidy and focal CNVs are a pervasive feature of cancer^1^. However, the specific influence of many cancer CNVs on the transcription programs and molecular processes within cells remains only partially understood^2,3^. While may studies have sought to integrate cancer CNVs with gene expression effects by examining differences across tumor samples, such studies are often impeded by the extensive genetic heterogeneity that exists between tumors, confounding the attribution of transcriptional variances to single genetic loci. As a consequence, integrative inter-tumor studies commonly require the profiling of large numbers of samples to minimize confounding variations, and their application to less common cancer types or CNVs may be problematic.

To assess the transcriptional effects of specific CNVs, examination of the intra-clonal effects of emergent CNV at single cell resolution would have the advantage that subclonal cells with and without CNV are otherwise virtually isogenic, and that gene expression changes could be more readily attributed to the CNV. Therefore, simultaneous sequencing of single cell genomes and transcriptomes should in theory enable direct pairing of CNVs with their transcriptional outcomes with high precision. Notably, sequencing of both genomic DNA and RNA within single cells has been reported^4–6^. Prior techniques, however, provided profiling of only a few cells and thus afford only a limited view of the heterogeneity within any cancer. For the identification of CNV function, sequencing of much larger numbers of cells is required to capture intra-tumor subclones and to adequately profile subclonal populations for the gene expression pattern that aligns with an individual CNV.

Whilst scRNA-seq can reveal the transcription state of single cells, it cannot directly relate this to DNA lesions. CNVs can be inferred from scRNA-seq, which could thus be leveraged to provide both layers of omics information within individual cells. However, previously reported approaches^7–9^ reveal constraints imposed by the sparsity of single-cell data and have not reported quantitative measures of accuracy. In particular, inconsistencies in the detection of lowly-expressed genes within single cells causes stochastic noise that influences transcriptome distribution and interferes with RNA-based CNV detection. Normalization is thus critical for accurate scRNA-seq interpretation^10–14^ and for secondary CNV detection.

Here we report RTAM1 and RTAM2 normalization methods for scRNA-seq that improve gene expression alignment between cells and thus enhance the sensitivity of scRNA-seq for the detection of small expression changes arising from gene copy number differences. We also report sciCNV, a new pipeline for inferring CNVs in single cells from RTAM-normalized scRNA-seq data. Together, these methods enable high-throughput paired profiling of both DNA copy number and RNA in the same cells, facilitating direct intra-clonal examination of the effects of cancer CNVs on gene expression programs at a cellular level.

## RESULTS

### Enhanced single-cell RNA-seq normalization methods: RTAM1 and −2

Single-cell RNA-seq enables gene expression comparisons between cells. However, the accuracy of these comparisons depends critically upon data normalization. Normalization is essential to convert raw RNA transcript data from single cells into gene expression values that are objectively comparable between cells, eradicating biases related to cellular sequencing depth, transcriptome size, technical factors or batch effect. As the best methods for normalizing scRNA-seq remain controversial, each with its own drawbacks, we developed RTAM1 and −2 (described in the **online methods** and **supplementary figures S1-3**) and compared the RTAM methods with other normalization strategies currently in use.

To compare the methods for their control of systemic and stochastic variations between cells due to size or sequence depth, we generated scRNA-seq data for normal and malignant cells belonging uniformly to the B cell lineage (n>15,000 cells) (**figure 1a**). We examined cells with a similar ancestry in order to minimize confounding variations between them due to lineage specification. However, we deliberately generated data from a mix of both small quiescent B cells and large proliferating plasma cells to ensure that the normalization methods would be challenged by cells embodying a full spectrum of sizes and transcriptional activities. The cells were isolated from human bone marrow samples by FACS and were profiled using the 10X Genomics single cell RNA-seq library kit. Cell- and gene-specific transcripts were enumerated using barcoded unique molecular identifiers (UMI).

**Figure 1.**
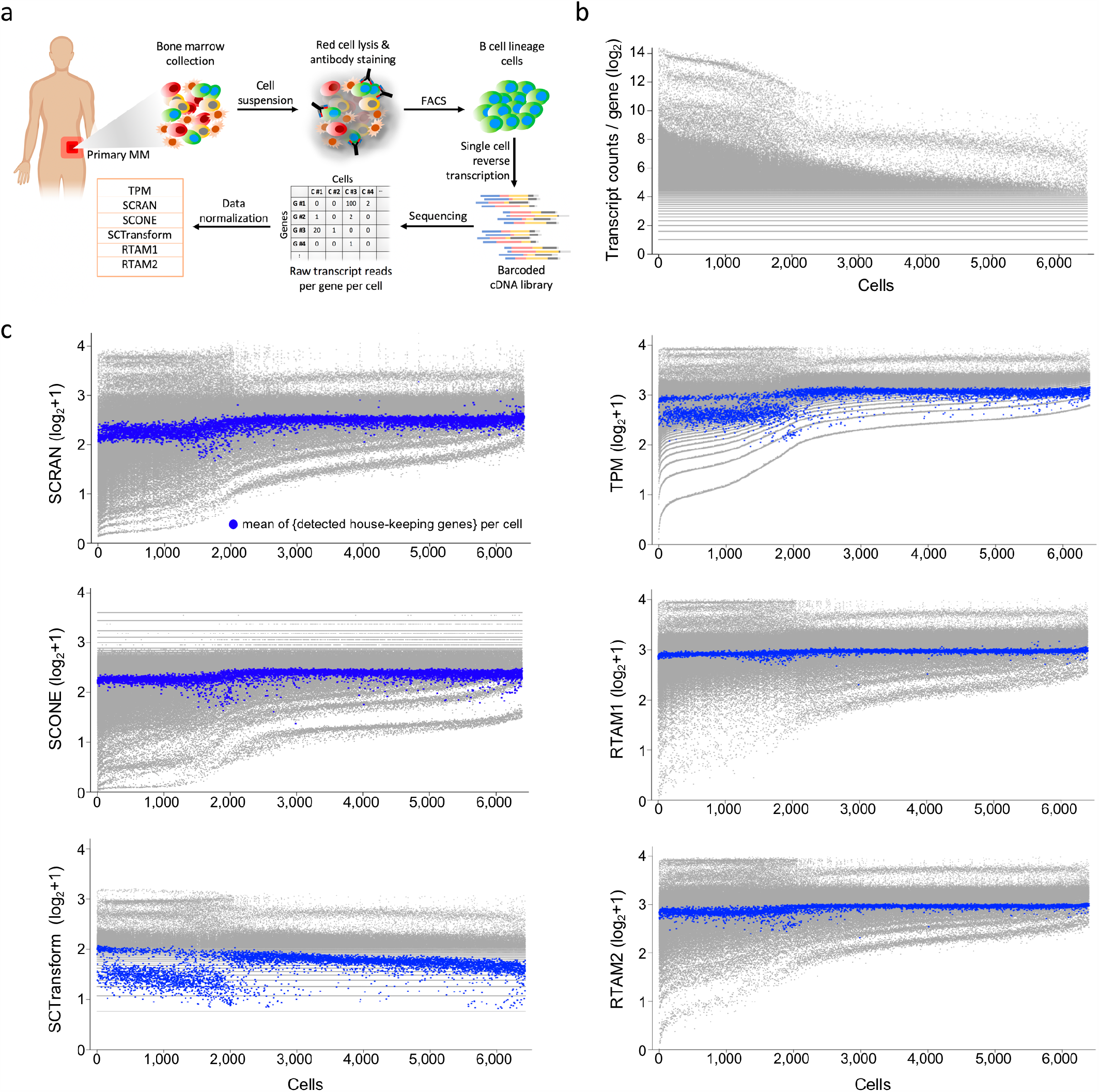
Comparison scRNA-seq normalization strategies, including RTAM1 and RTAM2 methods. **a.** Overview of workflow showing bone marrow sampling, FACS sorting of B cell lineage cells, scRNA-seq and data normalization. MM, multiple myeloma. FACS, fluorescence activated cell sorting. **b.** Plot of scRNA-seq data from >6,000 B cells and plasma cells, isolated from the bone marrow of a MM patient, depicting the raw (pre-normalized) transcript counts per gene per cell. Each dot represents an integer transcript count for one or more genes in a single cell; cells (columns) are ranked from left to right by their total transcript count. **c.** The same data is shown following normalization using TPM, SCRAN, SCONE, SCTransform, RTAM1 or RTAM2 methods (and following log transformation). To compare the methods, the mean expression of a curated set of house-keeping genes (HKG) is plotted in each cell as a blue dot. See also **supplemental figures S4-S20**.

The raw scRNA-seq data from one of the test samples is depicted in **figure 1b**. As shown, the distributions of transcript numbers per gene varied significantly from cell to cell, predominantly reflecting differences in the cellular transcriptome sizes and demonstrating a clear need for normalization. The test samples were next normalized using either TPM^15^, SCRAN^11^, SCONE^12^ or Seurat’s SCTransform function^16^ (**figure 1b** and **supplementary figures S4-S20**). To compare the alignments of the normalized transcriptomes, we examined the mean and median expression in each cell of a curated list of housekeeping genes (HKG) known to be broadly expressed with low variation^9^. We also examined the average expression in each cell of all of the ubiquitously-expressed genes (UEG) detected in >95% of the cells in the sample. For each sample tested, the UEG represent the largest possible set of genes that are commonly expressed across the test cells. Whilst the expression of any individual gene is expected to vary between cells for both biological and technical reasons, the average expression per cell of a large set of ubiquitous genes should be similar, particularly amongst cells of the same lineage, and its variance between cells provides a metric of normalization effectiveness.

As shown in **figures 1c, 2** and **supplementary figures S4-20**, TPM, which normalizes cellular transcriptomes primarily by their total transcript count, produced a very large variance in the average expression of HKG or UEG between individual cells, suggesting significant limitations for scRNA-seq application. By comparison, SCRAN and SCONE produced superior alignment of gene expression averages across cells. However, SCONE, which produced the better alignment, achieved this only by implementing quantile normalization – exchanging the actual distribution of transcript counts in each cell for a standardized distribution – which caused a significant loss of inter-cellular variation, particularly in key highly-expressed genes. The expression of IGH or IGL genes, for example, a critical feature of plasma cells, was reduced by quantile normalization into a virtual constant across cells (**supplementary figure S21**).

As each of these scRNA-seq normalization methods has limitations, we developed RTAM1 and - 2. The RTAM methods originate from a consideration of the strengths and weaknesses of scRNA-seq. Whereas lowly expressed genes are detected within single cells with low resolution (due to low integer transcript counts) and show significant stochastic variation, highly expressed genes are robustly detected and show finer quantisation of variation relative to intensity. RTAM1 and −2 thus utilize highly-expressed genes, whose expression is resolved with greater accuracy, to align cellular transcriptomes. Genes are ranked in each cell by their expression and the summed intensities of the top-ranked genes is standardized in log-space using unique non-linear cell- and gene-specific adjustments of gene expression determined either by cellular gene expression rank (RTAM1) or by gene expression intensity (RTAM2) (see **methods**).

Importantly, compared to TPM, SCTransform or SCRAN, both RTAM1 and RTAM2 reduce the cell-to-cell variance in the average (median or mean) expression of HKG and UEG sets (**figure 2** and **supplementary figures S4-S20**). The coefficients of variation (CV) produced by each normalization method for the “average” expression of HKG or UEG in individual cells is shown in **figure 2**, for 3 independent patient samples. As shown, RTAM1 and RTAM2 reduce variations in the average gene expression of single cells, irrespective of whether this average expression is calculated by any of 3 different methods. By design, the RTAM methods also standardize the average expression of highly-expressed genes, and thus overall these methods produce superior alignments of cellular transcriptomes and of gene expression between cells. At the same time, both RTAM1 and RTAM2 maintain the original variability observed between cells in the expression of individual highly-transcribed genes, unlike the quantile normalization implemented by SCONE (**supplementary figure S21**). Overall, therefore, the RTAM methods represent useful new strategies for normalizing scRNA-seq data that can enhance the accuracy of gene expression comparisons between cells.

**Figure 2.**
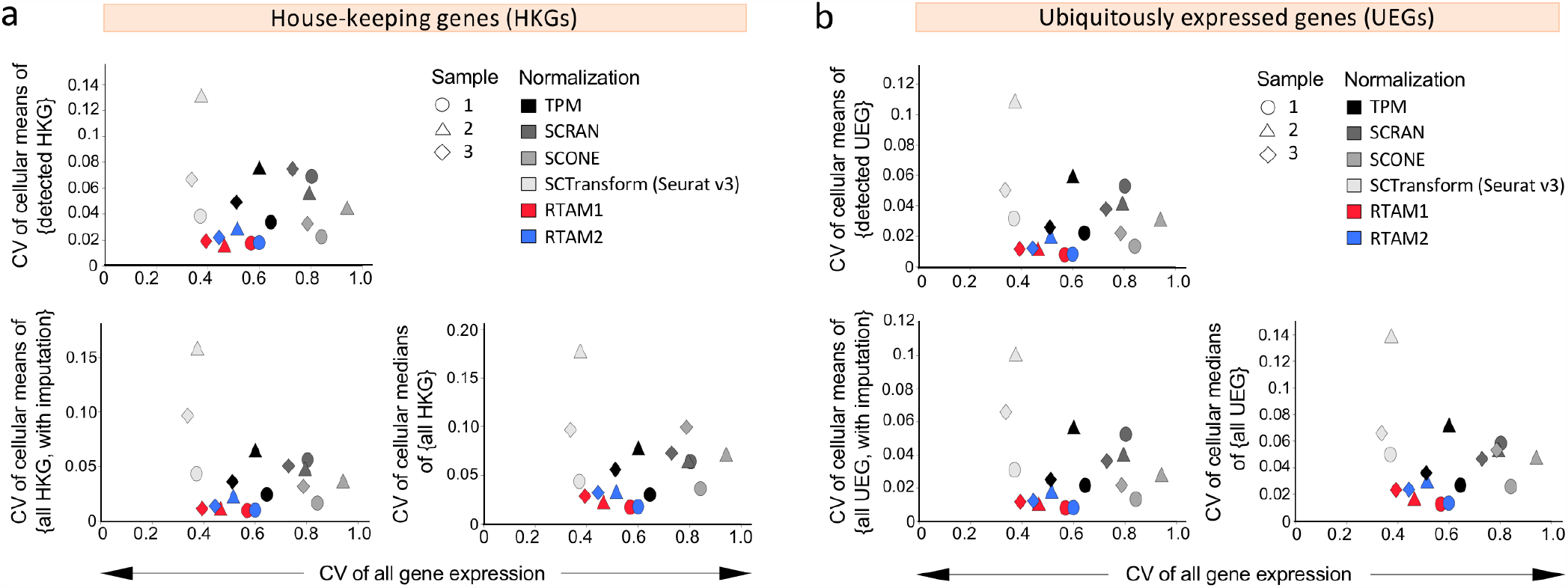
RTAM scRNA-seq normalization reduces cell to cell variation in average expression of house-keeping genes (HKG) and ubiquitously-expressed genes (UEG). Alignment of single cell transcriptomes is compared using 6 different methods for scRNA-seq normalization. **a.** The coefficient of variation (CV) across cells in the average expression of HKGs within each cell is shown for 3 scRNA-seq samples (containing >15,000 cells), normalized by 6 different methods. To avoid bias in the analysis, average expression was alternatively calculated as either the mean of the detected HKG (top panel), the mean of all HKG [with imputation of null dropout values] (bottom left panel), or the median of all HKG without imputation (right panel). As the various normalization methods expand or compress the distribution of the overall gene expression data to different extents, the CV of HKG averages is plotted against the CV of expression of all genes. **b.** The coefficient of variation (CV) for all genes ubiquitously expressed (UEGs)(in >95% cells) was determined for the same patient samples. Once again, the average expression of UEG in each cell was alternatively calculated as either the mean of the detected UEG (top panel), the mean of all UEG imputing null dropouts (bottom left panel), or the median of all UEG without imputation (right panel). A lower CV in the mean expression of HKG/UEG implies better alignment of average gene expression across cells. As the normalization methods expand or compress the distribution of the overall gene expression data to different extents, the CV of HKG/UEG averages (y-axis) is plotted against the CV of expression of all genes (x-axis).

### Single-cell inferred chromosomal copy number variation: sciCNV

We next sought to develop a method for detecting single-cell chromosomal CNV from scRNA-seq, leveraging the enhanced normalization provided by RTAM to increase the sensitivity of single-cell transcriptomics for CNV detection. To optimize DNA copy number estimates from gene expression, and to mitigate against data sparsity in single cells, we developed a two-pronged approach, called sciCNV (described in the **methods**). Briefly, for each cell RTAM-normalized gene expression was aligned with that of control diploid cells to develop expression disparity scores, which were averaged in a moving window across the genome. The average gene expression of the control cells was weighted by the probability of gene detection, enhancing its comparison with single cell data, where signal ‘*dropout*’ was common for many genes. In a parallel method, the expression disparity values for each gene were exchanged for binary +1 or - 1 values, which were summed cumulatively in each cell as a function of genomic location; the gradient of this function yielded a second estimate of the single cell CNV profile that was sensitive to small concordant expression variations in contiguous genes and that was insensitive to large single-gene variations. The CNV estimates of the two methods were then combined by their geometric mean.

**Figure 3** shows sciCNV applied to scRNA-seq data from primary multiple myeloma (MM) cells. Significantly, the CNV profile of a single MM cell, inferred from its RNA, closely resembles the average CNV profile of >10^4^ tumor bulk cells, derived from whole exome DNA sequencing (WES) of the same MM sample (R^2^=0.72) (**figure 3a**,**b**). In addition, the CNV predictions produced from a single cell by sciCNV also aligned with CNV estimates provided by FISH (**figure 3c**). Furthermore, examination of >1,700 individual plasma cells from the same MM patient biopsy using sciCNV revealed that the tumor clone CNVs were robustly detected in each of the MM cells (**supplementary figure S22a**), despite biological and technical variations between them; and were not detected in normal plasma cells (NPC). Thus, sciCNV can utilize scRNA-seq data to reveal CNVs in single cells. Moreover, it can distinguish cancer cells and normal cells on the basis of their CNV profile (**supplementary figure S22b**,**c**).

**Figure 3.**
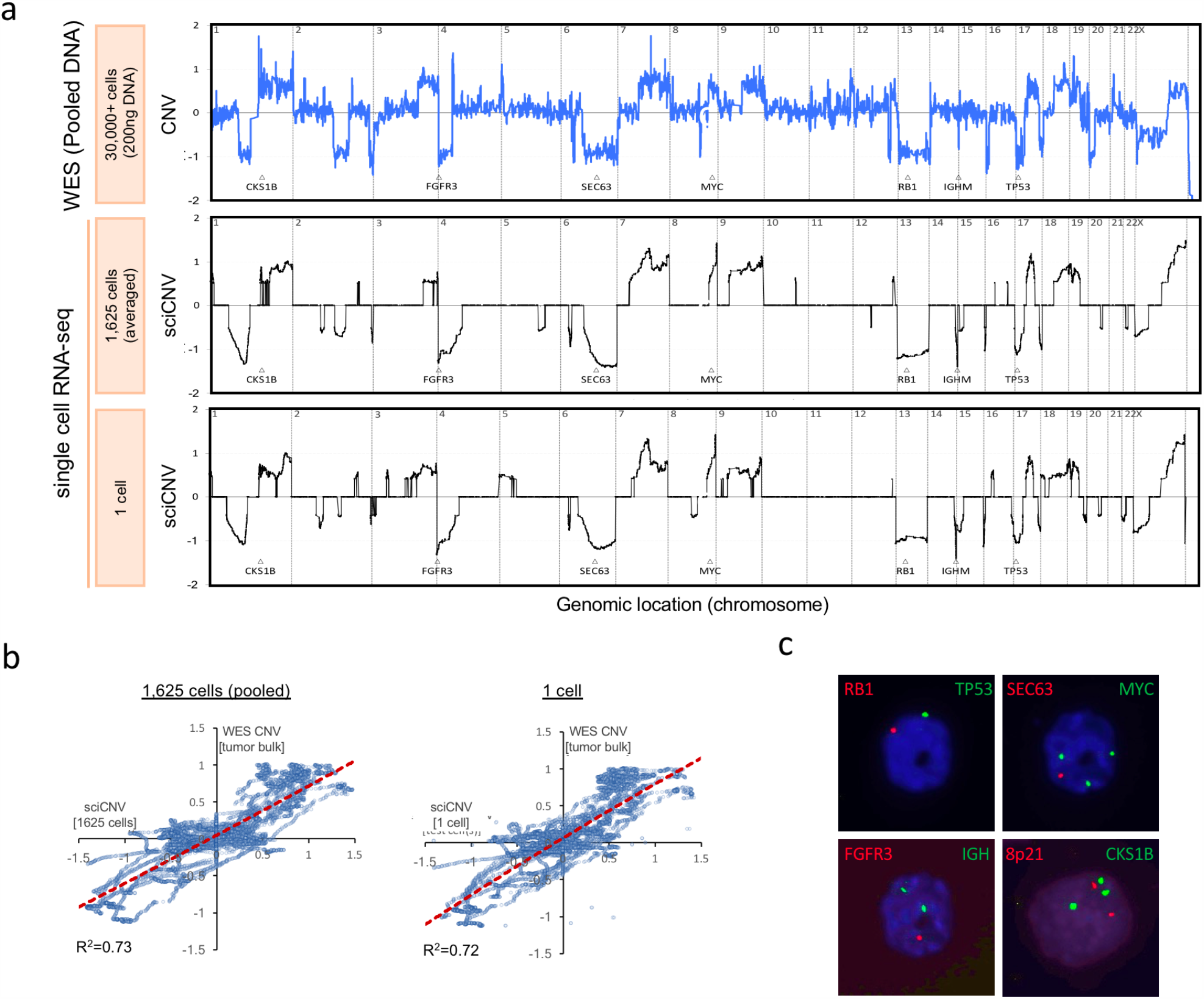
sciCNV can predict DNA copy number variations (CNV) in single cells from scRNA-seq data. **a.** The CNV profiles of 1,625 pooled tumor cells (middle panel) or of a single tumor cell (lower panel) from sample MM199 were inferred from scRNA-seq using RTAM2/sciCNV and are shown compared with the bulk tumor CNV profile (upper panel), which was obtained by whole exome sequencing (WES) of DNA purified from 1.9×10^6^ cells. Cells were isolated from bone marrow by FACS. **b.** The scRNA-seq-based sciCNV profiles from (**a**) are correlated with the tumor bulk CNV profile derived from WES. **c.** FISH verification of sciCNV copy number predictions, for the genes highlighted in **a**., showing 3 copies of CKS1B (1q21), 1 copy of FGFR3 (4p16), 1 copy of SEC63 (6q21), 2 copies of PNOC (8p21), 3 copies of MYC (8q24), 1 copy of RB1 (13q14,) and 1 copy of TP53 (17P13), as predicted. Brightness and contrast were adjusted during figure construction to enhance probe visualization.

To further examine the accuracy and consistency of sciCNV in detecting tumor CNVs in single cells, we next compared the sciCNV profiles of thousands of individual plasma cells from 4 MM samples with the corresponding tumor CNV profiles generated from DNA sequencing of >10^6^ tumor cells (**figure 4**). Remarkably, the single cell CNV profiles calculated by sciCNV from RNA substantively recapitulated the complex CNV landscape of each tumor clone, defined by tumor bulk DNA sequencing. Thus, for the MM199 sample, gains within 1q, 7q, 8q, 9, 17q, 18q, 19p and Xq, and losses within 1p, 2q, 4p, 6p, 6q, 13, 14q, 16p, 17p, 17q, and Xp were all identified by sciCNV at a single cell level, and these corresponded to CNVs identified by tumor bulk WES (**figure 4a**). Similarly, for sample MM241, trisomies of chromsomes 5, 7 and 19 and monosomy of chromosome 13 were all identified by sciCNV in single MM cells, matching the WES-defined CNV profile of pooled tumor bulk cells (**figure 4b**). Furthermore, sciCNV also identified a subpopulation of MM241 cells with subclonal gain of 1q, and the existence of this subclone appeared to be borne out by the tumor DNA WES profile which also identified a 1q-arm copy gain with an average CNV peak height of approximately 35%. Similarly, sciCNV profiling of copy number variations in individual cells from sample MM244, developed using RNA-seq data, identified clonal gains within chromosome 7 and clonal losses within chromosomes 1p, 10p, 13, 14q, 16q and 17p that were also identified by DNA sequencing of pooled tumor bulk cells from the same patient; and additionally identified subclonal loss of the distal arms of 1p and 2q, and subclonal gains of 11q and 19p (in subpopulations of cells) that were also detected at subclonal CNV levels (0.15-0.50) by WES (**figure 4c**). Finally, sciCNV identified clonal gains of 1q and 6p and clonal deletions within 1p, 2p, 6q, 13, 14q, 18 and 22 in single cells from MM254 that were all also identified on tumor bulk DNA sequencing (**figure 4d**). Thus, these data demonstrate that the genomic CNV landscape of individual tumor cells can largely be inferred from scRNA-seq transcriptome data by sciCNV, with results that show reasonable correlation to DNA-based CNV profiling of pooled tumor bulk cells, although the authors acknowledge that using RNA-based approaches small CNV regions may be missed.

**Figure 4.**
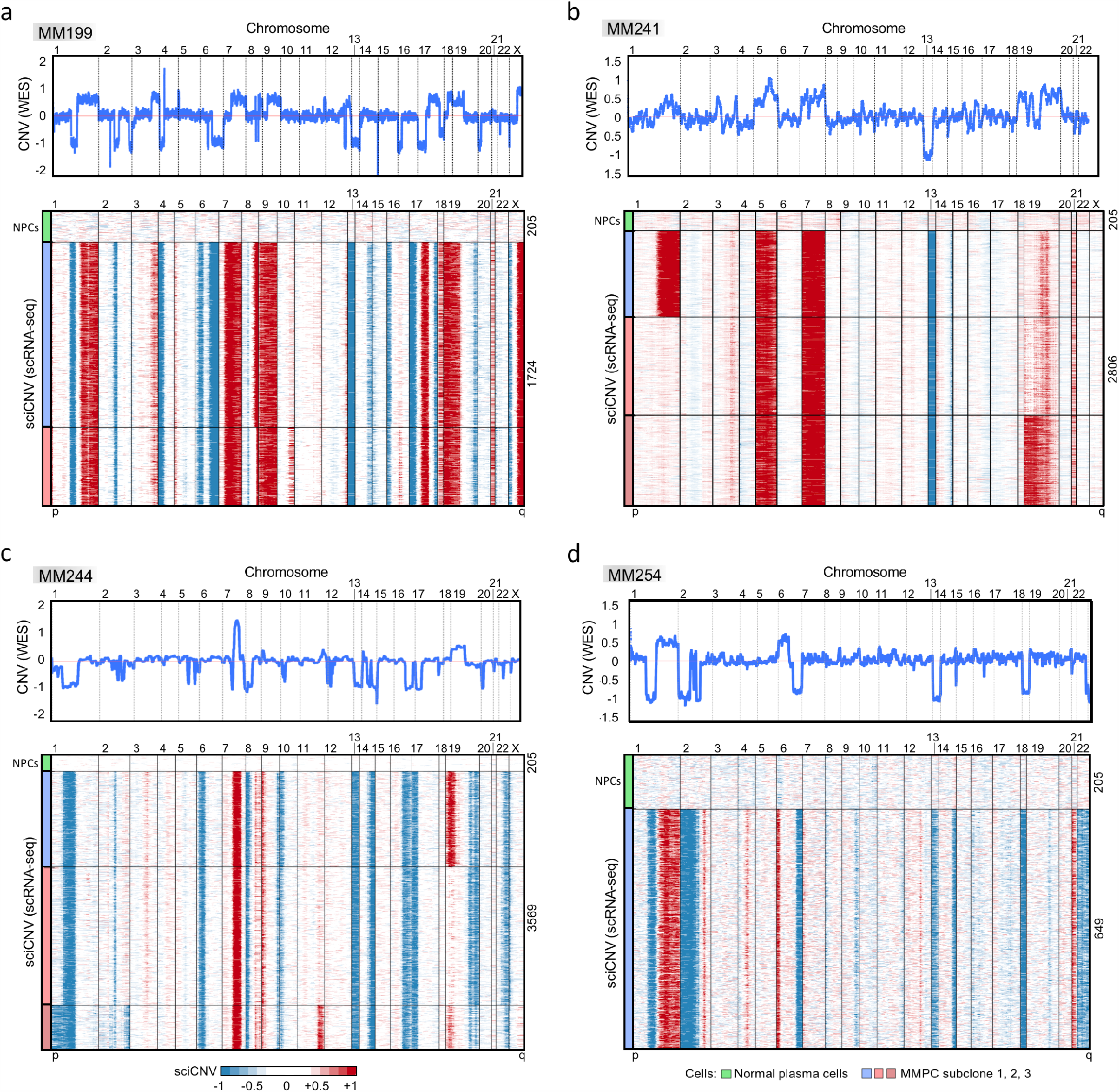
Using scRNA-seq data, sciCNV consistently infers single cell CNVs across thousands of cells that correlate with bulk tumor DNA-based CNV profiling. **a**.-**d**. The CNV profiles of thousands of individual plasma cells from MM patient samples MM199 (n=1724)(a), MM241(n=2806) (b), MM244 (n=3569)(c) and MM254 (n=649)(d) were inferred from scRNA-seq using RTAM2/sciCNV (lower panels) and are shown compared to the bulk tumor CNV profiles generated by DNA whole exome sequencing (upper panels). The sciCNV profiles of normal plasma cells (NPCs)(n=205) are shown as additional controls. For the heatmaps revealing CNV profiles, at single cell resolution, the tumor cells in each sample are grouped into inferred subclones (color bars at left) distinguished by divergent CNV.

Notably, several alternative methods exist for inferring CNVs from scRNA-seq^8,9,17,18^. Of these, inferCNV^17^ has been broadly used to calculate and visualize CNVs at single cell resolution. Next, therefore, we compared our RTAM2-sciCNV pipeline with inferCNV, applying both methods to the same set of normal and MM cells shown in figure 4 and comparing their single cell outputs with the CNV profiles obtained from tumor bulk WES (**supplementary figures S23-26**). As shown, sciCNV more reliably predicted tumor CNVs that were identified by WES, and did so more consistently across individual cells. For example, for MM199 cells sciCNV more readily detected gains of the distal arm of 1q, 3q, 7q, 9q and within 17q and 18q than inferCNV (**supplementary figures S23)**. At the same time sciCNV generally produced less false positive CNV noise than inferCNV. Thus, for MM241 both methods detected subclonal gain of 1q and trisomies of 5,7 and 19, and deletion of 13 and 14q, but inferCNV also reported a number of false positive deletions, particularly in chromosomes 2 and 4 (**supplementary figures S24)**. Overall, therefore, whilst both methods provide CNV predictions that are substantially correct, in general sciCNV offered a closer correlation with WES-based CNV findings.

### Dissecting the effects of CNVs on gene expression within single cells: the transcriptional consequences of +8q22-24 in MM

Simultaneous profiling of both DNA copy number and RNA in the same cell should enable examination of the effect of a CNV on transcriptional programs. To test this, we used sciCNV to study the effect of gain of the distal arm of 8q on cellular processes. We examined +8q as this is one of the most recurrent abnormalities in human cancer^2,17^ and is known to target MYC^18^, providing a benchmark result for testing our method.

Using sciCNV, sample MM199 was found to contain cells with subclonal gain of chromosome 8q22-24 (**figure 5a**). Importantly, MM199 also contained intra-clonal isogenic cells without +8q. To facilitate gene set enrichment analyses (GSEA)^19^ of these sibling cells, with and without +8q, the subclones were next subsampled to obtain subpopulations of cells with matching transcriptome depth (**figure 5b**). This eliminated biases in cell sequencing depths or in transcriptome sizes between the subclones, and thus balanced the likelihood of gene-set detection between subclones, at least in the absence of biological enrichment of specific genes. The gene expression of the matched intra-clonal subpopulations, representing isogenic sibling cells with and without +8q22-24, with equivalent transcriptome sizes, were then compared by GSEA using RTAM2-normalized data. From an analysis of 215 cytoband gene sets (comprising genes defined by chromosome location), +8q cells were strongly enriched for gene-sets located at 8q22-24, with striking statistical confidence (p=0.000, q=0.000, FWER=0.000), compared to cells without +8q (**figure 5d** and **supplemental figure S27)**. In contrast, no other genomic regions were significantly enriched in either subclone. Thus, sciCNV accurately resolved single MM cells into intra-tumor subclones, isolating +8q22-24 as a unique variation distinguishing these.

**Figure 5.**
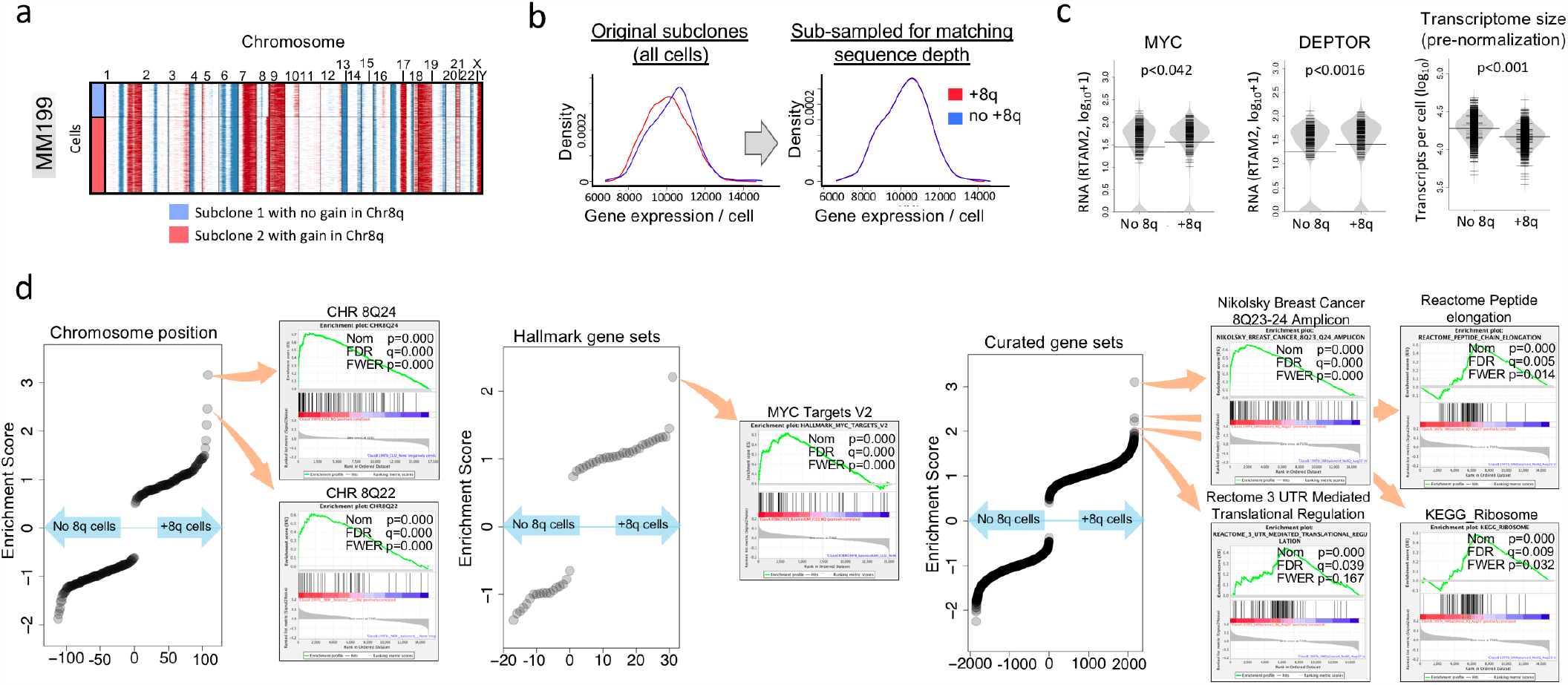
The transcriptional effects of +8q22-24 in MM cells, revealed by dual DNA CNV and RNA profiling of single cells using scRNA-seq and sciCNV. **a.** Heatmaps showing the sciCNV profiles of MM199 cells, showing intra-clonal subpopulations (subclones) that diverge at +8q. **b.** The transcriptome size distribution for cells in each subclones is shown. The original subclones (left) were sampled for subpopulations of cells with comparable transcriptome sizes (right panel), which were then compared in subsequent analyses. **c.** Plots showing the mRNA expression of MYC or DEPTOR genes, located on chromosome 8q24, in MM199 cells by +8q status. Expression data is compressed by normalization (RTAM2) and log scale. The pre-normalized transcriptome sizes of subclonal MM cells from MM199 are also shown, demonstrating fewer RNA transcripts in cells with +8q. P-values were calculated by t-test. **d.** Results of gene set enrichment analysis (GSEA) performed on MM199 intra-clonal subpopulations, comparing cells with or without +8q. GSEA of chromosome position gene sets (n=215) showed enrichment of chromosome 8q22-24 gene sets in cells identified by sciCNV at single cell resolution as containing +8q (left panels). Key results from GSEA of hallmark (n=49) and curated (n=3303) gene sets are shown in the middle and right panels; demonstrating upregulation of MYC target genes in MM199 +8q cells, and enriched expression of the gene sets: Nikolsky_Breast_Cancer_8Q23-24_Amplicon, KEGG_Ribosome, Reactome_Peptide Elongation and Reactome_3’_UTR_mediated translation_regulation.

We next used GSEA to explore the influence of +8q22-24 on cellular programming. Notably, the primary MM cells from MM199 identified by sciCNV as having +8q22-24 showed increased MYC expression (p<0.05) compared to sibling cells without +8q, as expected (**figure 5c**).

Furthermore, the +8q cells also showed broadly increased expression of hallmark MYC target genes (p=0.000, q=0.000, FWER=0.000). Finally, from an analysis of 3303 curated gene sets, primary MM199 cells with +8q22-24 also showed upregulation of gene-sets encoding the machinery of mRNA translation and protein synthesis, including specifically genes involved in 3’UTR-mediated mRNA translation regulation (enrichment rank 5/3303), ribosome biogenesis (rank 4/3303) and peptide chain elongation (rank 3/3303)(**figure 5d, supplemental figure S27**). Conspicuously, these transcriptional effects of +8q22-24 in MM cells were remarkably close to those of +8q23-24 in breast cancer (Nikolsky_Breast_Cancer_8q23-24 amplicon, FWER p=0.000, enrichment rank 1/3303) (**figure 5d, supplemental figure S27**). Thus, overall our analyses indicate that +8q22-24 in primary MM cells induces upregulation of MYC, MYC-target genes and transcriptional programs that promote the upregulation of mRNA translation and peptide synthesis. Analogous gene expression changes appear to occur across malignancies including MM and breast cancer. Strikingly, we were able to identify these transcriptional consequences of +8q22-24 using scRNA-seq analysis of a single MM patient sample. By comparison, the Nikolsky_Breast_Cancer_8q23-24 amplicon signature was derived from the combined genomic analyses of 191 breast cancer samples^20^.

The cellular re-programming induced by +8q22-24 might be expected to promote an increase in gene expression and in cell mass. Notably, however, in the MM sample examined the mTOR-interacting gene, DEPTOR, encoded at 8q24, was also upregulated in +8q cells (**figure 5c**). Upregulation of DEPTOR may limit cell size^21^ and counter any tendency to increased cell mass driven by MYC and ribosome biogenesis. Indeed, from our examination of +8q at a single cell level we observed that the transcriptome sizes of +8q cells were in fact mildly reduced, compared to sibling cells without the CNV (p<0.001) (**figure 5c**). Examination of cellular transcriptome size is a capability unique to scRNA-seq but is generally not feasible from bulk tumor gene expression studies. Thus, from single-cell analyses of +8q22-24 in primary MM cells we find that copy number gain of this region acts to boost MYC, MYC target gene expression and protein synthesis capacity (ribosomes, translation) without increasing cellular transcriptome size. As the transcription of MYC-target genes is relatively increased, this likely leads to enhanced expression of MYC-target genes as proteins. However, the increase in protein synthesis capacity with reduced transcriptome size may also serve to improve the dynamics of protein production by cells and may thus reduce the lag-time between gene expression and proteome change, potentially enhancing cellular adaptability. This represents a unique perspective on the function of +8q22-24 which is offered only by a multi-omic analysis of single cells.

## DISCUSSION

CNVs are critical drivers of cancer biology yet their specific effects on cellular processes often remain only partially characterized. Here, we report bioinformatic methods for the dual profiling of DNA copy number and RNA within the same cells, using scRNA-seq data, which enable direct examination of the effect of intra-clonal CNVs on gene expression. We describe RTAM normalization, which provides enhanced transcriptome alignment to improve gene expression comparisons between cells, and sciCNV, which provides robust RNA-based CNV profiling at single cell resolution. In order to demonstrate and validate these new tools, we examine the transcriptional effects of copy number gain of chromosome region 8q22-24, which represents one of the most common CNV in human cancer. From an examination of intra-clonal subpopulations of cells with and without 8q gain we show using only a single MM patient sample that +8q22-24 induces expression of MYC and MYC target genes, as expected, demonstrating that intra-clonal CNVs can be linked to transcriptional consequences using only scRNA-seq alone. This process can be compared to historical studies of cancer CNVs, in which DNA and RNA profiling of hundreds of tumor samples must be performed to achieve similar results. Moreover, single cell approaches potentially provide higher precision results due to the ability to use intra-clonal isogenic cells as controls. Tumor registry studies, in contrast, are encumbered by the immense genomic heterogeneity the exists between tumors, which persists even when tumors are grouped into cohorts.

Although +8q22-24 upregulates the expression of a broad spectrum of MYC target genes, using single cell profiling we demonstrate that the transcriptomes of MM cells with +8q are in fact smaller than those of cells lacking +8q, at least in the sample examined by us, likely as a result of DEPTOR overexpression from 8q24. At the same time, we find that +8q22-24 cells in primary MM consistently upregulate gene sets involved in mRNA translation, ribosomal biogenesis and peptide elongation. Thus +8q22-24 enhances protein synthesis capacity, without increasing transcriptome size. This likely improves the dynamics of protein expression responses to gene expression; and may enhance the malleability of MM cells with +8q24 to environmental challenges.

Study of the effects of CNVs on cellular programming via single-cell transcriptomics offers a number of advantages. As intra-clonal cells that diverge at a single CNV are virtually isogenic, divergences in their gene expression can be precisely attributed to the CNV. Furthermore, as cells with and without the CNV are derived from the same sample, differences in their gene expression due to the microenvironment, clinical factors or sample processing are minimized. By comparison, tumor cohort studies must instead rely upon the identification of correlations between CNVs and gene expression across unrelated samples and suffer from the substantial confounding genetic and clinical heterogeneity that exists between samples. As a result, the effects of many cancer CNVs on cellular processes remain only partially characterized.

Defining the molecular mechanisms by which oncogenic chromosomal events influence cellular behaviour is a critical step in understanding cancer pathogenesis. Indeed, it is essential knowledge that underlies the development of therapeutic strategies to address the foundations of malignant disease. Auspiciously, single cell multi omics offers the promise of a new era in cancer genomics, in which the multi-faceted effects of CNVs on gene expression and cancer cell programming can now be determined with precision.

## METHODS

### Samples

Bone marrow samples were collected from MM patients in heparinized tubes in compliance with the Declaration of Helsinki and a protocol approved by the University Health Network Research Ethics Board.

### Cell isolation

Marrow cells were separated by Ficoll-Plaque (600g × 30 minutes) and treated with ACK red blood cell lysis buffer (Sigma-Aldrich) for 2 minutes. Cells were then stained with fluorescently-labelled antibodies (Biolegend) targeting CD2 (APC-cy7, RPA-2.10), CD3 (APC-cy7, HIT3a), CD4 (APC-cy7, A161A1), CD11b (PEcy7, CBRM1/5), CD14 (PE-cy7, M5E2), CD34 (PE-Dazzle, 581), CD235a (PE-cy7, H1264), CD38 (APC, HB-7), CD138 (FITC, MI15), BCMA (PE, 19F2,) and CD20 (Pacific blue, 2H7) for 30 minutes in PBS 10% FBS 5mM EDTA at 4°C. B cells and plasma cells were enriched on a MoFlo Astrio (Beckman Coulter) by excluding duplets, non-viable cells (binding 7AAD) and cells expressing non-B lineage markers (CD2, CD3, CD4, CD11b, CD124, CD235a, CD34), and by gating cells with lymphoid FSC-SSC characteristics and expressing CD38, CD138 or CD20, as described^23^ and illustrated in **Supplemental Fig. S29**.

### Single cell RNA-sequencing

Sequencing libraries were prepared from enriched cells using the 10X Genomics Chromium Single Cell Library Kit chemistry v2 and Chromium Single Cell Controller; and were sequenced on an Illumina HiSeq 2500 using paired-ends, targeting >50K mean reads per cell, >80% sequencing saturation and phred-Q30 quality scores >80. Reads were de-multiplexed using Illumina bcl2fastq, mapped to GRCh38 and were then consolidated into unique transcripts per gene per cell using Cell Ranger 2.1.0 (10X Genomics). Poor quality cells with high mitochondrial transcript content were excluded if this represented >5% of total cellular transcripts or was >3 median absolute deviations from the median.

### Non-RTAM scRNA-seq normalization

Normalization and data analysis were performed with R 3.6.1. scRNA-seq expression matrices were normalized using RSEM’s transcript per million (TPM) approach^15^, SCRAN^11^, SCONE^12^ or Suerat’s (v3.1) SCTransform^16^ function, employing *scran, scone* or *seurat V3*.*1* R packages. To facilitate SCONE’s quantile normalization, we limited the input matrix to only those genes that had ≥ 1 read in ≥ 1 cell. When applying Seurat’s SCTransform we used the ‘corrected UMI’ count provided by this function. To compare the output data from each method on similar scales we applied log_2_(.+1) transformation as required.

### RTAM scRNA-seq normalization

#### RTAM1

The normalized gene expression *GE*_*i*,j_ of gene *i* in cell *j* was calculated from the raw transcript count *T*_*i,j*_, which was first normalized to the total transcript count per cell and then scaled to a standard transcriptome library size (10^5^) and log_2_ transformed, mirroring the TPM approach^15^, yielding library-normalized values *L*_*i,j*_:

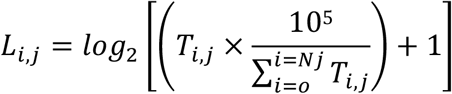

*N*_*j*_ is the number of expressed genes (with ≥ 1 transcript) detected in cell *j*. Genes with non-zero values were then assigned a rank in each cell *R*_*i,j*_ according to descending *L*_*i,j*_. Where multiple genes had the same *L*_*i,j*_ these were assigned an equal rank equivalent to the mean of the values had these genes instead been assigned sequential ranks. *M*_*j*_, the sum of *L*_*i,j*_ of the top-ranked *N*_*T*_ genes was calculated for each cell (from genes satisfying 1 ≤ *R*_*i,j*_ ≤ *N*_*T*_), where *N*_*T*_ was the number of highly-expressed genes used to normalize the dataset.

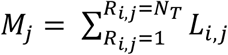

*M*_*k*_, a target constant for correcting *M*_*j*_ values, was calculated as the mean of the initial *M*_*j*_ of all cells. The adjustment variable, *h*_*i,j*_, for gene *i* in cell *j*, was derived such that it adjusts the initial *M*_*j*_ for each cell to the target *M*_*k*_ by differentially adjusting cellular gene expression on a log_2_ scale according to *R*_*i,j*_, and was calculated in the form:

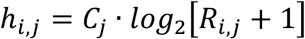

where *C*_*j*_ is a constant for each cell and equals:

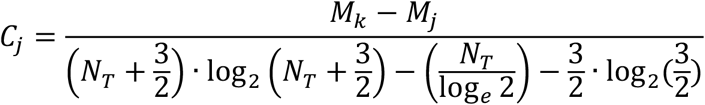

Normalized *GE*_*i,j*_ was calculated as: *GE*_*i,j*_ = *L*_*i,j*_ + *h*_*i,j*_. Or overall,

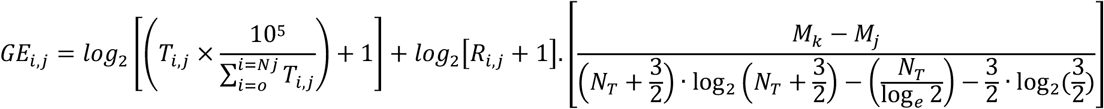

#### RTAM2

*N*_*j*_, *L*_*i,j*_, *R*_*i,j*_, *M*_*j*_ and *M*_*k*_ were calculated as for RTAM1. *G*_*i,j*_ was defined as

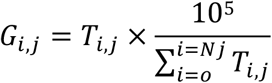

and reflects the transcript count *T*_*i,j*_ of gene *i* in cell j, corrected by scaling to a standard library size (10^5^). The RTAM2 adjustment variable, *h*_*i,j*_, once again was derived to correct cellular *M*_*j*_ to the constant *M*_*k*_ by adjusting gene expression on a non-linear log_2_ scale. However, the RTAM2 adjustment differs from RTAM1 in that the per-gene adjustment is derived from the gene’s expression intensity rather than its expression rank and was calculated as:

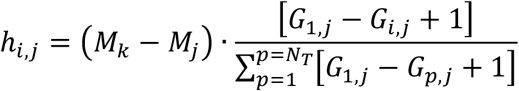

RTAM2 normalized gene expression, *GE*_*i,j*_, was calculated as: *GE*_*i,j*_ = *L*_*i,j*_ + *h*_*i,j*_. Or,

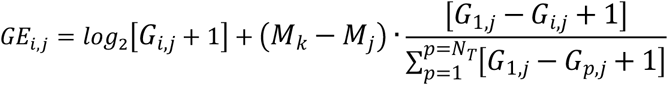

The derivation of RTAM1 and RTAM2 functions are described in the supplemental materials.

#### sciCNV

Single-cell CNV were inferred from the RNA counts of test cells by comparing these normal control cells of the same lineage. In brief, test and control cells were co-normalized using RTAM2. The combined *GE*_*i,j*_ matrix was then truncated to informative genes (*N*_*CNV*_) by excluding genes expressed in <2% cells; and was re-sorted by gene location. *i was* redefined by the rank location of informative genes, numbered from chromosomes 1→Y, p-arm→q-arm, *i* = 1, 2, 3, ⋯, *N*_*CNV*_. To mitigate against data sparsity within individual cells, a moving average of gene expression (ma*GE*_*i*_) was calculated across chromosome segments extending ±*n*/2 genes from gene *i*, where *n* = *N*_*CNV*_/(*S*×50). The variable *S* defined the sharpness (default=1.0) of the analysis resolution and was adjustable to offset sample data sparsity. Data from control cells was merged into a single ‘*average cell*’ with the expression of each gene calculated from its average across all control cells, including zero values, thus integrating the probability of gene detection. A moving average of gene expression across chromosome segments extending ±*n*/2 genes from gene *i* was calculated for the merged control cell, ma*GE*_*i,control*_, mirroring that of test cells. A weighted disparity score, *W*_*i,j*_, comparing gene expression in test cell *j* and the merged control cells was calculated as:

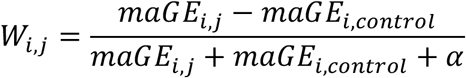

To avoid division by zero, *α* = 0.00001. *W*_*i,j*_ was centred on zero in each cell by subtracting its median, and then linearized by the equation:

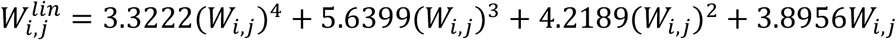

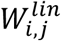 provided a first-pass estimate of single cell CNVs. To further enhance CNV detection, we deployed a second method of assessing gene expression disparity that was sensitive to small concordant expression variations in contiguous genes and insensitive to large single-gene variations. A moving average (*ma*) of gene expression, *U*_*i,j*_, across chromosome segments was calculated for both test and control cells using the *W*_*i,j*_ function above but employing a smaller window extending from *i* ± *n/*(5×*S*) genes. *U*_*i,j*_ was centered on zero in each cell by subtracting its median and then positive/negative values were digitized into +1/−1 values respectively. The digital values were summed cumulatively across the genome as a function of *i*, generating a digital moving score (*DMS)*.

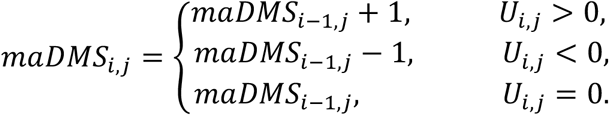

A second digital moving score, *V*_*i,j*_, was calculated as for *U*_*i,j*_ but employed gene expression values without average smoothing in order to obtain enhanced resolution for small CNV. The non-moving average DMS (*nmaDMS*) was then calculated as:

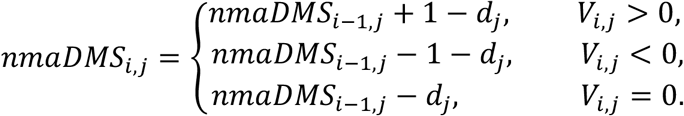

The *nmaDMS* function in each cell *j* was adjusted to sum cumulatively to zero at the last gene (it’s distal terminus) by cumulatively subtracting *d*_*j*_ at each gene, where:

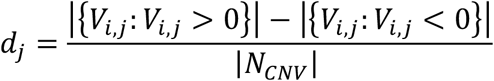

nma*DMS* was scaled to ma*DMS* by the ratio of their respective ranges, (*maDMS*_*MAX*_ *-maDMS*_*MIN*_) / (nma*DMS*_*MAX*_ *– nmaDMS*_*MIN*_). maDMS and nmaDMS were then combined by weighted average to provide the merged function, *DMS*_*i,j*_, integrating both smooth low-resolution and noisy high-resolution data. The gradient of this function, *dDMS*_*i,j*_/*di*, measured over chromosome segments *i* ± *n*/2, provided a second measure of CNV.

The 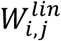 and *dDMS*_*i,j*_ CNV methods were combined by geometric mean. When the two methods yielded conflicting polarity results, *sciCNV*_*i,j*_ → 0. Using the mean *sciCNV*_*i,j*_ result from all clonal test cells a scale factor, *s*, was manually derived to ensure that CNV values optimally centered on integers. For example, for MM samples, *s* was calculated so that clone mean sciCNV for chromosome 13 deletion scaled to −1. *s* was then uniformly applied to all cells:

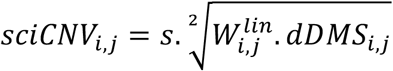

Heatmaps were generated using heatmap.3 R package. Cells were clustered iteratively to identify subclones, using gene expression-based tSNE clustering and sciCNV-based Pearson correlation coefficients. After subclones were identified, supervised clustering was applied. Stochastic noise was reduced by standardizing each sciCNV profile against the median of its nearest neighbours. To further reduce low-level sciCNV noise centered around zero, |*sciCNV*_*i,j*_| values less than threshold, *T*, were set to zero. For tumors in which chromosome gains and losses were substantially imbalanced (with respect to the numbers of genes involved) an optional sciCNV baseline correction, *b*, was applied to correct the zero-set point and improve CNV detection. The method is described further in the **supplemental materials**.

### Whole exome sequencing

DNA was isolated from tumor plasma cells using a Micro kit (Qiagen) and then sheared into 150-200bp fragments using a Covaris E220. Exome libraries were prepared using a Hyper Prep Kit (KAPA Biosystems), pre-capture PCR and SureSelectXT Human All Exon V5+UTR baits (Agilent). Post-capture fragments were amplified by PCR, purified using AMPure-XP beads (Beckman Coulter Genomics) and sequenced using paired-ends on an Illumina HiSeq. FASTQ files were generated with Illumina bcl2fastq and reads were aligned to GRCh38 using BWA v0.7.8^24^ and SAMTOOLS v1.2^22^.

### Correlation of sciCNV and WES-derived CNV

MM199 scRNA-seq data was RTAM2 normalized and sciCNV profiles were calculated (*S*=1.4, *b*=0.1) without a noise filter. MM199 WES was performed using 200ng DNA (>30,000 equivalent exomes) extracted from 1.9×10^6^ pooled cells. WES-derived CNVs were identified using DNAcopy R package v1.54.0. CNV estimates from the two methods were smoothed using windows of identical genomic size and the datasets were paired by nearest genomic location.

### Fluorescence in-situ hybridization (FISH)

Cells on cytospin slides were hybridized with Vysis LSI IGH/CCND1 (14q32:11q13), IGH/FGFR3(14q32:4p16), D5S23, D5S721 SGn/CEP9 SA/CEP15 probes (Abbott Molecular) or Cytocell Aquarius CKS1B/CDKN2C probes according to the manufacturer’s instructions. At least 100 cells were scored for each.

### Tumor CNV score

For each sample, we identified cells with clonal CNVs by sciCNV and using these derived a tumor-specific average sciCNV profile. The sciCNV profiles of all individual cells were then assessed against the tumor average. The tumor score (TS) for cell *j* was summed cumulatively across the genome as:

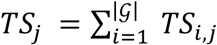

where 𝒢 is the set of all genes in sciCNV matrix and

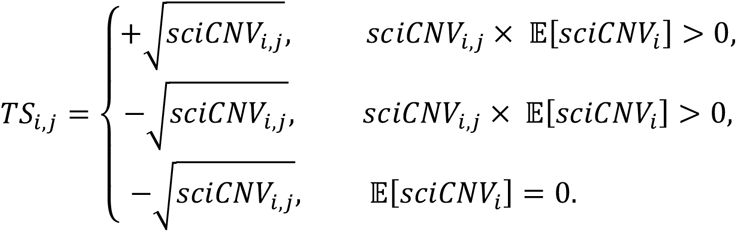

𝔼[*sciCNV*_*i*_] is the average tumor cell *sciCNV*_*i*_ for gene *i*.

### Immunoglobulin isotype score

To evaluate if plasma cells belonged to the tumor clone, we scored their immunoglobulin heavy and light chain (IGH, IGK, IGL) gene expression for restriction to the tumor isotype. Using RTAM2 data, a threshold was identified for IGHM, IGHD, IGHG1/2, IGHA1/2, IGHE and for IGKC and IGL(C2-7) which segregated non-expressing and expressing cells. The immunoglobulin isotype score was calculated for each cell by summing the supra-threshold expression of tumor immunoglobulin genes and subtracting the supra-threshold expression of non-tumor immunoglobulin genes.

### Gene-set enrichment analysis

Intra-clonal cells with divergent CNV were segregated by sciCNV. Paired subpopulations of cells that diverged at a single CNV were then subsampled for cells that were matched for normalized cellular transcriptome size. The matched cells were then compared using gene-set enrichment analysis (GSEA v3.0)^19^ of RTAM2-normalized transcriptomes.

### DATA & CODE AVAILABILITY

The raw and processed scRNA-seq and WES data are deposited at NCBI GEO, GSE141299. The microarray data (Affymetrix U133Plus2.0) for MM tumors with 1q-FISH status was obtained from the NCBI GEO, GSE2658^22^. The code used in this study is available at https://www.github.com/TiedemannLab/sciCNV.

## SUPPLEMENTARY INFORMATION

Supplementary methods, notes and figures can be found on-line.

## ACKNOWLEDGEMENTS

The authors thank the patients and physicians who made this study possible. They also thank N. Winegarden, N. Khuu and G. Basi in the Princess Margaret Genomics Facility and Z. Lu in the Princess Margaret Bioinformatics Core for technical assistance; and Drs. Gary Bader and Caleb Stein for independent review and comments. This work was supported by funding from The Princess Margaret Cancer Centre Foundation, the Terry Fox Foundation and the Canadian Cancer Society Research Institute.

## Author Contribution

A.M-S performed research and analyzed data. N.E., C.L-H. FISH, FACS and whole exome sequencing, respectively. R.E.T. designed research, analyzed data and wrote the paper.

