## Supplemental Methods and Figures for "sciCNV: High-throughput paired profiling of transcriptomes and DNA copy number variations at single cell resolution"

**Supplementary Materials**

1. Online Methods page 3

2. Supplemental References page 25

3. Supplementary Figures page 26

#### Methods

##### Tumor Samples

Human bone marrow samples were obtained from multiple myeloma (MM) patients at The Princess Margaret Cancer Centre at the time of routine clinical biopsy following informed consent. Samples were collected in compliance with the Declaration of Helsinki and a protocol approved by the University Health Network Research Ethics Board. Cells were collected into sterile heparinized tubes.

**Single cell sorting.** Bone marrow cells were diluted 1:1 in phosphate buffered saline (PBS) 10% FBS and separated by Ficoll-Plaque density centrifugation gradient (600g x 30 mintues). The mononuclear fraction was then further processed by exposure to ACK red blood cell lysis buffer (Sigma-Aldrich) for 2 minutes. Cells were washed with PBS 10% FBS and resuspended in PBS 10% FBS 5mM EDTA. An aliquot was retained as an unstained control. The remaining cells were stained for 30 minutes with fluorescently-labelled antibodies targeting CD2 (APC-cy7, clone RPA-2.10, Biolegend), CD3 (APC-cy7, clone HIT3a, Biolegend), CD4 (APC-cy7, clone A161A1, Biolegend), CD11b (PEcy7, clone CBRM1/5 Biolegend), CD14 (PE-cy7, clone M5E2, Biolegend), CD34 (PE-Dazzle, clone 581, Biolegend), CD235a (PE-cy7, clone H1264, Biolegend), CD38 (APC, clone HB-7, Biolegend), CD138 (FITC, clone MI15, Biolegend), BCMA (PE, clone 19F2, Biolegend) and CD20 (Pacific blue, clone 2H7, Biolegend); and were then washed in PBS 10FBS 5mM EDTA. Samples were processed at 4$^{o}$C or on ice. Plasma cells and B cells were enriched on a MoFlo Astrio (Beckman Coulter) using Summit v6.2.7 by gating on cells with lymphoid FSC-SSC characteristics, excluding duplets by FSC-W x FSC-H and SSC-W x SSC-H gates, excluding cells expressing non B cell lineage markers (CD2, CD3, CD4, CD11b, CD124, CD235a) or the hematopoietic stem cell marker CD34, excluding non viable cells (binding 7AAD), and by positive selection for cells expressing CD38, CD138 or CD20, as described[1] with specified modifications. See supplemental figure S28. Cell viability was >95% post FACS by Trypan Blue exclusion.

**Single cell RNA-sequencing**

Sequencing libraries were prepared from FACS-purified plasma cells and B cells (pooled) using the 10X Genomics Chromium Single Cell 3' or 5' Library & Gel bead kit v2 and Chromium Single Cell Controller according to the user guide, targeting 3-6,000 cells per sample. Library constituents incorporated a 26bp read 1, 98bp read 2 and an 8bp sample index and were sequenced on an Illumina HiSeq 2500 instrument using paired-end sequencing, targeting >50K mean reads per cell, >80% sequencing saturation and phred Q30 base quality scores >80%. Raw sequencing reads were de-multiplexed using Illumina bcl2fastq software and then mapped to the GRCh38 human genome reference (using the gene model available at the 10X Genomics online resource), filtered and consolidated into unique mRNA counts per gene per cell using the Cell Ranger 2.1.0 pipeline (10X Genomics). Damaged and non viable cells were identified by their high content of mitochondrial transcripts relative to other mRNA and were excluded from further analyses. Specifically, cells were excluded if the total transcript count from 13 mitochondrial genes represented >5% of total cellular transcripts or if the mitochondrial gene transcript count for the cell was > 3 median absolute deviations from the median for all cells. Secondary data analysis including normalization was performed with R 3.6.1 (as described below).

**Non-RTAM scRNA-seq data normalization**

The raw data obtained from profiling purified B cell lineage cells from primary MM bone marrow samples using the 10X Genomics Chromium scRNA-seq plateform was initially normalized with TPM[2], SCRAN[3], SCONE[4] and Suerat's (v3.1) SCTransform[5] function. TPM normalized values were calculated in R whilst SCRAN, SCONE and SCTransform normalized values were calculated using *scran*, *scone* or *seurat v3.1* R packages. To facilitate SCONE's quantile normalization procedure we limited the input gene expression matrix to include only those genes that had a non-zero read in at least one cell when generating the SconeExperiment object. The Seurat v3.1 SCTransform function is reported to replace earlier NormalizeData, ScaleData and FindVariableFeatures functions and we used the corrected UMI count provided by this new normalization function for the comparison with RTAM. To ensure that the output data from each method was compared on a similar scale we applied $\log_2(x+1)$ transformation to methods that did not natively log transform the expression data.

**RTAM scRNA-seq data normalization**

The following variables were defined:

$T_{i,j}$ = count of transcripts mapped to gene $i$ in cell $j$, with unique molecular identifiers (UMI).

$N_j$ = number of expressed genes (with >= 1 transcript) detected in cell $j$.

$Library_j$ = Library size of cell $j$. $Library_j = \sum_{i=o}^{i=Nj} T_{i,j}$

$G_{i,j}$ = gene expression values for gene $i$ in cell $j$, normalized only by scaling transcript values to a standard library size (e.g. $10^5$).

$$G_{i,j} = T_{i,j} * \frac{10^5}{Library_j}$$

$L_{i,j}$ = $\log_2$-scaled $G_{i,j}$+1, attained after step 1 below.

$R_{i,j}$ = rank of gene $i$ within cell $j$, based on $L_{i,j}$ values ($1^{st}$ rank = highest expression); where genes have identical $L_{i,j}$ they are all assigned an equal midpoint rank.

$N_T$= the number of top-ranked highly expressed genes used to further normalize gene expression intensities per cell. Ideally, $N_T$ is less than the number of genes detected in the smallest (least complex) allowable cell.

$M_j$ = the sum for each cell $j$ of $L_{i,j}$ for top-ranked genes whose $R_{i,j}$ satisfy: $1 \le R_{i,j} \le N_T$. Thus,

$$M_j = \sum_{R_{i,j}=1}^{R_{i,j}=N_T} L_{i,j}$$

$M_k$, a constant, is the target value for normalization of $M_j$ in cell $j$.

$GE_{i,j}$ = final normalized gene expression of gene $i$ within cell $j$.

RTAM1 method:

Normalized expression levels of gene $i$ in cell $j$, $GE_{i,j}$, were derived from raw transcript counts, $T_{i,j}$, normalized as follows:

(1) $T_{i,j}$ were first normalized per cell by library size (the sum of transcript counts) and then scaled to a standard library size ($10^5$) and $\log_2$ transformed, mirroring the transcript per million (TPM) approach utilized by RSEM[2], yielding partially normalized values, $L_{i,j}$:

$$L_{i,j} = log_2\left[\left(T_{i,j} * \frac{10^5}{\sum_{i=o}^{i=Nj} T_{i,j}}\right) + 1\right]$$

(2) Expressed non-zero genes $i$ in cell $j$ were next assigned ranks $R_{i,j}$ = 1,2,.. $N_j$ in order of descending expression, $L_{i,j}$. Where multiple genes had the same transcript count and thus identical $L_{i,j}$ these were assigned an equal rank that represented the midpoint of rank values had these genes instead been individually assigned ranks of sequential integers. An example is shown:

| | | | | | | | | | | |
|---|---|---|---|---|---|---|---|---|---|---|
| $L_{i,j}$ | 10 | 9 | 8 | 7 | 7 | 6 | 5 | 5 | 5 | 5 | 5 |
| $R_{i,j}$ | 1 | 2 | 3 | 4.5 | 4.5 | 6 | 9 | 9 | 9 | 9 | 9 |

(3) $M_j$ was calculated for each cell $j$ as the sum of $L_{i,j}$ of top-ranked genes $i$ whose $R_{i,j}$ satisfy: $1 \leq R_{i,j} \leq N_T$. Thus,

$$M_j = \sum_{R_{i,j}=1}^{R_{i,j}=N_T} L_{i,j}$$

(4) $M_k$, the target value for $M_j$ normalization, was calculated for initial datasets as the average of $M_j$ of all cells, and was later fixed as a constant for subsequent datasets to ensure standardized normalization between data batches, permitting side by side analysis of similarly normalized datasets.

(5) The adjustment variable, $h_{i,j}$, for gene $i$ in cell $j$, was derived such that it normalizes $M_j$ for each cell to the target constant $M_k$ whilst differentially adjusting gene expression on a log2 scale according to $R_{i,j}$, and was calculated as:

$$h_{i,j} = C_j . log_2[R_{i,j} + 1]$$

where $C_j$ is a constant for each cell and equals:

$$C_j = \frac{M_k - M_j}{\left(N_T + \frac{3}{2}\right) . \log_2 \left(N_T + \frac{3}{2}\right) - \left(\frac{N_T}{\log_e 2}\right) - \frac{3}{2} . \log_2 (\frac{3}{2})}$$

(6) Normalized gene expressions, $GE_{i,j}$, were calculated as: $GE_{i,j} = L_{i,j} + h_{i,j}$

Or,

$$GE_{i,j} = log_2\left[\left(T_{i,j} * \frac{10^5}{\sum_{i=o}^{i=Nj} T_{i,j}}\right) + 1\right] + log_2[R_{i,j}+1].\left[\frac{M_k - M_j}{\left(N_T + \frac{3}{2}\right).\log_2\left(N_T + \frac{3}{2}\right) - \left(\frac{N_T}{\log_e 2}\right) - \frac{3}{2}.\log_2(\frac{3}{2})}\right]$$

<u>RTAM2 method:</u>

Steps (1)-(4) were the same as for RTAM1.

(5) The adjustment variable, $h_{i,j}$, for gene (i) in cell (j), was derived in an alternative fashion from RTAM1 whilst still normalizing $M_j$ for each cell to the constant $M_k$ and adjusting gene expression on a non-linear $\log_2$ scale. The RTAM2 adjustment differs from RTAM1 in that the per gene adjustment is calculated from each gene's expression intensity rather than expression rank and requires less assumptions regarding the underlying distribution of gene expression data. The adjustment, $h_{i,j}$, was calculated as:

$$h_{i,j} = (M_k - M_j).\frac{[G_{1,j} - G_{i,j} + 1]}{\sum_{p=1}^{p=N_T}[G_{1,j} - G_{p,j} + 1]}$$

(6) Normalized gene expressions, $GE_{i,j}$, were calculated as: $GE_{i,j} = L_{i,j} + h_{i,j}$

Or,

$$GE_{i,j} = log_2[G_{i,j}+1] + (M_k - M_j).\frac{[G_{1,j} - G_{i,j} + 1]}{\sum_{p=1}^{p=N_T}[G_{1,j} - G_{p,j} + 1]}$$

<u>Rationale for RTAM:</u>

Due to inefficiencies in the processes of RNA capture, reverse transcription (RT) and sequencing quantification of transcripts in any cell, scRNA-seq assays may routinely fail to detect the expression of many lowly expressed genes in some or all cells, and may thus provide variable results between cells. This variability is stochastic in nature and introduces noise into the distribution of RNA transcript data for each cell, particularly amongst lowly expressed genes, some of which may return zero transcript values in some cells despite actual low intensity expression (“drop out”). This stochastic noise within the lower end of the transcript distribution may then be amplified by the fact that the measurement of gene expression in single cells is discretely quantised (raw transcript counts are integer values) and by the effect of log transformation of transcript data, which results in a relative increase in the separation of genes with low expression values (with only 1, 2 or several transcripts detected per cell) compared to genes with higher transcript values, which are instead relatively compressed into small increments that merge into a near continuous spectrum (Fig. 1A-B). This combination of factors serves to increase the quantitative effects of stochastic noise arising from the measurements of lowly expressed genes on global (total) transcript measurements. The issue is further compounded by the high numerical frequency of lowly expressed genes, to the point that the stochastic noise arising in their detection may exert an undue influence on the normalization of the total transcript data generated for cells, potentially outweighing the influences of more reliably detected highly expressed genes.

In contrast, highly expressed genes, with high transcript counts, show more reliable experimental detection across cells and their expression may be more finely resolved relative to their intensity. Thus, expression values for highly expressed genes provide a more robust measure of the cellular transcript distribution than do values from lowly expressed genes and provide a more consistent handle for normalization of cellular data. We therefore sought to enhance cell-specific normalization of scRNA-seq gene expression data utilizing specifically the distribution of top-ranked genes as our guide. Rather than employ a fixed percentile of top expressed genes per cell, which is influenced by cell size and sequencing depth, and by stochastic noise in the detection of lowly expressed genes, we instead normalize transcript distributions using a constant and equivalent number of highly expressed genes per cell, diminishing the influence noisy lowly expressed genes. Transcript distributions are fixed by two points: by their highest expressed gene and by the mean expression of highly expressed genes. Supporting this approach, scaling of TPM-normalized log-transformed scRNA-seq data to achieve uniform mean intensities of highly expressed (top-ranked) genes across cells resulted in an improvement in the uniformity of expression of intermediate intensity housekeeping genes across cells. However, notably when linear scaling of this type was performed on previously log transformed data, the approach, whilst improving the alignment of commonly expressed genes at intermediate expression intensity, also resulted in misalignment of highly expressed common genes. Attempts to achieve a similar normalization goal on log transformed data, via manipulations of pre-log linear data, resulted in insoluble equations (not shown). Instead, we sought to achieve a similar outcome by non-linear log-sensitive differential scaling of gene data that had already been log transformed. Notably, the goal of this normalization approach remained unchanged: to align cell specific transcript measurements on a common scale by standardizing the means of the high-quality data from highly expressed genes across cells (in log-transformed space), irrespective of the identities of those genes within cells.

RTAM Derivation:

The motivation of RTAM1 and RTAM2 normalization is to enhance library-size normalized scRNA-seq data and to align cell-specific sc-RNAseq transcript measurements on a common scale. This is achieved by standardizing the mean intensities of highly-expressed (top-ranked) genes across cells in log-transformed space, irrespective of the identities of those genes within cells. In effect, this resembles normalization of cell-specific data to a standard mean, however in our case lowly expressed genes and zero genes are ignored to avoid their stochastic noise and only data from highly expressed genes, which is more reliably measured, is considered. Furthermore, as transcript counts vary over an exponential scale, the undue influence of one or two very highly expressed genes over that of other intermediate- or highly expressed genes is avoided by utilizing early log transformation. RTAM1 and RTAM2 then seek to achieve a standardized mean intensity of highly expressed genes across cells in log rather than linear space. Finally, as log transformation causes relative contraction of high values and relative expansion of low values, superimposition of a linear adjustment on log-transformed data may cause relatively excessive effect on higher values (which have been contracted by log transformation) and insufficient effect on smaller values (which have been expanded by log transformation) values; therefore RTAM1 and RTAM2 both achieve standardized mean intensities amongst top-ranked genes in log space through a log-sensitive differential scaling of genes according to the expression rank-order (RTAM1) or actual expression intensity (RTAM2) of genes within cells.

RTAM1 derivation:

1. Transcript counts $T_{i,j}$ for genes $i$ in cell $j$ were first normalized per cell by library size (the sum of transcript counts) and were then scaled to a standard library count ($10^5$) and $\log_2$ transformed, mirroring the transcript per million (TPM) approach utilized by RSEM and previously described[2], yielding partially normalized values, $L_{i,j}$:

$$L_{i,j} = log_2\left[\left(T_{i,j} * \frac{10^5}{\sum_{i=o}^{i=Nj} T_{i,j}}\right) + 1\right]$$

(1)

The target library size of $10^5$ transcripts per cell was used as a normalization standard as the 10X Genomics scRNAseq datasets described here typically involved library complexities in the order of ~$10^4$-$10^5$ mapped UMI/cell.

2. As the distribution of gene frequency per transcript count (quantized) tier is approximately exponentially distributed (with the number of genes, and therefore gene ranks, in lower transcript tiers increasing approximately exponentially); and as log transformation of the exponentially-distributed transcript counts yields linearized $L_{i,j}$ values, a plot of $L_{i,j}$ versus gene expression rank, $R_{i,j}$, (with genes ranked in descending order of expression) typically yields an inverted logarithmic curve. This is demonstrated using real world sc-RNAseq data in **supplemental Fig S1**. Consider now a hypothetical cell with a similar distribution of gene rank order versus intensity, as shown in **Supplemental Fig. S2a**: the histogram columns represent the log-transformed expression values $L_{i,j}$ of genes 1,2,3.. $i$.. $N_T$.. $N_j$ (not shown) in cell $j$, in rank-order, $R_{i,j}$, of their expression, where $N_j$ is the number of expressed genes (with transcripts >= 1) detected in the cell and $N_T$ is a constant defining the number of top-ranked genes to be used in the normalization. Each column has a width of 1 rank (1 gene). Irrespective of the actual distribution of $L_{i,j}$ values, the area ($M_j$) under the curve defined by the $L_{i,j}$ values, for cell $j$ from the top-ranked genes 1 through to $N_T$ can be calculated as:

$$M_j = \sum_{R_{i,j}=1}^{R_{i,j}=N_T} L_{i,j}$$

(2)

Division of $M_j$ by $N_T$ yields the average expression of the highly expressed genes in cell $j$.

3. As $N_T$ is constant across all cells, alignment of the mean expression intensities of highly expressed genes across cells requires that cell-specific gene expression values must be scaled such that all $M_j$ are standardized to a constant, $M_k$. To achieve non-linear scaling of genes such that rare highly expressed genes (with log contracted values) are minimally transformed whilst more numerous lowly expressed genes (with expanded values) are transformed to a greater extent, but at a reducing rate reflecting their increasing frequency, a logarithmic function $h$ of gene rank $R_{i,j}$, $h(R_{i,j})$, for cell $j$, is assumed, which is defined by mapping an area ($M_k - M_j$) between gene ranks 1 through to $N_T$. Addition of the adjustments defined by $h(R_{i,j})$ to $L_{i,j}$, results in an area defined by the newly scaled $L_{i,j}$, equal to: $M_j + (M_k - M_j) = M_k$. Therefore application of $h(R_{i,j})$ corrections to $L_{i,j}$ values in cell $j$ will standardize $M_j$ to $M_k$, as desired (**Supplemental Fig. S2b**).

4. $h(R_{i,j})$ is derived as follows. From supplemental Fig. S2c:

$$h(R_{i,j}) = c.\log_2(R_{i,j} + 1) \tag{3}$$

The constant $c$, can be defined at $R_{i,j} = R_{N_T,j}$ (at the rank of the $N_T$-th gene) as:

$$c = \frac{h(R_{N_T,j})}{\log_2(R_{N_T,j} + 1)} \tag{4}$$

Combining (3) and (4):

$$h(R_{i,j}) = h(R_{N_T,j}).\frac{\log_2(R_{i,j} + 1)}{\log_2(R_{N_T,j} + 1)} \tag{5}$$

From **supplemental Fig. S2b-c**, the adjustment area defined by $h(R_{i,j})$ is equal to:

$$M_k - M_j = \int_{\frac{1}{2}}^{R_{N_T,j}+\frac{1}{2}} h(R_{i,j}).dR = \int_{\frac{1}{2}}^{R_{N_T,j}+\frac{1}{2}} h(R_{N_T,j}).\frac{\log_2(R_{i,j} + 1)}{\log_2(R_{N_T,j} + 1)}.dR \tag{6}$$

$$= \frac{h(R_{N_T,j})}{\log_2(R_{N_T,j}+1)}.\left[\left(R_{N_T,j}+\frac{3}{2}\right).\log_2\left(R_{N_T,j}+\frac{3}{2}\right)-\frac{R_{N_T,j}}{\log_e(2)}-\frac{3}{2}.\log_2(\frac{3}{2})\right]$$

(7)

Thus, from equations (6) and (7), $h(R_{N_T,j})$ can be defined in terms of $M_k - M_j$, and constants derived from $R_{N_T,j}$. From equation (5), $h(R_{i,j})$ can be defined in terms of $h(R_{N_T,j}), R_{i,j}$ and constants. As $R_{N_T,j} = N_T, j$ and is identical for all cells, substitution of $N_T$ and rearrangement of (5)-(7) yields:

$$h(R_{i,j}) = C_j.\log_2(R_{i,j}+1)$$

(8)

where $C_j$ is a constant for cell $j$, equal to:

$$C_j = \frac{M_k - M_j}{\left(N_T+\frac{3}{2}\right).\log_2\left(N_T+\frac{3}{2}\right)-\frac{N_T}{\log_e(2)}-\frac{3}{2}.\log_2(\frac{3}{2})}$$

(9)

5. The adjusted gene expression, $GE_{i,j}$, of non-zero genes $i$ in cell $j$ is calculated as:

$$GE_{i,j} = L_{i,j} + h(R_{i,j})$$

(10)

Or,

$$GE_{i,j} = log_2\left[\left(T_{i,j} * \frac{10^5}{\sum_{i=o}^{i=Nj} T_{i,j}}\right)+1\right] + log_2[R_{i,j}+1].\left[\frac{M_k - M_j}{\left(N_T+\frac{3}{2}\right).\log_2\left(N_T+\frac{3}{2}\right)-\left(\frac{N_T}{\log_e 2}\right)-\frac{3}{2}.\log_2(\frac{3}{2})}\right]$$

(11)

RTAM2 derivation:

1. The initial derivation of RTAM2 mirrors RTAM1. Derivation point 1 is identical. Expressed (non-zero) genes are again ranked, $R_{i,j,}$ for $i$=1,2,.. $N_j$ in order of descending relative expression ($G_{i,j}$ or $L_{i,j}$) using the same ranking rules described above.

2. The goal of RTAM2, like RTAM1, is to achieve alignment across cells of the cell-specific mean expression value of the $N_T$ top-expressed genes within each cell. In applying the method, the sum of the log-transformed expression values $L_{i,j,}$ of the $N_T$ top expressed genes for each cell $j$ ($M_j$) is aligned to $M_k$, a standardized constant across cells (typically equal to the mean of $M_j$). Thus:

$$M_j = \sum_{R_{i,j}=1}^{R_{i,j}=N_T} L_{i,j} \tag{2}$$

   Division of $M_j$ by $N_T$ yields the average expression of the highly expressed genes in cell $j$. As $N_T$ is identical for all cells, alignment of $M_j$ with $M_k$ serves to align the cell-specific mean expression value of the $N_T$ top-expressed genes within each cell.

3. Alignment of $M_j$ with $M_k$ across cells is produced by non-linear scaling of cell-specific gene expression values by addition of a variable adjustment value, $h_{i,j,}$ to the log-scale data, $L_{i,j}$. As the log scaled expression data of highly expressed genes (which are more precisely measured) is relatively contracted whereas the expression intensities of lowly expressed genes is relatively expanded by log transformation (despite reduced precision in their measurement), a correction function $h_{i,j}$ for each cell $j$ is assumed which is a linear function of gene intensity, $G_i$, thus $h(G_{i,j})$. **Supplemental Figure S3** shows the relationship between $G_i$, $L_i$ and $h_i$ in cell $j$. $h(G_{i,j})$ is defined as:

$$h(G_{i,j}) = c_j.(G_{1,j} - G_{i,j} + 1) \tag{12}$$

   To achieve the goal of the RTAM2 normalization, addition of the adjustments $h(G_{i,j})$ to $L_{i,j}$ in cell $j$ must cumulatively result in a correction that alters $M_j$ to the target $M_k$. As the correction is equivalent to $M_k - M_j$:

$$M_k - M_j = \sum_{i=1}^{i=N_T} h\left(G_{i,j}\right) \tag{13}$$

4. From equation 12, at the gene with expression rank $N_T$,

$$h\left(G_{N_T,j}\right) = c_j.\left(G_{1,j} - G_{N_T,j} + 1\right)$$

Therefore:

$$c_j = \frac{h\left(G_{N_T,j}\right)}{\left(G_{1,j} - G_{N_T,j} + 1\right)} \tag{14}$$

5. Combining equations 12 and 14:

$$h\left(G_{i,j}\right) = h\left(G_{N_T,j}\right).\frac{\left(G_{1,j} - G_{i,j} + 1\right)}{\left(G_{1,j} - G_{N_T,j} + 1\right)} \tag{15}$$

6. Combining equations 13 and 15:

$$M_k - M_j = h\left(G_{N_T,j}\right).\sum_{i=1}^{i=N_T} \frac{\left(G_{1,j} - G_{i,j} + 1\right)}{\left(G_{1,j} - G_{N_T,j} + 1\right)} \tag{16}$$

Therefore:

$$h\left(G_{N_T,j}\right) = \frac{M_k - M_j}{\left[\sum_{i=1}^{i=N_T} \frac{\left(G_{1,j} - G_{i,j} + 1\right)}{\left(G_{1,j} - G_{N_T,j} + 1\right)}\right]} \tag{17}$$

7. Combining equations 15 and 17, and substituting the summation function variable *p* in place of *i* in the summation operator, to distinguish it from *i* outside of the operator:

$$h(G_{i,j}) = (M_k - M_j).\frac{[G_{1,j} - G_{i,j} + 1]}{\sum_{p=1}^{p=N_T}[G_{1,j} - G_{p,j} + 1]}$$

(18)

8. The adjusted gene expression, $GE_{i,j}$, of non-zero genes $i$ in cell $j$ is calculated as:

$$GE_{i,j} = L_{i,j} + h(G_{i,j})$$

(19)

Or,

$$GE_{i,j} = log_2[G_{i,j} + 1] + (M_k - M_j).\frac{[G_{1,j} - G_{i,j} + 1]}{\sum_{p=1}^{p=N_T}[G_{1,j} - G_{p,j} + 1]}$$

(20)

RTAM2 limits:

To avoid overcorrection of data and output of negative values or non-sensible re-ordered results which might otherwise occur with extreme instances of $M_k - M_j$ disparity and unusual $G_{i,j}$ intensity distributions the following limits were developed from the above derivation:

9. Limit 1. If:

$$h(G_{N_T,j}) \geq \frac{(G_{1,j} - G_{N_T,j} + 1)}{(G_{1,j} + 1).\ln(2)}$$

(21)

then the $h(G_{i,j})$ adjustment factors for cell $j$ are corrected by multiplying by the limit factor, $Lim1_j$, prior to addition to $L_{i,j}$, where $Lim1_j$ is calculated as:

$$Lim1_j = \frac{(G_{1,j} - G_{N_T,j} + 1)}{(G_{1,j} + 1).\ln(2).h(G_{N_T,j})}$$

(22)

10. Limit 2. For cell $j$, let $min_j$ correspond to the $i$ value for the lowest expressed genes (lowest gene expression tier). If:

$$-h\left(G_{min_j,j}\right) \geq \log_2\left(G_{min_j,j}+1\right) \tag{23}$$

then the $h(G_{i,j})$ adjustment factors for cell j are corrected by multiplying by the limit factor, $Lim2_j$, calculated as:

$$Lim2_j = \frac{\log_2\left(G_{min_j,j}+1\right)}{-h\left(G_{min_j,j}\right)} \tag{22}$$

In actual practice, these limits were never applied for any cell in any of the datasets examined.

**Single cell inferred copy number variation (sciCNV) analysis.**

DNA copy number variations in single cells were inferred from mRNA counts by comparing the gene expression profiles of test cells with that of normal diploid control cells of the same or similar lineage. Test and control cell data was co-normalized using RTAM-1 or -2. The normalized gene expression data from test and control cells ($GE_{i,j}$ matrix) was first condensed to informative genes (approximately 10,000 genes in each of our samples, $=N_{CNV}$) by excluding loci expressed in < 2% cells; and was then sorted by chromosomal location, with $i$ redefined by genomic location rank, $i$= 1,2,3... $N_{CNV}$. The sharpness (resolution) of the CNV analysis was defined by the variable *sharpness*, with default value of 1.0; and was adjusted in the range 0.6-1.4 to offset sample data sparsity: higher values permit sharper detection of small CNV but require greater data density for accuracy. Variable $n$ was calculated as $N_{CNV}/(50*sharpness)$. To help mitigate against data sparsity within individual test cells, a moving average of gene expression (ma$GE_i$) was developed across each chromosome in segments extending $-n/2$ genes retrograde (towards p-termini) and $+n/2$ genes anterograde (towards q termini) of gene $i$. For genes located near chromosome termini the moving average window boundary was reduced from $n/2$ genes to the maximum allowed by the terminus, but maintained at $n/2$ on the other side. In contrast to test cells, data sparsity in control cells was first addressed by merging the data from all control cells into a single “ideal” control cell. In this “ideal control cell” the expression of each gene was calculated as its average expression across all control cells, including zero values from cells in which the gene was not detected. Thus, in the merged control cell the expression values for genes that were not uniformly detected but which were detected in only a fraction, $F_c$, of control cells (the majority of detected genes) were reduced from their average expression level in the cells in which the gene was detected, to a lower value that was reflective of the factor $F_c$. The inclusion of this detection probability factor on average gene expression intensity in the idealized control cell permits a fair comparison of stochastically detected gene expression data from single test cells (which suffer high frequencies of zero counts for lowly expressed genes) with the control cell data (in which the probabilistic rate across cells for gene drop-out is factored into the reference gene expression value), particularly when comparisons are made across a large number of genes. To achieve a fair comparison across multiple genes a moving average of gene expression across chromosome segments was calculated for the merged control cell, ma$GE_{control}$, mirroring the approach used for test cells. A weighted disparity score, $W_{i,j}$, comparing the gene expression in the chromosome segment centred on gene $i$, in test cell $j$, with the expression from the same region in the merged control cell was calculated across all genes as:

$$W_{i,j} = \frac{maGE_{i,j} - maGE_{i,control}}{maGE_{i,j} + maGE_{i,control} + \alpha}$$

The difference between test and merged control cell gene expression across chromosome segments normalized by the sum of gene expression to diminish the influence of baseline gene expression patterns on CNV-driven changes in gene expression. No prior median normalization of data had been invoked. To avoid division by zero in instances where no gene expression was detected across chromosome segments in either test or control cells, $\alpha = 0.00001$. As $W_{i,j}$ is non-linear in reporting copy number changes relative to gene expression changes and may be influenced by variations in data distribution between cells, $W_{i,j}$ was next centred to zero by subtracting its median, and was then linearized by the quadratic equation:

$$W_{i,j}lin = 3.3222(W_{i,j})^4 + 5.6399(W_{i,j})^3 + 4.2189(W_{i,j})^2 + 3.8956W_{i,j}$$

The solution for $W_{i,j}$ linearization is derived from the observation that $W_{i,j}$ plotted against theoretical copy number changes yields the relationship y = -0.0035x$^4$ + 0.0224x$^3$ - 0.0694x$^2$ + 0.244x + 0.0025 when $\alpha$ is 0.00001 with $R^2$=0.99997. Following scaling, $W_{i,j}lin$ provides a first pass inference of CNV in single cells.

To further enhance the detection of CNV in single cells, we complemented the $W_{i,j}lin$ function with a second method of assessing changes in gene expression within chromosomal segments that is insensitive to quantitative changes in expression values but sensitive to coherent qualitative changes in the expression of multiple contiguous genes. To best capture major trends a moving average of gene expression across chromosome segments was first developed for both test cells and the idealized control cell, as above, however using a smaller moving window extending from *i–n/a* through *i+n/a* genes, where *a*=5**sharpness*. The relative expressions of genes in test and control cells within sequential chromosome segments were compared using the $W_{i,j}$ calculation above, with these results (from the smaller moving average window) defined as $U_{i,j}$ and centered on zero by subtracting the median. The $U_{i,j}$ values were then exchanged for a 'vote' of +1 or -1 at each locus according to whether expression was greater or lower in the test cell versus the merged control cell. The +1 or -1 values were next summed cumulatively from the p terminus of chromosome 1 (gene 1) to the q terminus of the X or Y chromosome (final $N_{CNV}$ gene), generating a democratic moving score (*ma-DMS)*. At the first locus (at the p-terminus of chromosome 1) *ma-DMS* → 0. For successive chromosome segments centered on gene *i*, the *ma-DMS* was cumulatively adjusted:

if $U_{i,j} > 0 \rightarrow \text{ma-}DMS_{i,j} = \text{ma-}DMS_{i-1,j} + 1$
if $U_{i,j} < 0 \rightarrow \text{ma-}DMS_{i,j} = \text{ma-}DMS_{i-1,j} - 1$
if $U_{i,j} = 0 \rightarrow \text{ma-}DMS_{i,j} = \text{ma-}DMS_{i-1,j}$

To improve the resolution for small copy number gains or deletions a second democratic moving score was calculated in parallel, based on gene expression values that were not subjected to smoothing by moving average. The relative expression of each individual gene *i* in the test cell and merged control cell were compared using the $W_{i,j}$ calculation above, with results defined as $V_{i,j}$. The non moving average DMS, *nma-DMS*, was calculated as:

if $V_{i,j}$ >0 → nma-$DMS_{i,j}$ = nma-$DMS_{i-1,j}$ + 1 - $d_j$

if $V_{i,j}$ <0 → nma-$DMS_{i,j}$ = nma-$DMS_{i-1,j}$ - 1 - $d_j$

if $V_{i,j}$ =0 → nma-$DMS_{i,j}$ = nma-$DMS_{i-1,j}$ - $d_j$

The *nma-DMS* function in cell *j* was adjusted to sum to zero at the last gene (ie. at it's distal terminus) by cumulatively subtracting $d_j$, where $d_j$ was calculated as:

$$d_j = \frac{count(V_{i,j}\ > 0) - count(V_{i,j}\ < 0)}{N_{CNV}}$$

The *nma-DMS* function was next scaled to the ma-DMS function by multiplying it by the ratio of their respective ranges (*ma-DMS*$_{MAX}$ - *ma-DMS*$_{MIN}$) / (nma-*DMS*$_{MAX}$ – *nma-DMS*$_{MIN}$). The ma-DMS and nma-DMS were then combined in a 2:1 weighted average to provide the merged function, $DMS_{i,j}$, consisting of both smooth low resolution data plus noisy high resolution data.

The gradient of the merged function, $dDMS_{i,j}$, plotted against *i* (gene location rank) provided a second measure of copy number change. $dDMS_{i,j}$ was measured from $DMS_{i,j}$ over each chromosome segment *i*–*n*/2 to *i*+*n*/2 for each cell *j*.

The two CNV methods were combined by their geometric mean. In regions where the two methods yielded conflicting results of opposite polarity (gain versus loss), $sciCNV_{i,j}$ was set to zero. Using the average of $sciCNV_{i,j}$ from all test cells a scale factor, *s*, was manually derived to ensure CNV values optimally centered on integers (for example, *s* was often defined for MM samples so that chromosome 13 deletion scaled to -1) and was then uniformly applied:

$$sciCNV_{i,j} = s.\sqrt[2]{W_{i,j} lin.dDMS_{i,j}}$$

Heatmaps were generated using R package heatmap.3. Cell (rows) were clustered initially by either t-distributed Stochastic Neighbor Embedding (tSNE)-based clustering of gene expression which, in some instances, helped reveal subclones, or were ordered initially by Pearson correlation coefficients between sciCNV profiles to reveal subpopulations with similar CNV. As noise was a potential confounder of automated clustering, supervised clustering was then applied to better isolate specific CNVs and subclones. After the final clustering of cells into subclones, sciCNV noise was reduced by standardizing each profile against the median of its nearest neighbours. This had the effect of reducing stoichastic noise and improving signal:noise ratio. For CNV heatmap representations $|sciCNV_{i,j}|$ values less than a noise threshold of approximately 0.5 were set to zero.

As real world copy number changes in single cells are integers, $sciCNV_{i,j}$ where $z$-0.5 < $sciCNV_{i,j}$< $z$+0.5, $\{z \in Z(integers)\}$ can be optionally adjusted to converge on the closest integer:

$$sciCNV_{adj} = z\sqrt{\frac{sciCN_{orig}}{z}}$$

sciCNV baseline correction: For the default sciCNV analysis an assumption is made that chromosome gains and losses are balanced (with respect to the numbers of genes involved in each) and that the median CNV result for genes across the genome in each cell is zero. When these assumptions are substantially invalid (for example, as in the case of a hyperdiploid MM karyotype), a correction can be applied to improve CNV detection and correct the CNV zero set point. Let $b$ represents the net fraction of $N_{CNV}$ genes expected to be gained or lost in excess of an equal balance. If a cell or cells is hyperdiploid with a large fraction of genes duplicated by trisomies with no corresponding chromosome losses, then the default sciCNV analysis (with b=0) yielded evidence of the trisomies but as a result of median centering the data the initial analysis diminishes the intensity of sciCNV gains and offsets these gains by inferring complementary regions of loss across the non-trisomic chromosomes. Notably, erroneous median centering is a potential limitation of any copy number estimation approach that is based upon comparing regional intensities. The assumption of balanced gains and losses can be altered in the sciCNV algorithm by adjusting $b$ to a fraction representing the approximate net genomic change (using a positive fraction for gain and a negative fraction for net genomic loss). The net number of genes that should be detected as gained is $b^* N_{CNV}$; when b is non-zero the assumption that the cumulative DMS functions should terminate with gene $N_{CNV}=0$ is invalidated and the cumulative DMS is corrected to terminate at $b^* N_{CNV}$ by cumulatively adding the adjustment $+b$ over $N_{CNV}$ genes. Like the DMS functions, $W_{i,j}$ is normalized by default to it's median value, which is assumed to represent zero CNV; however when $b$ is set to a non zero value $W_{i,j}$ is normalizing to a non-50$^{th}$ percentile value (0.5+$b$/2), shifting the $W_{i,j}$ zero point and correcting the corresponding CNV estimates.

**Correlation of scRNA-seq and WES derived CNV**

Single cell inferred CNV data was compared with WES-derived copy number measurements using a patient bone marrow sample for which matching tumor bulk WES and scRNA-seq data were available. scRNA-seq data was normalized using RTAM2 and sciCNV results were calculated as described above with settings: *sharpness*=1.4 (representing a window size of 164 genes) and $b$=0.1. The sciCNV output was analysed without a noise threshold filter or integer convergence algorithm. The exome library was prepared from 200ng DNA (corresponding to >30,000 average exomes) derived from $1.9 \times 10^6$ pooled cells as described earlier. DNA WES-based CNVs were calculated using DNAcopy R package version 1.54.0. Smoothed CNV estimates from the two methods were derived using a moving average window of identical genomic size (corresponding to 164 genes in the scRNA analysis) and the datasets were paired by nearest genomic location, measured from the start of genes. Locus-specific CNV values from the two methods were compared by XY plot and regression analysis.

**Single cell tumor CNV score**

Copy number analysis of MM sample scRNAseq data alongside controls readily revealed tumor-specific sciCNV curves that were highly correlated between tumor cells. For each tumor we segregated a population of cells showing tumor CNVs and derived a tumor-specific average CNV curve from sciCNV data. The sciCNV curves of all cells were then assessed against the tumor-specific CNV curve to assess their likeness to the tumor karyotype. The tumor score (TS) for cell j was summed cumulatively across the genome as:

$$TS_j = \sum_{i=1}^{all\ genes} TS_{i,j}$$

where $TS_{i,j} = \begin{cases} +\sqrt{sciCNV_{i,j}}, & if\ sciCNV_{i,j} * average\ tumor\ cell\ sciCNV_i > 0 \\ -\sqrt{sciCNV_{i,j}}, & if sciCNV_{i,j} * average\ tumor\ cell\ sciCNV_i < 0 \\ -\sqrt{sciCNV_{i,j}}, & if average\ tumor\ cell\ sciCNV_i = 0 \end{cases}$

**Single cell tumor immunoglobulin (Ig) isotype score**

To evaluate if plasma cells or B cells potentially belonged to the tumor clone we assessed immunoglobulin gene expression in single cells and scored this for restriction to the tumor Ig isotype. Immunoglobulin heavy chain (IGH) and light chain (IGK,IGL) gene expression was assessed using RTAM2 normalized scRNA-seq. From a review of the data for all cells a minimum threshold value was determined for each IGH (IGHM, IGHD, IGHG1/2, IGHA1/2, IGHE) and for IGKC and IGL(C2-7) which segregated non-expressing cells from expressing cells. The tumor Ig-restriction score was then derived for each cell by adding expression values for the Ig genes expressed by the tumor (after deducting the expression threshold for these genes) and subtracting expression values for all other Ig genes not expressed by the tumor (after deducting the background expression threshold for these genes).

**In silico separation of normal plasma cells (NPC) and multiple myeloma plasma cells (MMPC) from other cells.**

Sequenced cells were initially categorized by gene-expression based tSNE clustering. Plasma cell populations were identified by the expression of canonical markers including SDC1, SLAMF7 and TNFRSF17. NPC and MMPC formed distinct clusters. NPC expressed polyclonal immunoglobulin while MMPC expressed immunoglobulin of the tumor clone isotype or no detectable immunoglobulin. To avoid contamination of control NPC by MMPC we excluded rare cells expressing the tumor immunoglobulin isotype.

**Tumor bulk whole exome sequencing**

Tumor bulk DNA was isolated from FACS-purified plasma cells using a Qiagen Micro kit and quantified using a Qubit 3.0 (Life Technologies); 200ng DNA was then sheared into 150-200bp fragments using a Covaris E220. Exome libraries were constructed using a Hyper Prep Kit (KAPA Biosystems), pre-capture PCR amplifcation and SureSelectXT Human All Exon V5+UTR baits (Agilent). Post capture fragments were further amplified by PCR, purified using

AMPure XP beads (Beckman Coulter Genomics) and analyzed by Agilent 2100 Bioanalyzer. Paired-end sequencing was performed on an Illumina HiSeq to an average depth of approximately 150x. FASTQ files were generated with Illumina bcl2fastq and reads were aligned to GRCh38 using BWA v 0.7.8[6] and SAMTOOLS v1.2[7].

**Fluorescence in situ Hybridization (FISH)**

Cells adherent to cytospin slides were fixed in ice-cold 3:1 methanol/acetic acid, incubated in 2X SSC for 30 min at 37$^{o}$C, dehydrated in a series of ethanol washes and hybridized with Vysis LSI (Abbott Molecular) or Cytocell Aquarius DNA Probes according to the manufacturer's instructions. The probe sets included Vysis LSI IGH/CCND1 (14q32:11q13) and LSI IGH/FGFR3(14q32:4p16) Dual Color translocation probes, Vysis LSI D5S23, D5S721 SGn/CEP 9 SA/CEP 15 SO Probes and Cytocell Aquarius CKS1B/CDKN2C(p18) Amplification/Deletion Probes (1q21/1p32.3). At least 100 cells from each slide were scored. The background rate for each set of FISH probes was determined by counting >200 cells from pooled bone marrow from 5 healthy male donors.

**Gene set enrichment analysis**

Cells belonging to genetically-divergent intraclonal populations within single tumor samples were identified and segregated using single cell sciCNV profiles; these populations were then subsampled to achieve paired subpopulations that were matched in distribution density for total summed gene expression per cell (normalized cellular transcriptome size). The gene expression activities of cells within the standardized subpopulations were then compared using gene set enrichment analysis (GSEA v3.0)[8] of RTAM2-normalized transcriptome data. GSEA of intra-clonal subpopulations was performed against chromosome position (cytogenetic band) gene sets, hallmark gene sets and curated pathway database gene sets. To understand similarities and dissimilarities between these intra-tumor single cell analysis and classical inter-tumor bulk sample studies, microarray gene expression data (Affymetrix U133Plus2.0) for bulk tumor samples from 532 MM patients, annotated with FISH 1q status, was obtained from the NIH Gene Expression Omnibus, GSE2658[9] and similar GSEA analyses were performed using tumors segregated by the same CNV feature (for example absence or presence of +1q).

**Data & Code availability**

The raw and processed single cell RNA-seq and whole exome sequencing data are deposited at NCBI GEO, accession number GSE141299. The tumor bulk microarray gene expression data analyzed in this study is available at GSE2658. The code used in this study is available at https://www.github.com/TiedemannLab/sciCNV.

#### Supplementary Figures

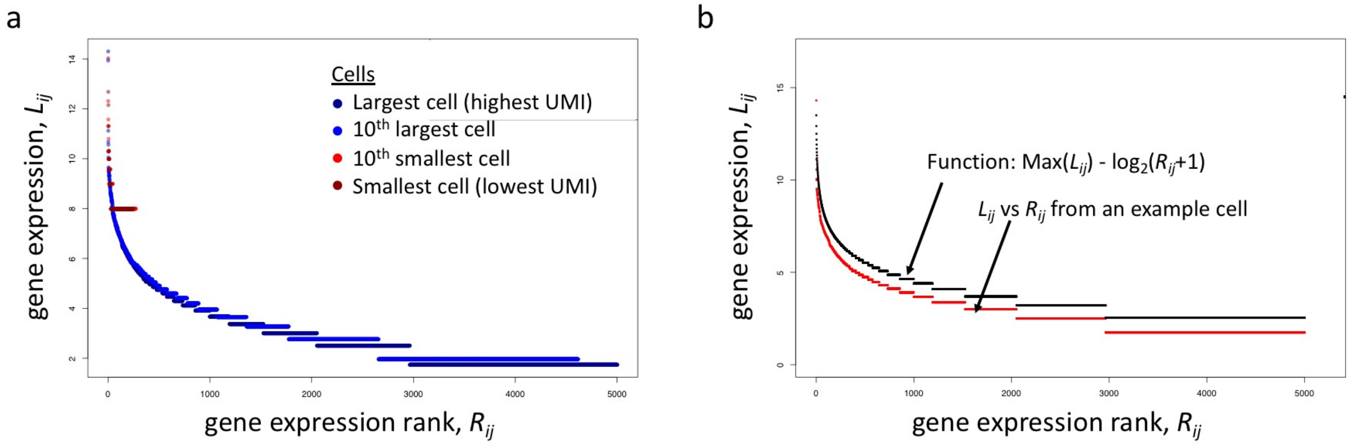


**Figure S1. Semi-standardized gene expression ($L_{ij}$) versus expression rank ($R_{ij}$) within single cells.**

(See the *RTAM Derivation* section in the methods).

**a.** The partially-normalized gene expression values, $L_{i,j}$, for gene $i$ in cell $j$ are plotted against the corresponding rank of each gene in each cell (defined by its expression intensity), $R_{ij}$, for four single cells of varying sizes (large and small), using real world data. In order to calculate $L_{i,j}$ the raw transcript counts for gene $i$ in cell $j$ were first partially normalized by adjustment of cell $j$'s total transcript count (and constituent gene transcript counts) to a standard library size, as described in the methods, yielding a semi-standardized estimate of gene expression $G_{i,j}$. $L_{i,j}$ is the $\log_2$(+1)-transformation of $G_{i,j}$. $L_{i,j}$ gene expression tiers with multiple genes are shown as horizontal bars with a midpoint at $R_{ij}$; the bars derive from the fact that initial scRNA-seq transcript counts are whole numbers. The length of each bar represents the number of genes an identical transcript count. The bars (or transcript count tiers) are shorter for highly-expressed genes, where the curve of $L_{i,j}$ vs $R_{i,j}$ is more tightly defined for each cell. For small (or shallowly sequenced) cells, with low library size, genes with low transcript counts and broader gene expression tiers occur at lower $R_{i,j}$. UMI, unique molecular identifiers (representing total unique transcripts or library size per cell).

**b.** The plot of $L_{i,j}$ versus $R_{i,j}$ data, for a single representative cell $j$, is compared with the mathematical log function, Max($L_{i,j}$)-log2($R_{i,j}$+1), showing an initial close correlation, which can readily be further aligned.

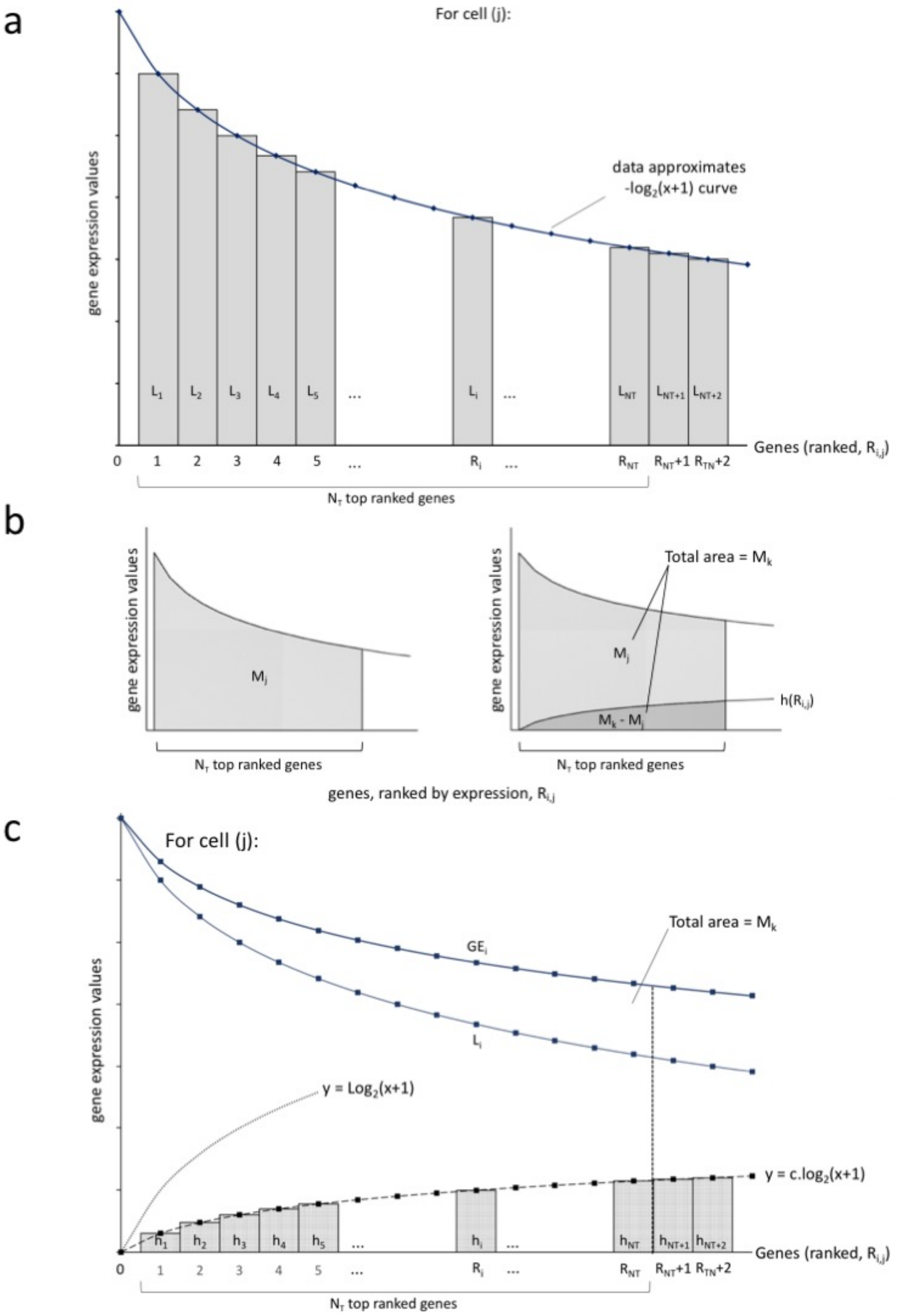


**Figure S2. RTAM1 derivation**

(See the *RTAM Derivation* section in the methods).

**a.** The raw transcript counts per gene for cell (j) are partially normalized by adjustment of the cellular total transcript count (and constituent gene transcript counts) to a standard library size, as described in the methods. The schema above then shows $L_{i,j,}$ (dark blue line with diamonds), the $\log_2(+1)$-function of the partially normalized linear gene expression, $G_{i,j}$ (for genes 1,2,3..*i* in the single cell *j*) plotted against the gene expression rank, $R_{i,j}$ of each gene in cell (j) (on x-axis). The top-ranked $N_T$ genes in cell (j) are demonstrated. Each gene represents a single $R_i$ rank. For RTAM1 the area under $L_{i,j,}$ for the top ranked 1,2..$N_T$ genes ($M_j$), representing the sum of expression in log-space of top-ranked genes, can be approximated by summing successive columns of area= $L_i$ (height $L_i$, width=1) for the top-expressed genes *i*=1 through to $N_T$.

**b.** The RTAM1 adjustment function, h($R_{i,j}$), for gene *i* in cell *j* is shown and by design acts to adjust the log-transformed gene expression using an inverted log-based correction. The function defines a correction area, $M_k$-$M_j$, between top-ranked genes 1 though $N_T$ in cell j. Addition of h($R_{i,j}$) to $L_{i,j,}$ converts the initial area $M_j$ (the sum of $L_{i,j}$ for genes *i*=1 through $N_T$ for cell *j*), which is defined both by $L_{ij}$ and gene expression rank $R_{ij}$ (left panel), to $M_k$, a constant across cells.

**c.** Schema illustrating the derivation of $h(R_{i,j})$, a log2 function of $R_{ij}$+1, which sweeps out an area of $M_k$-$M_j$, between the top-ranked genes 1 though $N_T$ in cell j. A full description of the derivation of $h(R_{i,j})$ is included in the methods.

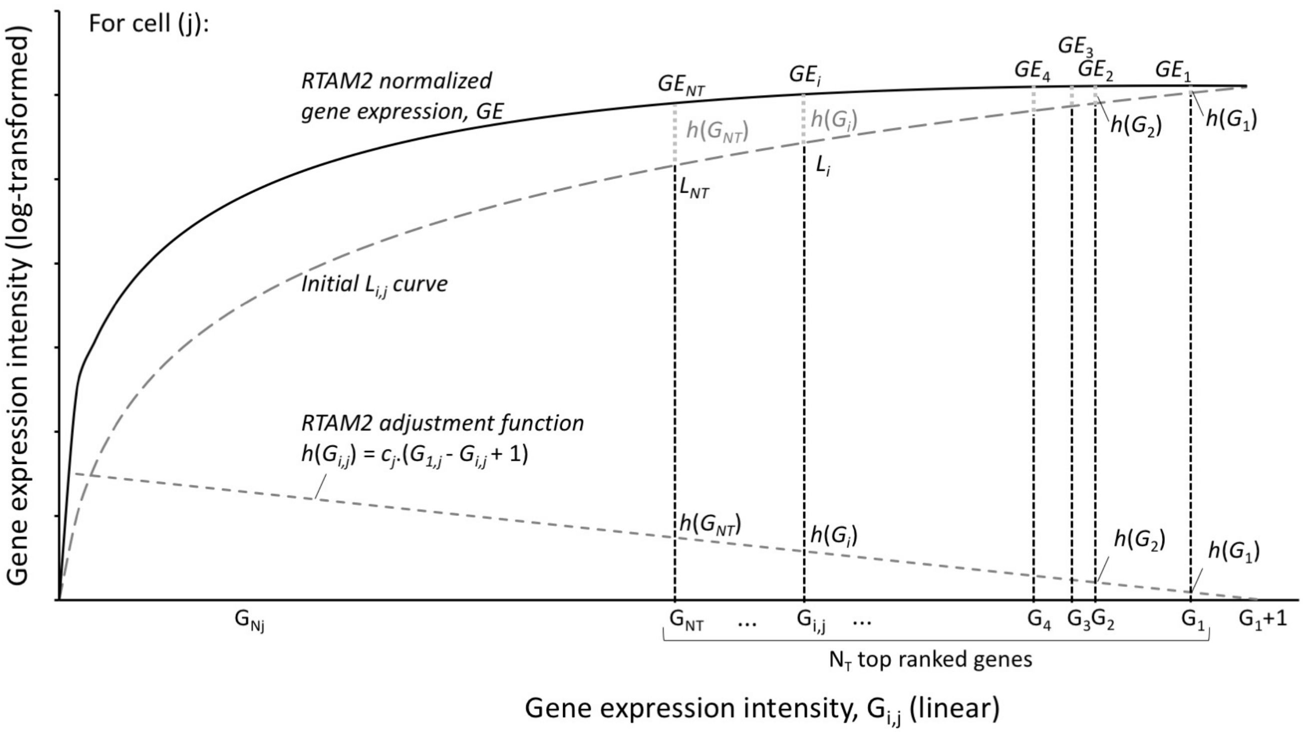


**Figure S3. RTAM2 derivation**

(See the *RTAM Derivation* section in the methods).

The raw transcript counts per gene for cell (j) were partially normalized by scaling of the cellular total transcript count (and constituent gene transcript counts) to a standard library size. The schema above then shows $L_{i,j,}$ (grey line, long dashes), the $\log_2(+1)$-derivative of the partially normalized linear gene expression, $G_{i,j}$. The gene- and cell-specific RTAM2 adjustment function $h(G_{i,j})$ is also shown (grey line, short dashes) and functions to adjust the log-transformed gene expression, $L_{i,j,}$. The final normalized gene expression curve, *GE*, represents $L_{i,j,} + h(G_{i,j})$. The RTAM2 adjustment function $h(G_{i,j})$ is derived for each cell such that it standardizes the sum ($M_j$) of the log-transformed gene expression of the top-ranked genes ($\Sigma L_1, L_2, L_3.. L_i.. L_{NT}$) in each cell (j) to a constant ($M_k$). RTAM2 differs from RTAM1 in that the per gene adjustment is calculated from the gene's expression intensity rather than the gene's expression rank.

**Figures S4-S8 (on following pages). Single cell RNA-seq normalization strategies – comparison of the means of the HKG or UEG detected in each cell**

Plots depicting scRNA-seq data from B cell lineage cells from the bone marrows of MM patients MM238 (Figure S4), MM199 (Figure S5-6) and MM244 (Figure S7-8). In each figure, the raw (pre-normalized) transcript counts per gene per cell are shown in the upper left plot. Each grey dot represents an integer transcript count for one or more genes in a single cell. Cells are ranked on the x-axis by their total transcript count (defined by unique molecular identifiers, UMI) in descending order from large (deeply sequenced) cells on the left to small (shallowly sequenced) cells on the right. The subsequent plots in each figure show the same sample data following normalization using TPM, SCRAN, SCONE, scTransform, RTAM1 or RTAM2 methods. The normalized data is log-transformed. For figures 1, S5 and S7, the methods are compared by examining the mean expression (blue) in each cell of a set of curated house-keeping genes[10] (HKG), known to be broadly expressed, omitting any 'dropout' results in individual cells with undetected expression (zero transcripts). For figures S4, S6 and S8, the methods are similarly compared by examining the mean expression (red) of the ubiquitously expressed genes (UEG) detected in each cell. ). UEG are defined for each sample as the set of genes with detectable expression in >95% of the cells in the dataset and thus represent the largest possible set of genes that are commonly expressed across the cells (allowing pan-cellular comparison) with a dropout or non-expression rate of less than 5%.

Fig. S4

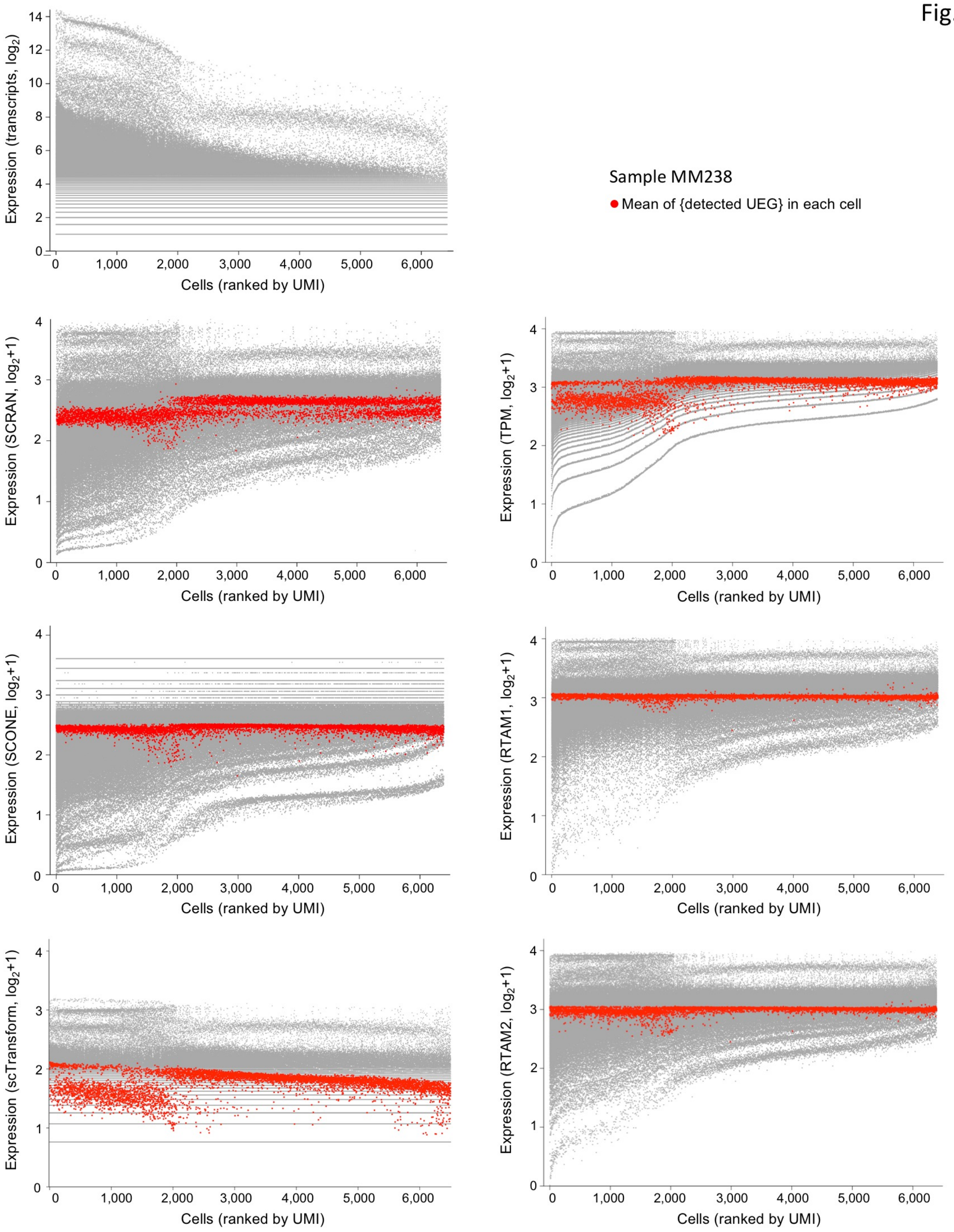
Sample MM238
Mean of {detected UEG} in each cell
Expression (transcripts, log2)
Expression (SCRAN, log2+1)
Expression (TPM, log2+1)
Expression (SCONE, log2+1)
Expression (RTAM1, log2+1)
Expression (scTransform, log2+1)
Expression (RTAM2, log2+1)
Cells (ranked by UMI)

Fig. S5

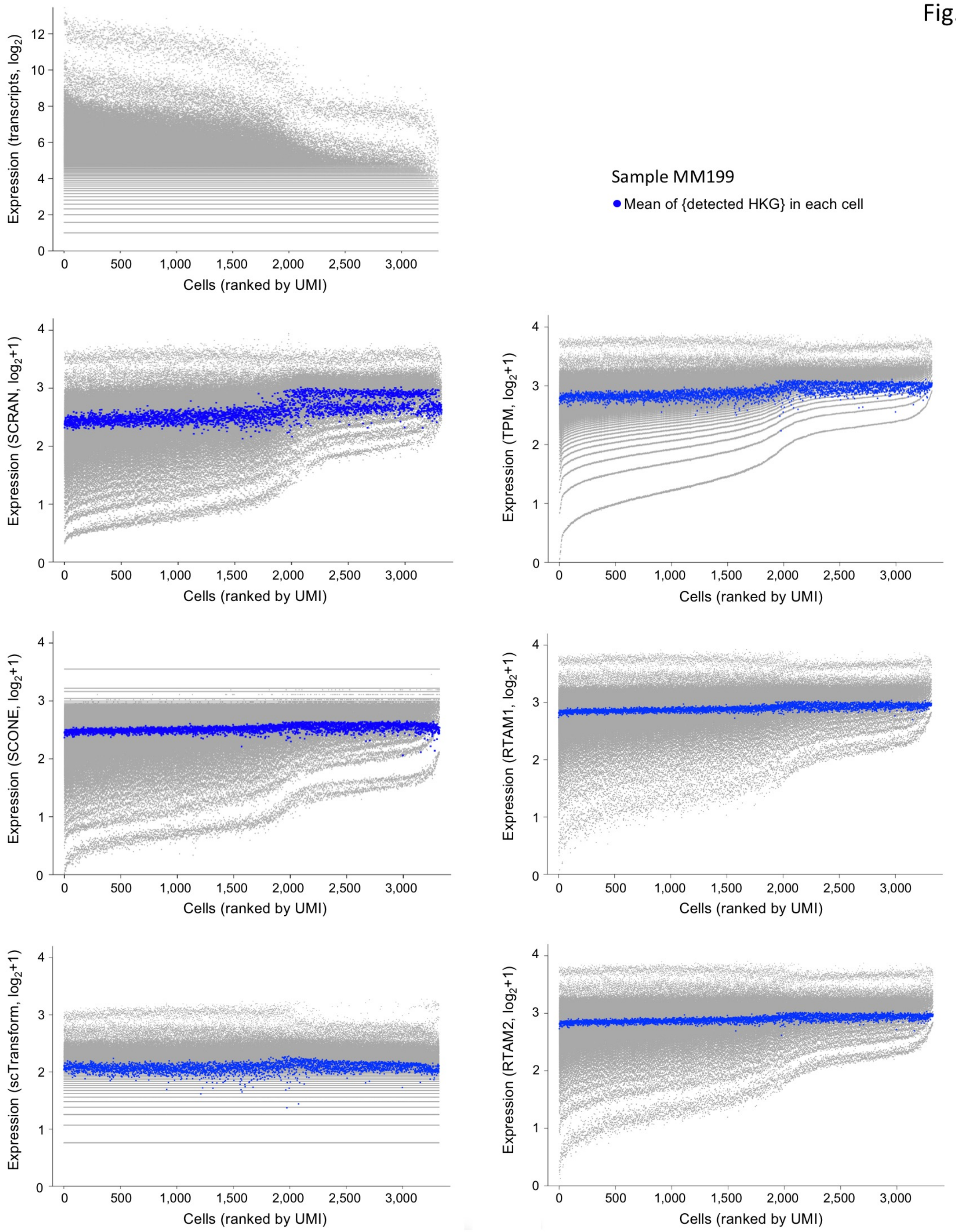
Sample MM199
Mean of {detected HKG} in each cell
Expression (transcripts, log2)
Expression (SCRAN, log2+1)
Expression (TPM, log2+1)
Expression (SCONE, log2+1)
Expression (RTAM1, log2+1)
Expression (scTransform, log2+1)
Expression (RTAM2, log2+1)
Cells (ranked by UMI)
0
500
1,000
1,500
2,000
2,500
3,000

Fig. S6

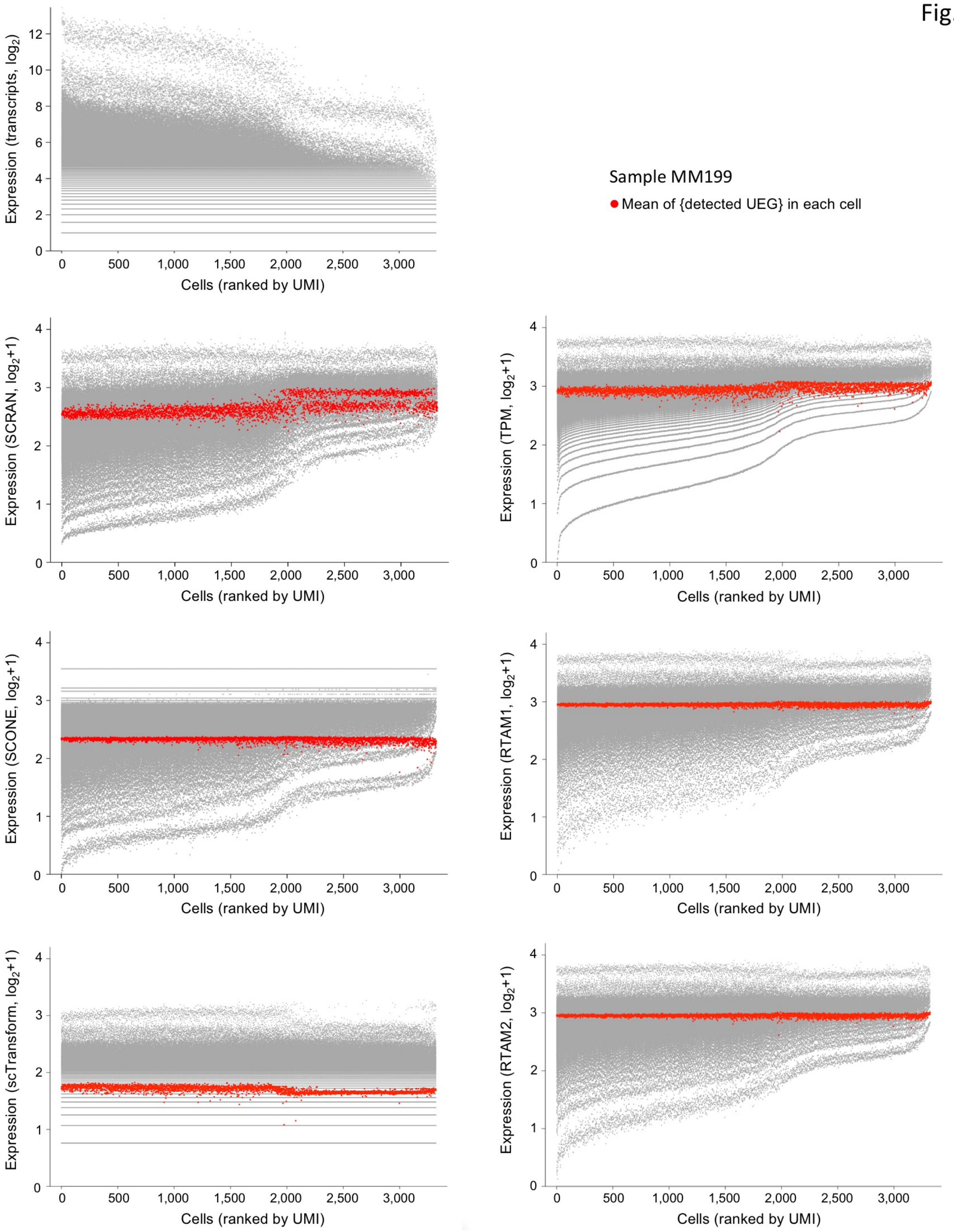
Sample MM199
Mean of {detected UEG} in each cell
Expression (transcripts, $log_2$)
Expression (SCRAN, $log_2$+1)
Expression (TPM, $log_2$+1)
Expression (SCONE, $log_2$+1)
Expression (RTAM1, $log_2$+1)
Expression (scTransform, $log_2$+1)
Expression (RTAM2, $log_2$+1)
Cells (ranked by UMI)
0
500
1,000
1,500
2,000
2,500
3,000

Fig. S7

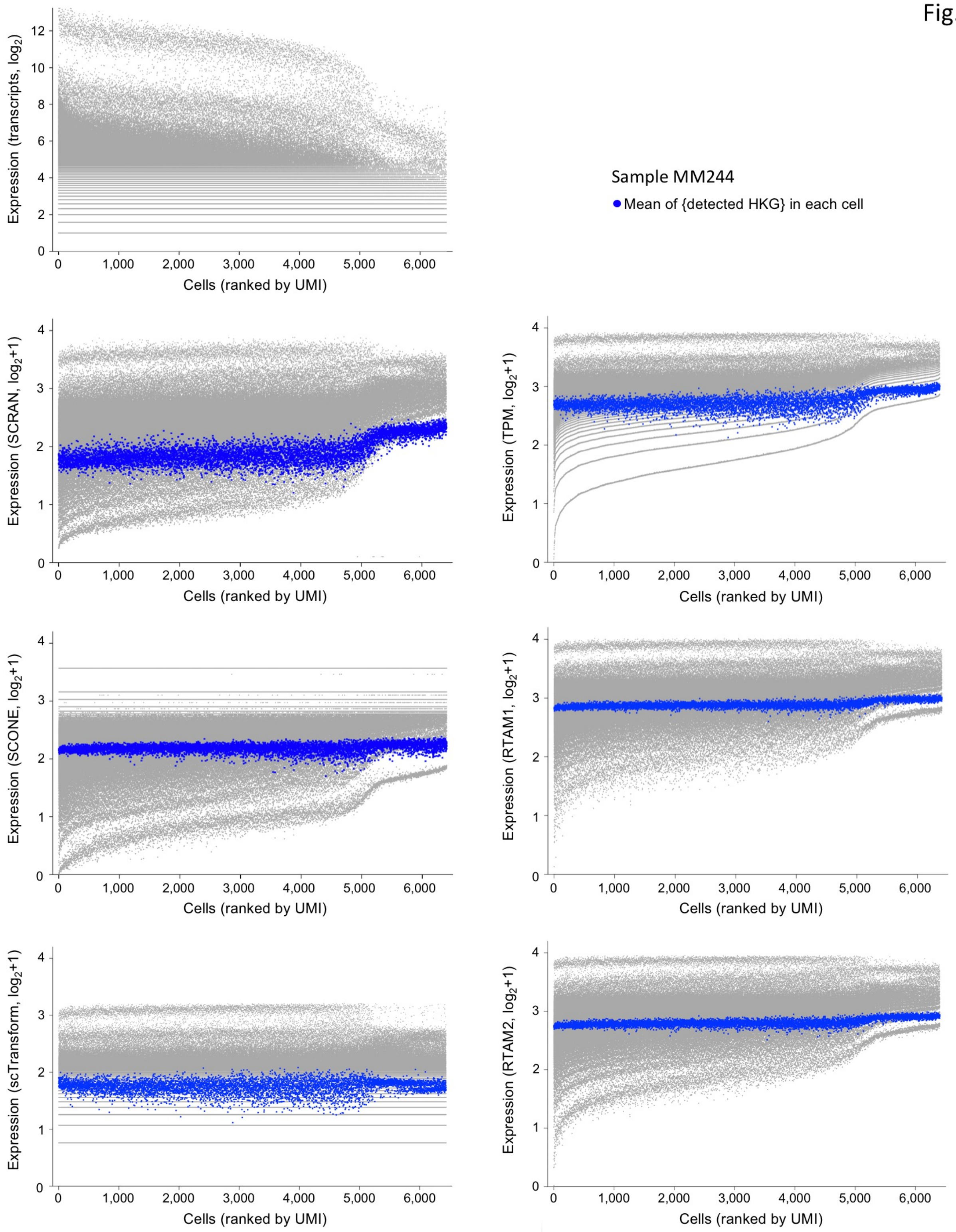
Sample MM244
Mean of {detected HKG} in each cell
Expression (transcripts, log2)
Expression (SCRAN, log2+1)
Expression (TPM, log2+1)
Expression (SCONE, log2+1)
Expression (RTAM1, log2+1)
Expression (scTransform, log2+1)
Expression (RTAM2, log2+1)
Cells (ranked by UMI)
0
1,000
2,000
3,000
4,000
5,000
6,000

Fig. S8

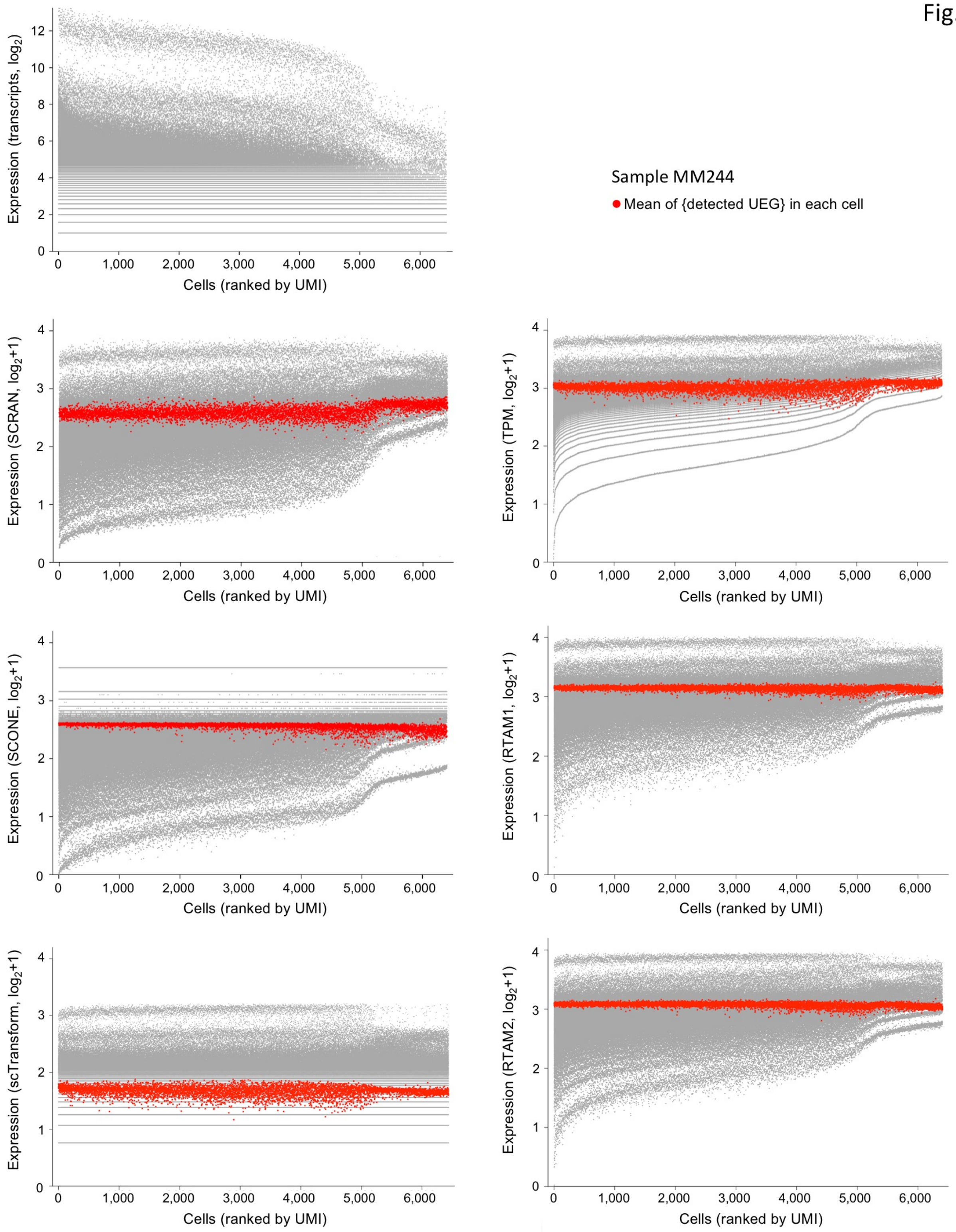
Sample MM244
Mean of {detected UEG} in each cell
Expression (transcripts, log2)
Expression (SCRAN, log2+1)
Expression (TPM, log2+1)
Expression (SCONE, log2+1)
Expression (RTAM1, log2+1)
Expression (scTransform, log2+1)
Expression (RTAM2, log2+1)
Cells (ranked by UMI)
0
1,000
2,000
3,000
4,000
5,000
6,000

**Figures S9-S14 (on following pages). Single cell RNA-seq normalization strategies – comparison of the single cell means of HKG or UEG using imputation of 'dropout' null values**

Plots depicting scRNA-seq data from B cell lineage cells from the bone marrows of MM patients MM238 (Figure S9-10), MM199 (Figure S11-12) and MM244 (Figure S13-14). In each figure, the raw (pre-normalized) transcript counts per gene per cell are shown in the upper left plot. Each grey dot represents an integer transcript count for one or more genes in a single cell. Cells are ranked on the x-axis by their total transcript count (defined by unique molecular identifiers, UMI) in descending order from large (deeply sequenced) cells on the left to small (shallowly sequenced) cells on the right. The subsequent plots in each figure show the same sample data following normalization using TPM, SCRAN, SCONE, scTransform, RTAM1 or RTAM2 methods. The normalized data is log-transformed. For figures S9, S11 and S13, the methods are compared by examining the mean expression (blue) in each cell of a set of curated house-keeping genes[10] (HKG), known to be broadly expressed. For figures S10, S12 and S14, the methods are similarly compared by examining the mean expression (red) in each cell of ubiquitously expressed genes (UEG). UEG are defined for each sample as the set of genes with detectable expression in >95% of the cells in the dataset and thus represent the largest possible set of genes that are commonly expressed across the cells (allowing pan-cellular comparison) with a dropout or non-expression rate of less than 5%. To include all HKG or all UEG in the analysis of each cell, whilst minimizing the influence of null expression values (which are potentially due to signal "dropout") on the cellular HKG means, or UEG means, the expected results for zero-value genes were first imputed by mean substitution calculated across all cells.

Fig. S9

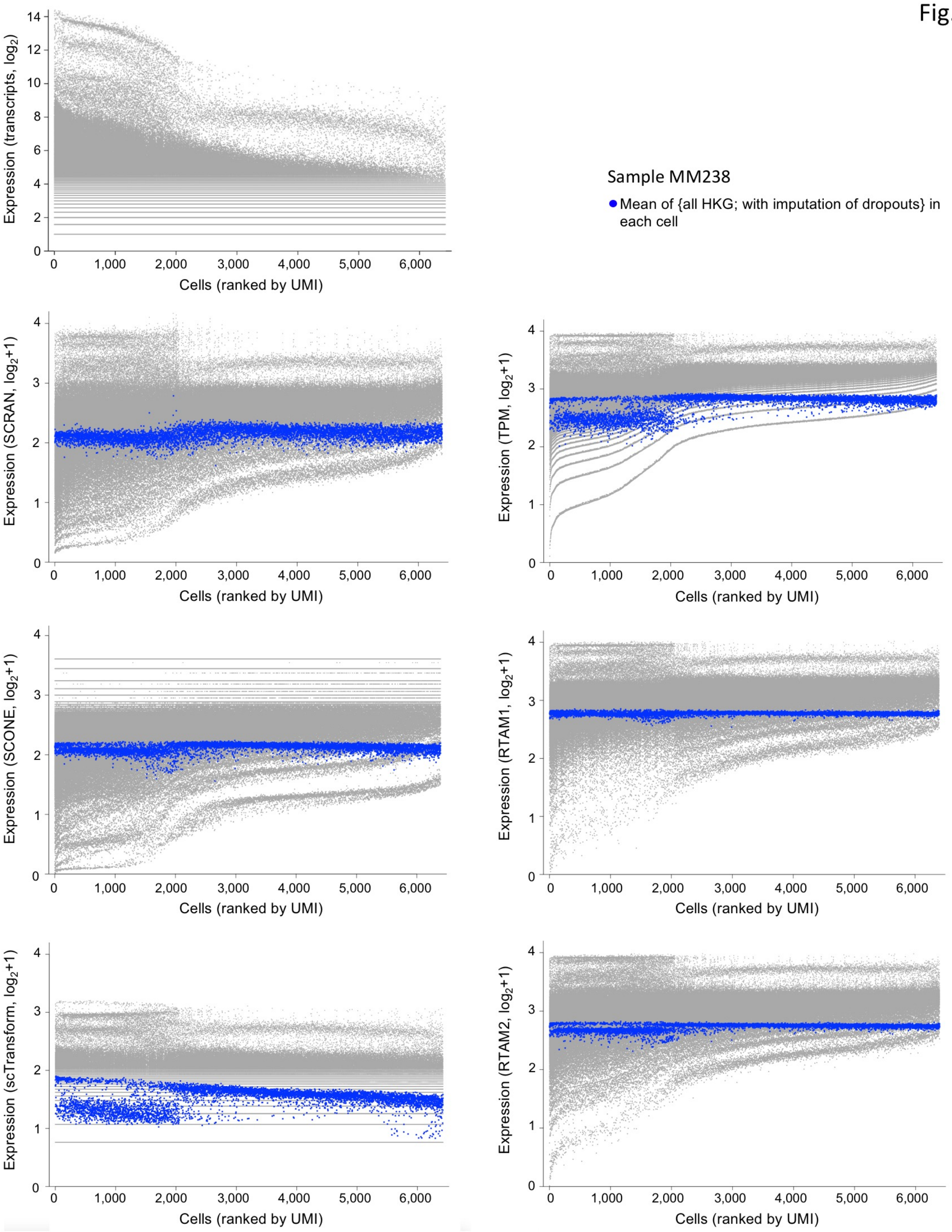

Sample MM238
Mean of {all HKG; with imputation of dropouts} in each cell
Expression (transcripts, log2)
Expression (SCRAN, log2+1)
Expression (TPM, log2+1)
Expression (SCONE, log2+1)
Expression (RTAM1, log2+1)
Expression (scTransform, log2+1)
Expression (RTAM2, log2+1)
Cells (ranked by UMI)
0
1,000
2,000
3,000
4,000
5,000
6,000

Fig. S10

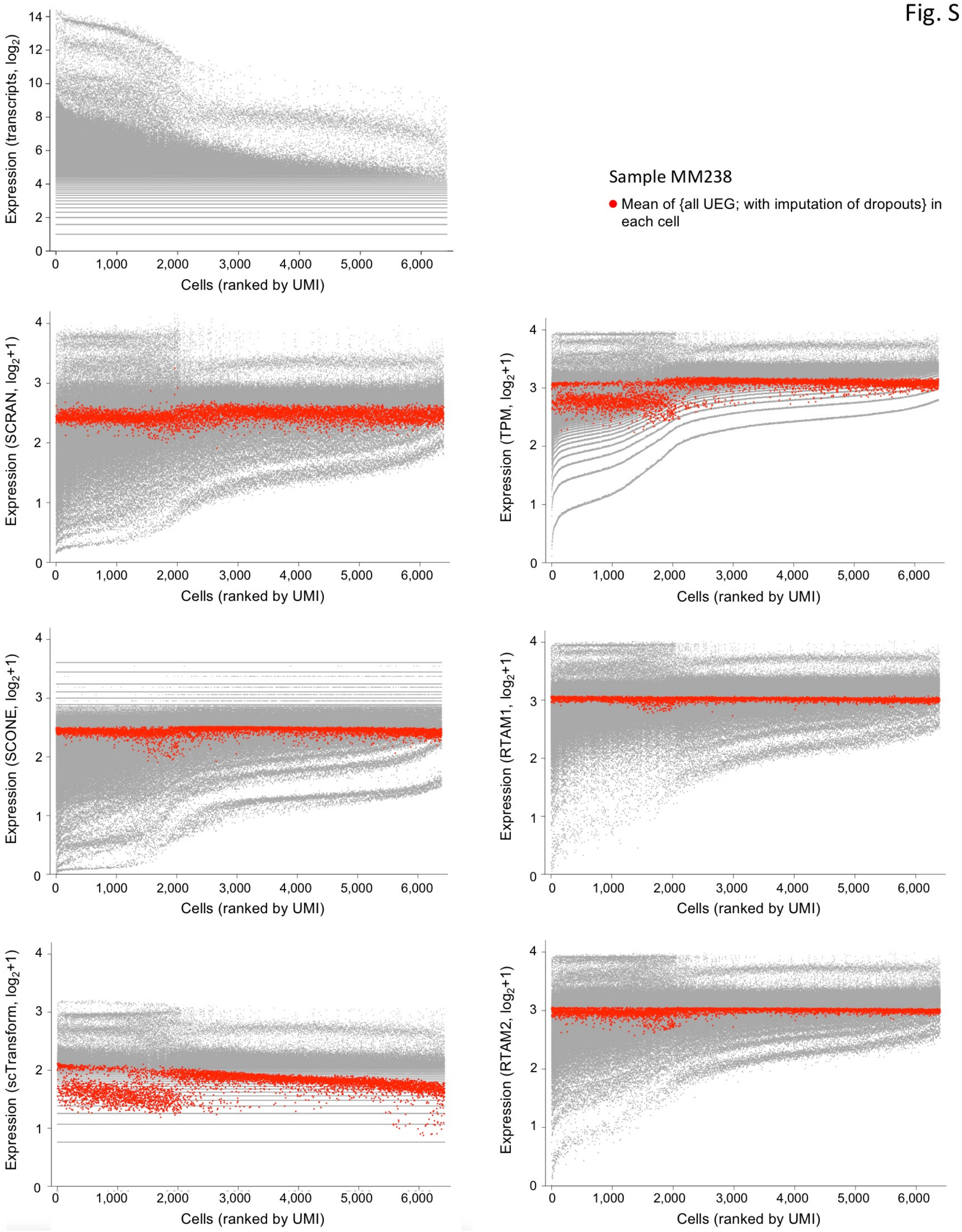

Sample MM238
Mean of {all UEG; with imputation of dropouts} in each cell
Expression (transcripts, log2)
Expression (SCRAN, log2+1)
Expression (TPM, log2+1)
Expression (SCONE, log2+1)
Expression (RTAM1, log2+1)
Expression (scTransform, log2+1)
Expression (RTAM2, log2+1)
Cells (ranked by UMI)
0
1,000
2,000
3,000
4,000
5,000
6,000

Fig. S11

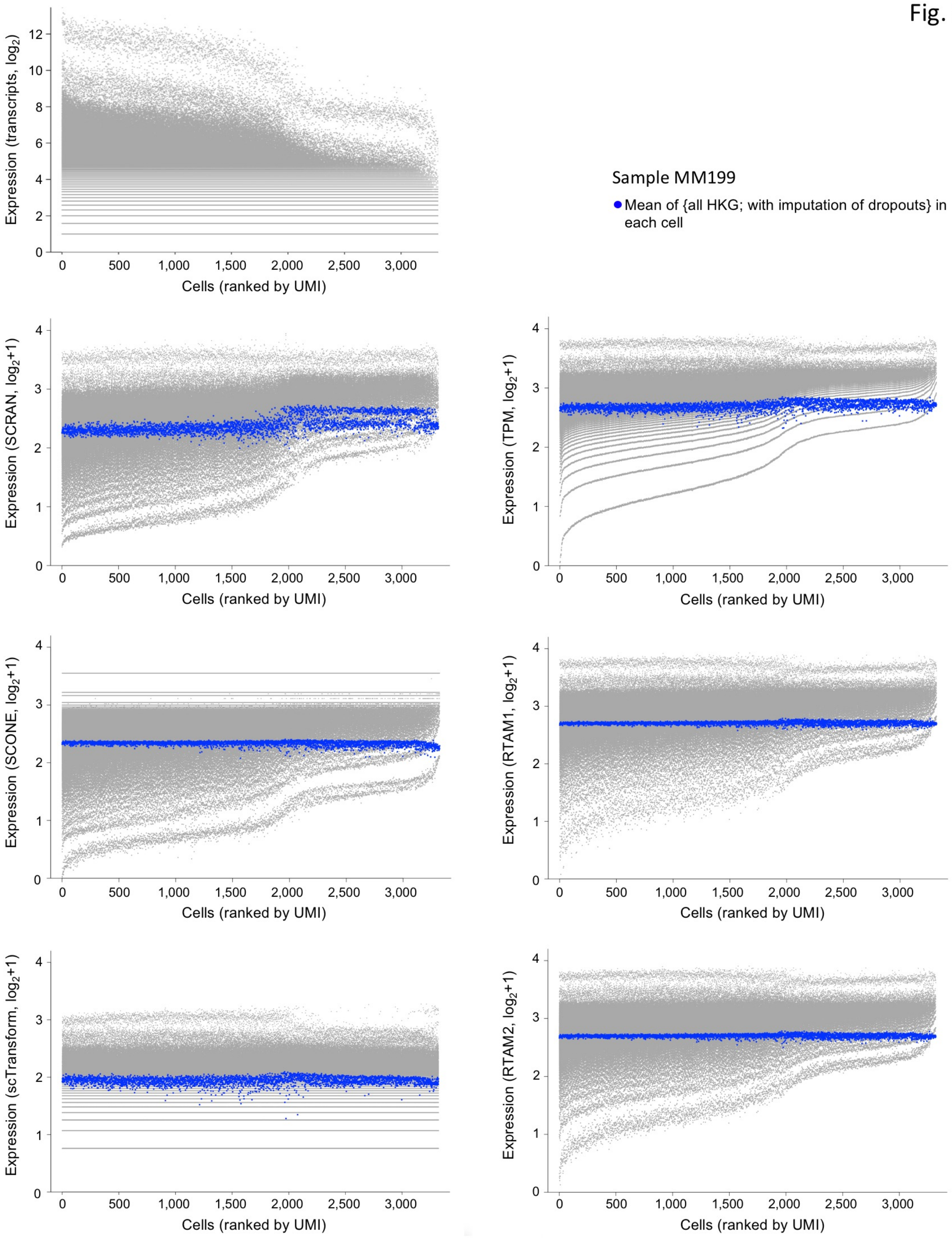
Sample MM199
Mean of {all HKG; with imputation of dropouts} in each cell
Expression (transcripts, log2)
Expression (SCRAN, log2+1)
Expression (TPM, log2+1)
Expression (SCONE, log2+1)
Expression (RTAM1, log2+1)
Expression (scTransform, log2+1)
Expression (RTAM2, log2+1)
Cells (ranked by UMI)
0
500
1,000
1,500
2,000
2,500
3,000

Fig. S12

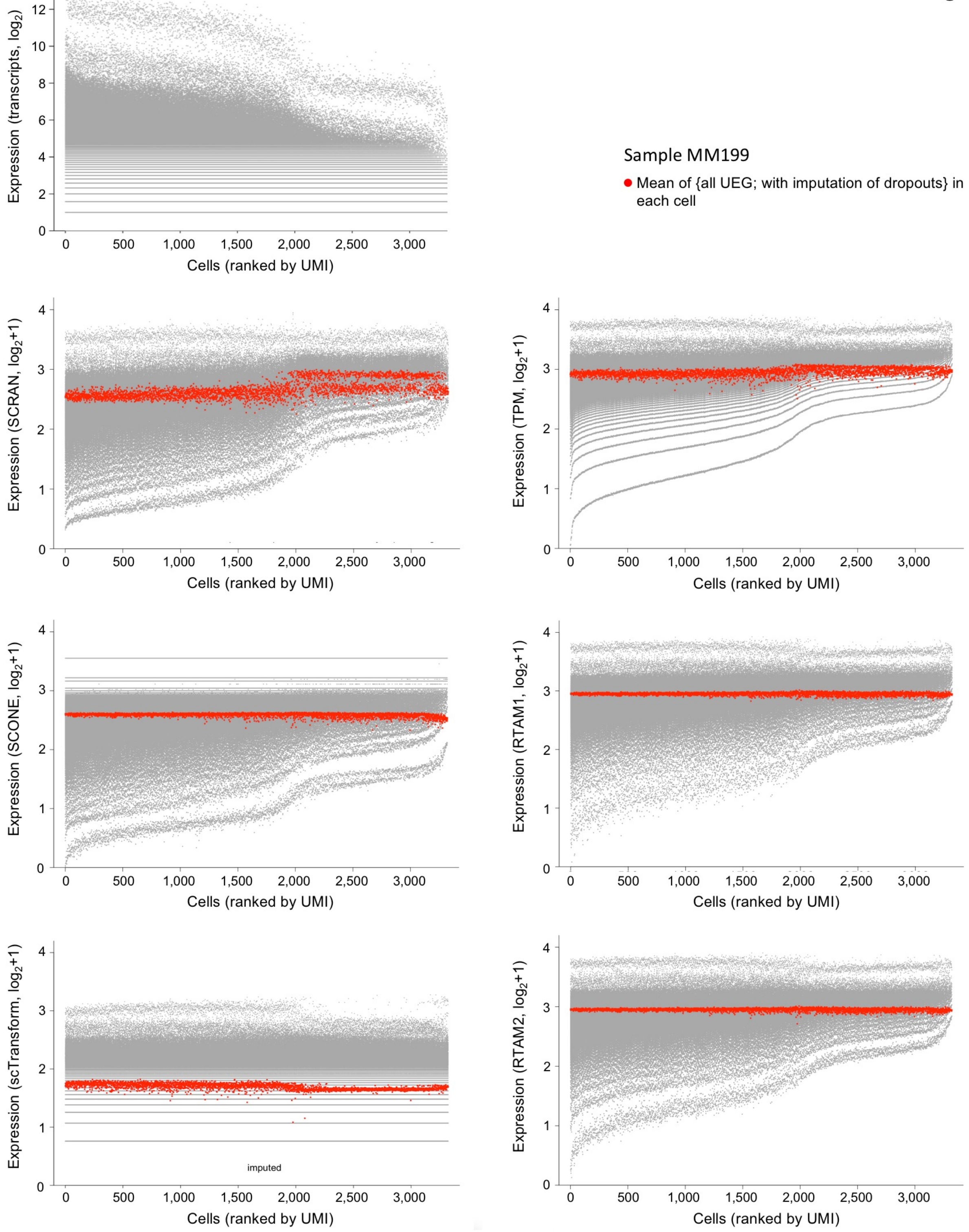
Sample MM199
Mean of {all UEG; with imputation of dropouts} in each cell
Expression (transcripts, log2)
Expression (SCRAN, log2+1)
Expression (TPM, log2+1)
Expression (SCONE, log2+1)
Expression (RTAM1, log2+1)
Expression (scTransform, log2+1)
Expression (RTAM2, log2+1)
imputed
Cells (ranked by UMI)
0
500
1,000
1,500
2,000
2,500
3,000

Fig. S13

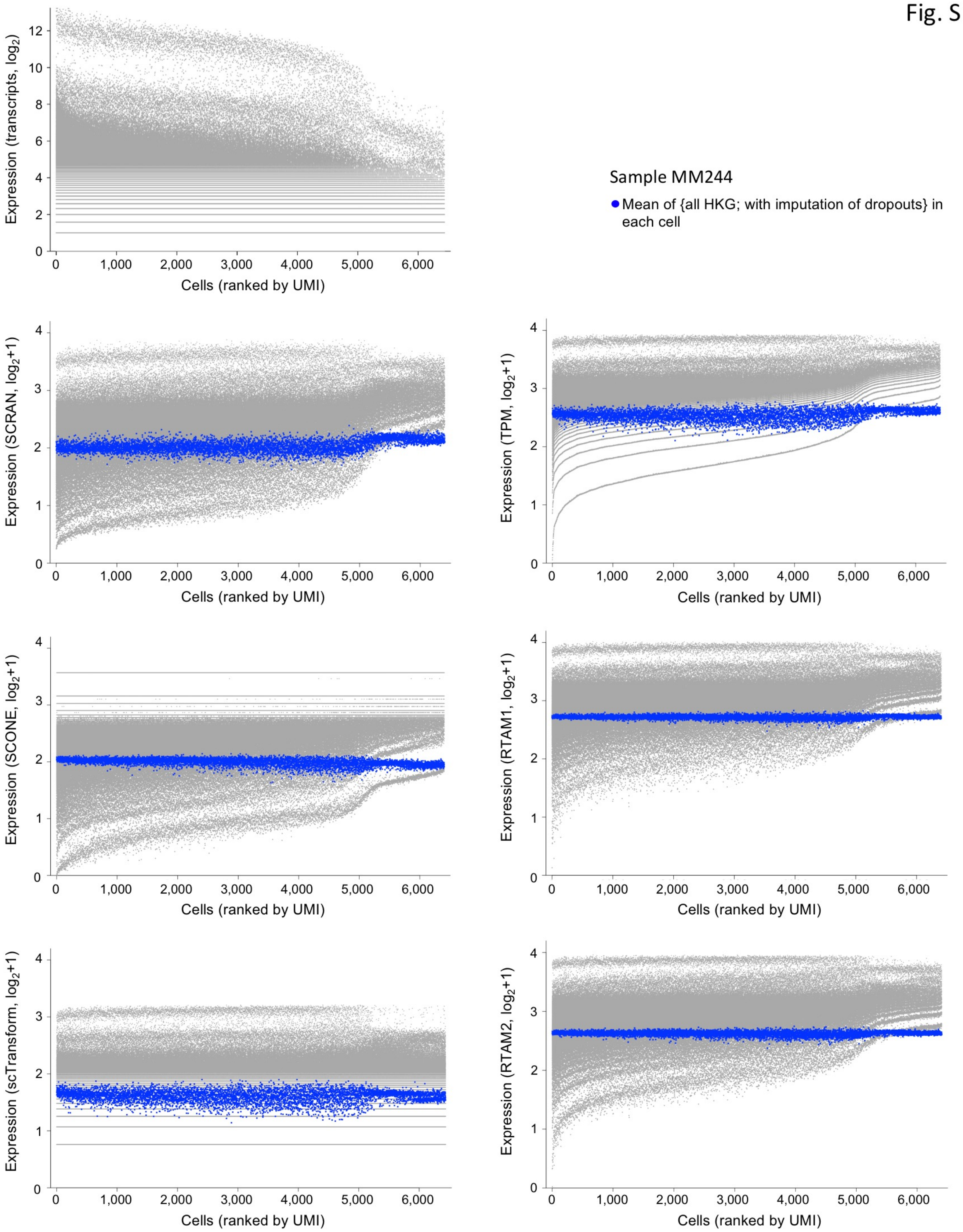
Sample MM244
Mean of {all HKG; with imputation of dropouts} in each cell
Expression (transcripts, log2)
Expression (SCRAN, log2+1)
Expression (TPM, log2+1)
Expression (SCONE, log2+1)
Expression (RTAM1, log2+1)
Expression (scTransform, log2+1)
Expression (RTAM2, log2+1)
Cells (ranked by UMI)
0
1,000
2,000
3,000
4,000
5,000
6,000
0
2
4
6
8
10
12
1
3

Fig. S14

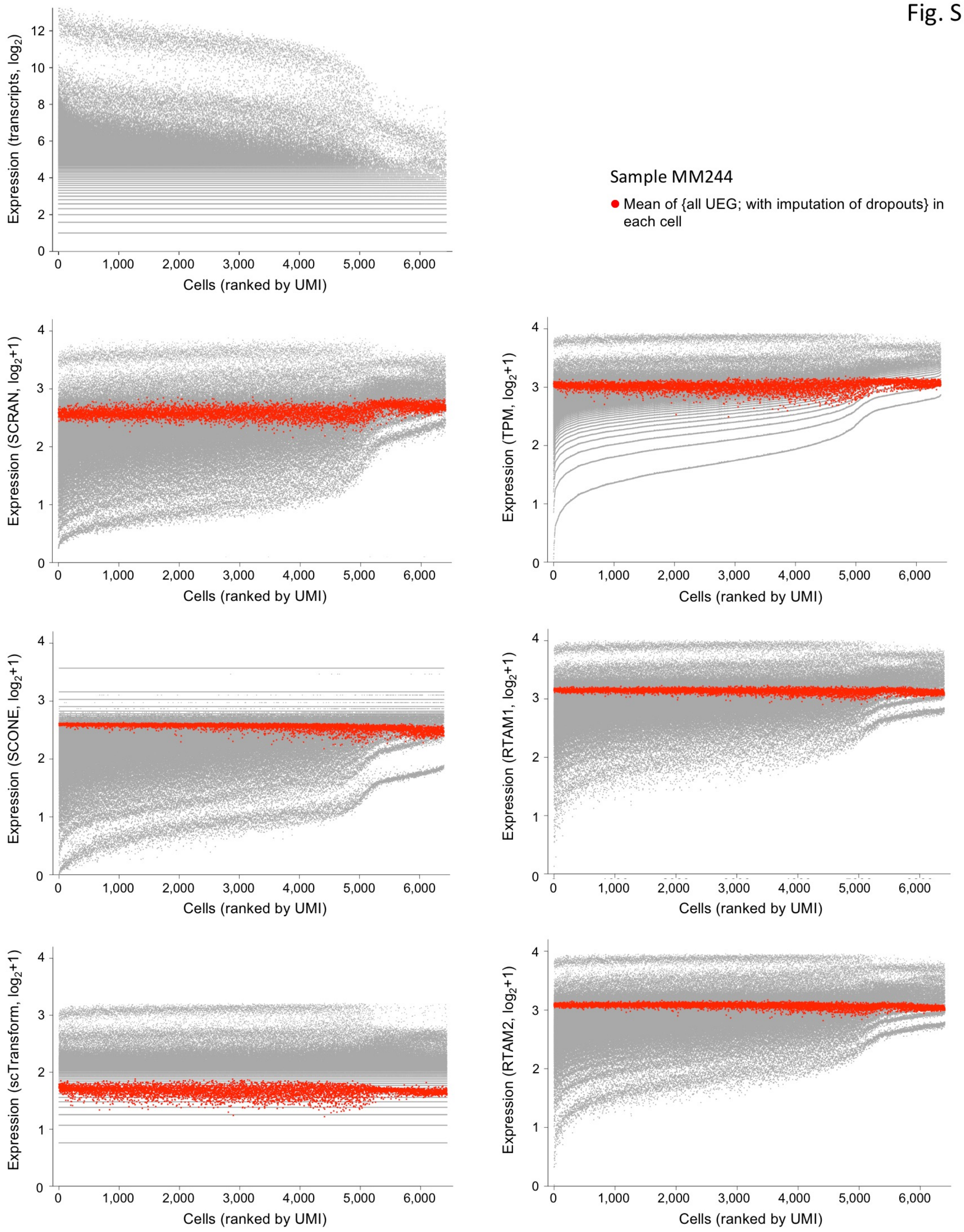
Sample MM244
Mean of {all UEG; with imputation of dropouts} in each cell
Expression (transcripts, $\log_2$)
Expression (SCRAN, $\log_2$+1)
Expression (TPM, $\log_2$+1)
Expression (SCONE, $\log_2$+1)
Expression (RTAM1, $\log_2$+1)
Expression (scTransform, $\log_2$+1)
Expression (RTAM2, $\log_2$+1)
Cells (ranked by UMI)
0
1,000
2,000
3,000
4,000
5,000
6,000

**Figures S15-S20 (on following pages). Single cell RNA-seq normalization strategies – comparison of single cell medians of HKG or UEG**

Plots depicting scRNA-seq data from B cell lineage cells from the bone marrows of MM patients MM238 (Figure S15-16), MM199 (Figure S17-18) and MM244 (Figure S19-20). In each figure, the raw (pre-normalized) transcript counts per gene per cell are shown in the upper left plot. Each grey dot represents an integer transcript count for one or more genes in a single cell. Cells are ranked on the x-axis by their total transcript count in descending order from large (deeply sequenced) cells on the left to small (shallowly sequenced) cells on the right. The subsequent plots in each figure show the same sample data following normalization using TPM, SCRAN, SCONE, scTransform, RTAM1 or RTAM2 methods. The normalized data is log-transformed. For figures S15, S17 and S19, the methods are compared by examining the median expression (blue) in each cell of a set of curated house-keeping genes[10] (HKG), known to be broadly expressed. For figures S16, S18 and S20, the methods are similarly compared by examining the median expression (red) in each cell of ubiquitously expressed genes (UEG). UEG are defined for each sample as the set of genes with detectable expression in >95% of the cells in the dataset and thus represent the largest possible set of genes that are commonly expressed across the cells (allowing pan-cellular comparison) with a dropout or non-expression rate of less than 5%. All HKG or UEG are included in the analysis of each cell, including genes with null expression values (which are potentially due to signal "dropout").

Fig. S15

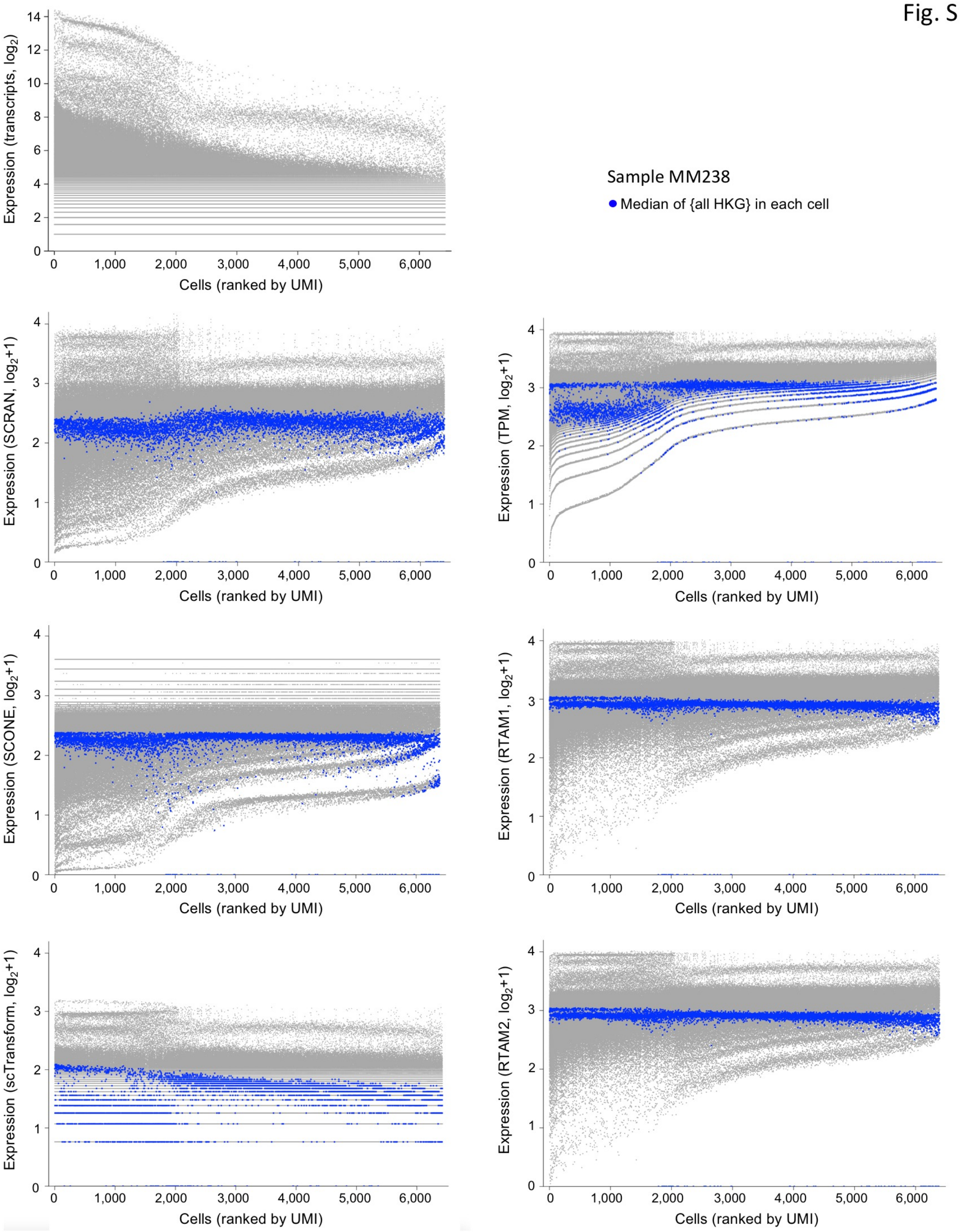
Sample MM238
Median of {all HKG} in each cell
Expression (transcripts, $log_2$)
Expression (SCRAN, $log_2$+1)
Expression (TPM, $log_2$+1)
Expression (SCONE, $log_2$+1)
Expression (RTAM1, $log_2$+1)
Expression (scTransform, $log_2$+1)
Expression (RTAM2, $log_2$+1)
Cells (ranked by UMI)
0
1,000
2,000
3,000
4,000
5,000
6,000

Fig. S16

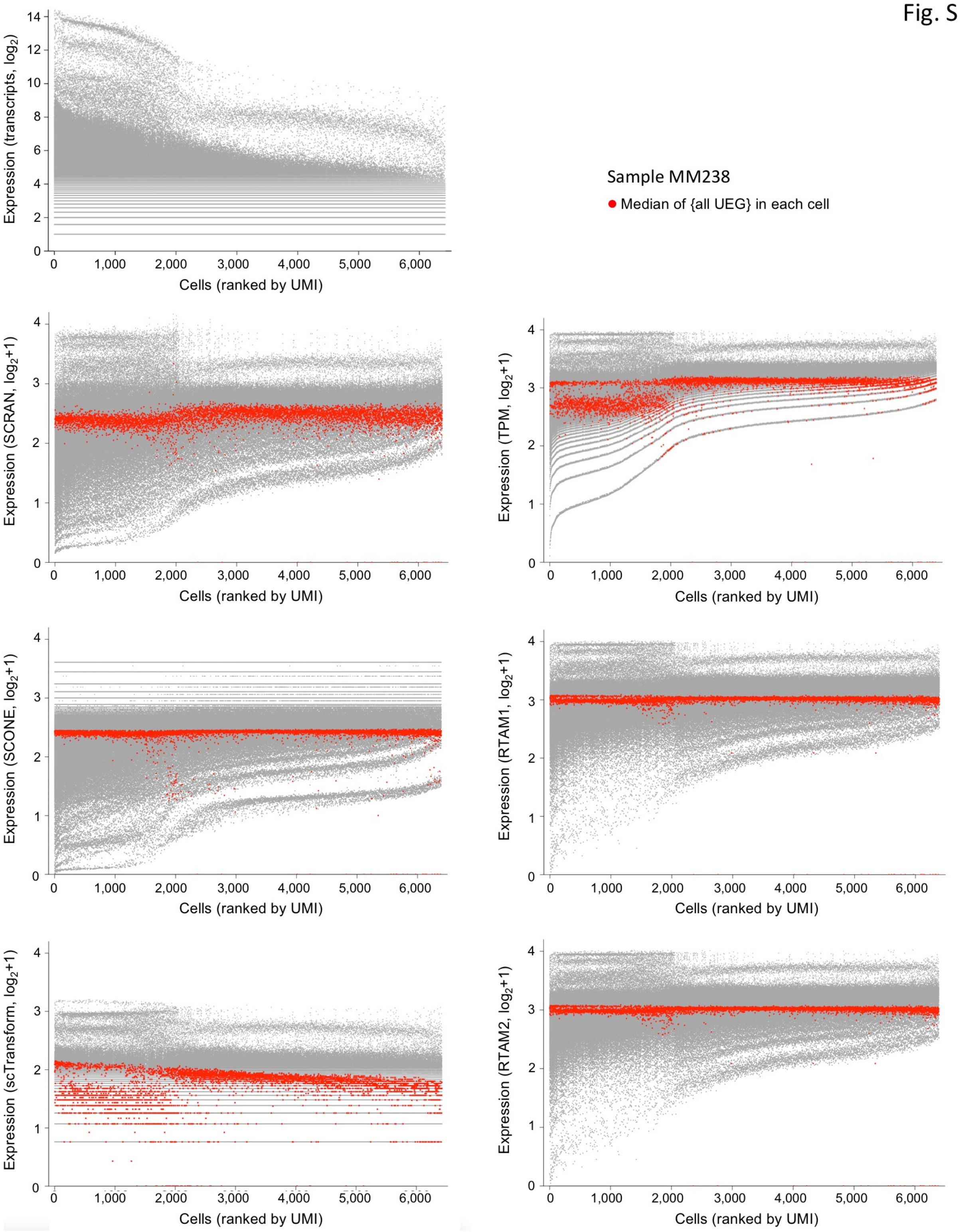
Sample MM238
Median of {all UEG} in each cell
Expression (transcripts, log2)
Expression (SCRAN, log2+1)
Expression (TPM, log2+1)
Expression (SCONE, log2+1)
Expression (RTAM1, log2+1)
Expression (scTransform, log2+1)
Expression (RTAM2, log2+1)
Cells (ranked by UMI)
0
1,000
2,000
3,000
4,000
5,000
6,000

Fig. S17

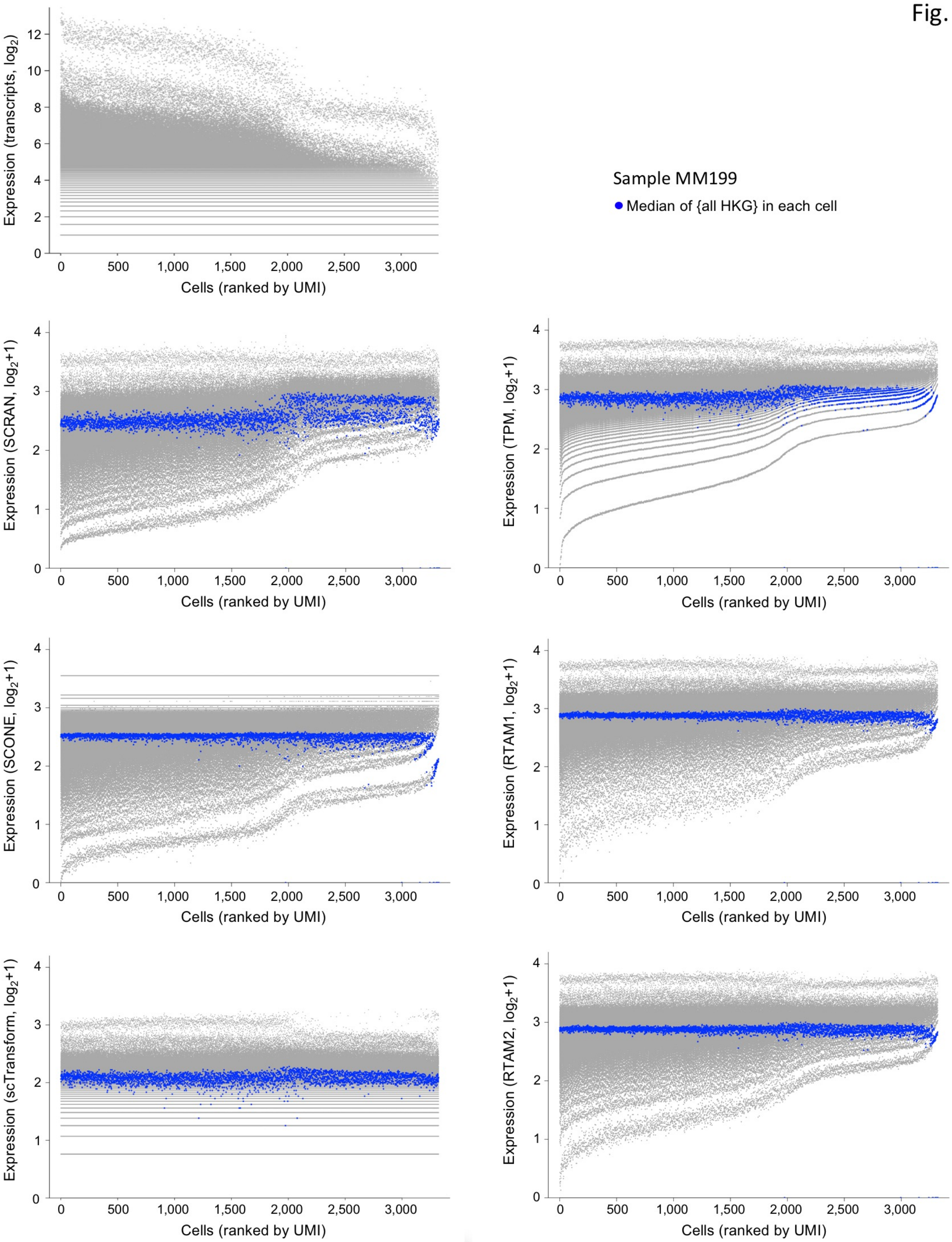
Sample MM199
Median of {all HKG} in each cell
Expression (transcripts, log2)
Expression (SCRAN, log2+1)
Expression (TPM, log2+1)
Expression (SCONE, log2+1)
Expression (RTAM1, log2+1)
Expression (scTransform, log2+1)
Expression (RTAM2, log2+1)
Cells (ranked by UMI)
0
500
1,000
1,500
2,000
2,500
3,000

Fig. S18

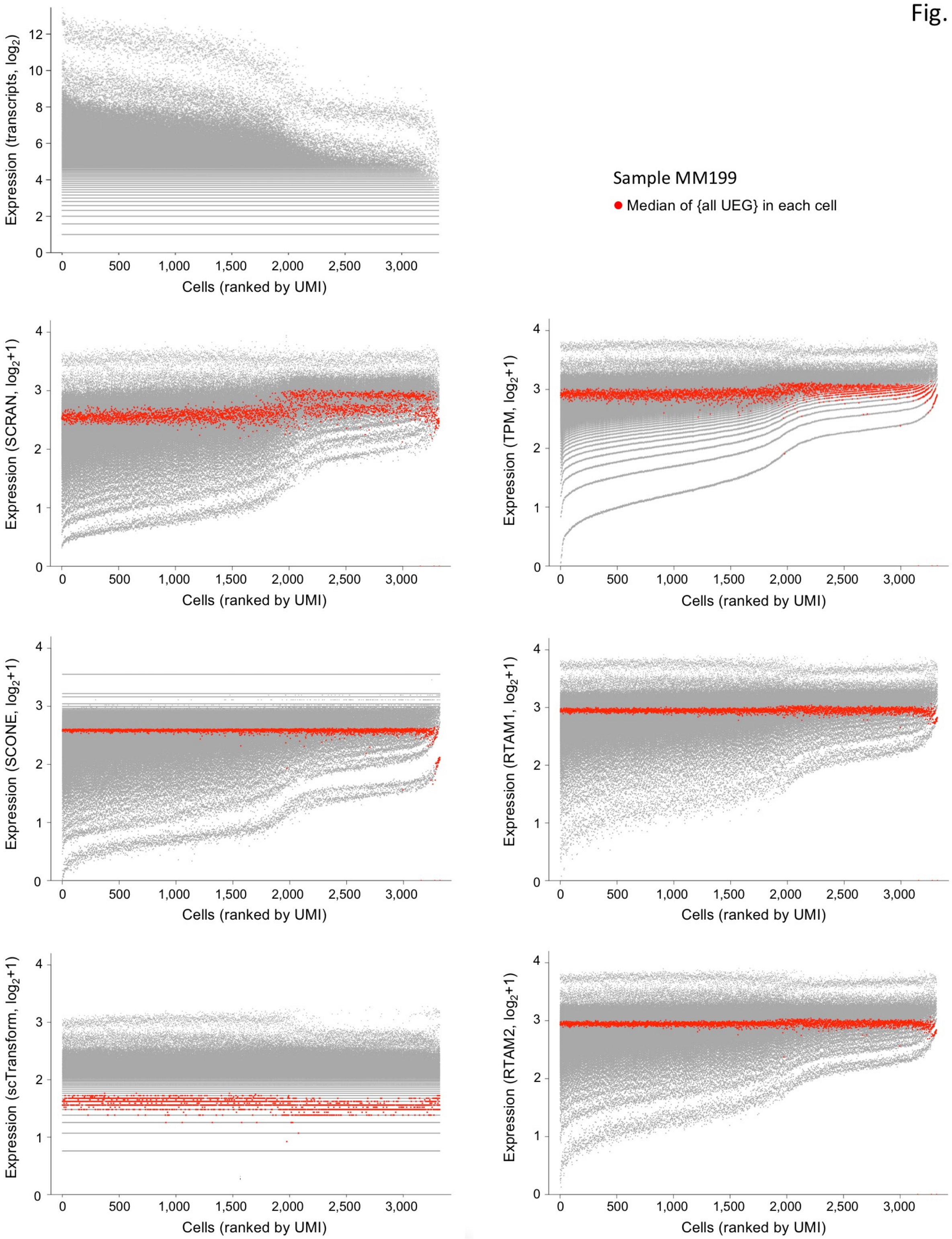
Sample MM199
Median of {all UEG} in each cell
Expression (transcripts, log2)
Expression (SCRAN, log2+1)
Expression (TPM, log2+1)
Expression (SCONE, log2+1)
Expression (RTAM1, log2+1)
Expression (scTransform, log2+1)
Expression (RTAM2, log2+1)
Cells (ranked by UMI)
0
500
1,000
1,500
2,000
2,500
3,000

Fig. S19

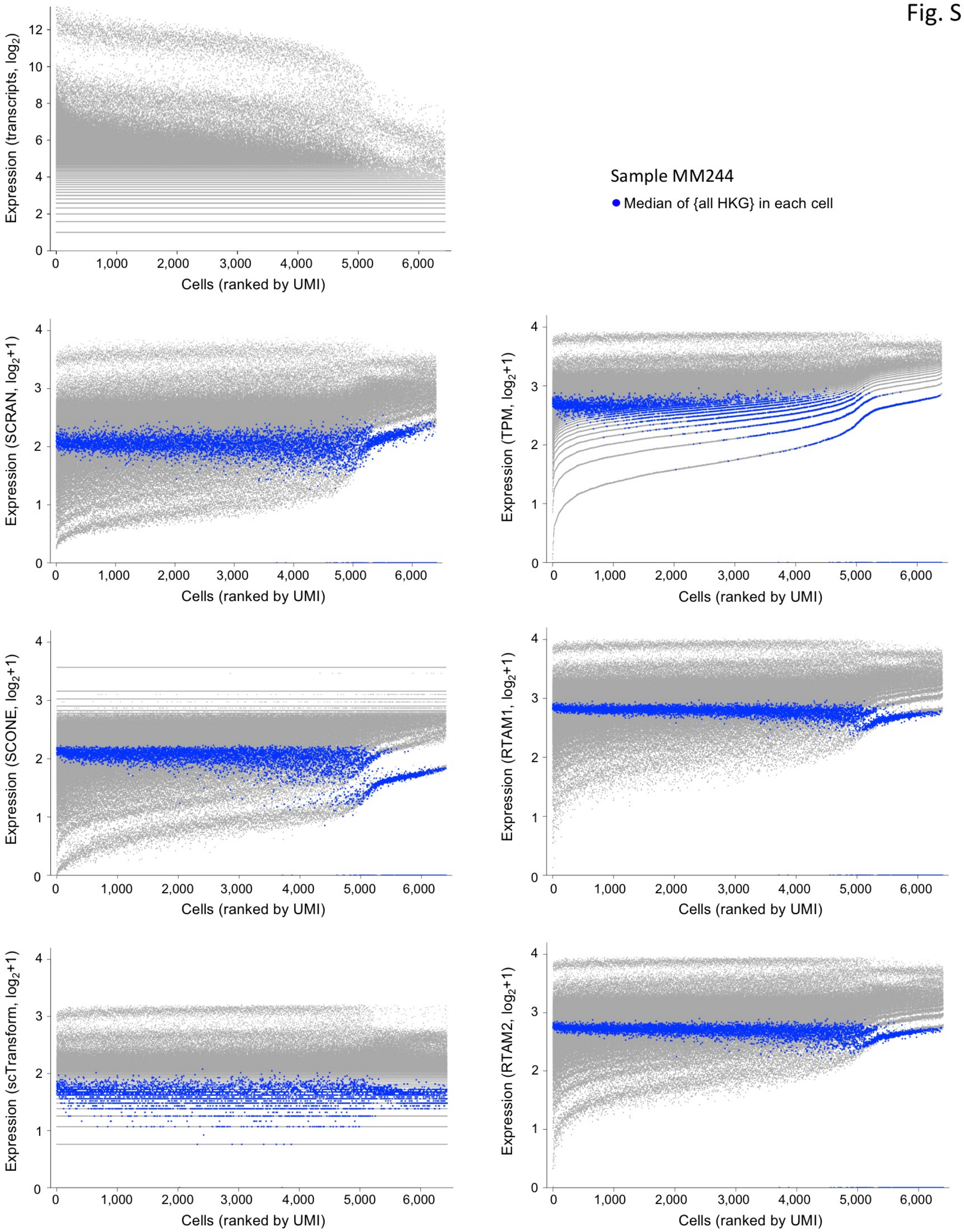
Sample MM244
Median of {all HKG} in each cell
Expression (transcripts, log$_2$)
Expression (SCRAN, log$_2$+1)
Expression (TPM, log$_2$+1)
Expression (SCONE, log$_2$+1)
Expression (RTAM1, log$_2$+1)
Expression (scTransform, log$_2$+1)
Expression (RTAM2, log$_2$+1)
Cells (ranked by UMI)
0
1,000
2,000
3,000
4,000
5,000
6,000

### Fig. S20

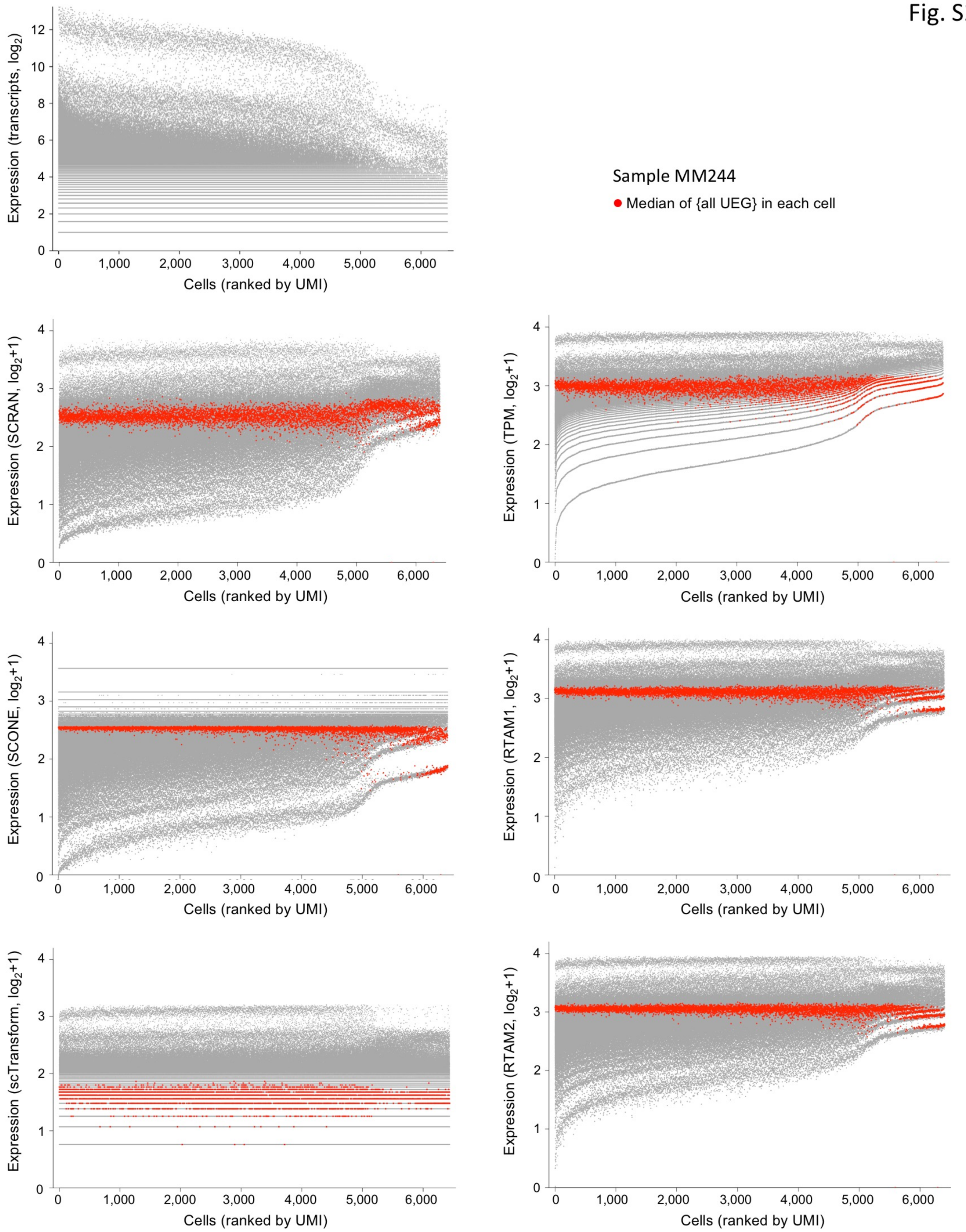
Sample MM244
Median of {all UEG} in each cell
Expression (transcripts, log2)
Expression (SCRAN, log2+1)
Expression (TPM, log2+1)
Expression (SCONE, log2+1)
Expression (RTAM1, log2+1)
Expression (scTransform, log2+1)
Expression (RTAM2, log2+1)
Cells (ranked by UMI)
0
1,000
2,000
3,000
4,000
5,000
6,000

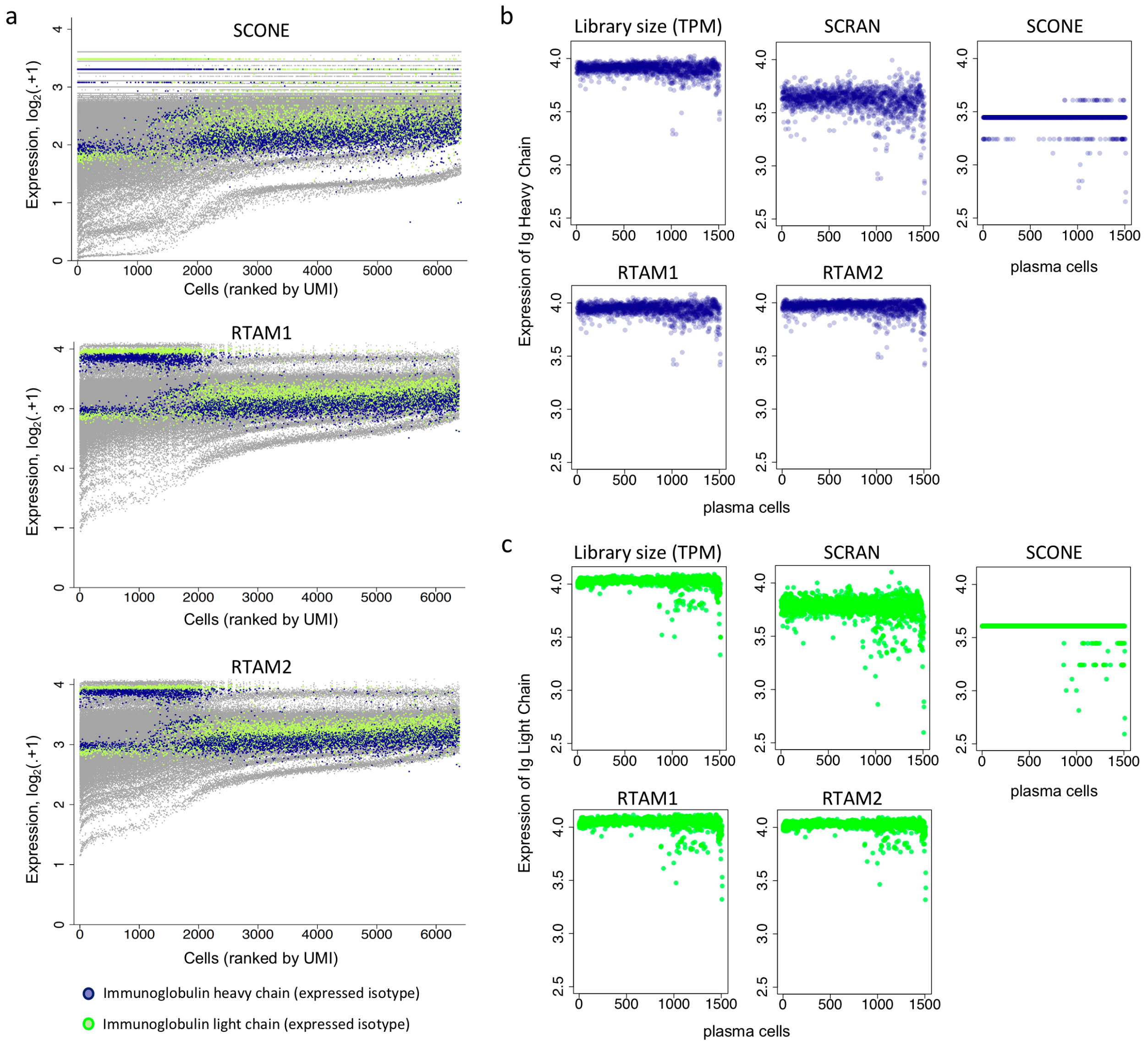


**Figures S21. Additional comparisons of scRNA-seq normalization strategies – immunoglobulin expression**

**a.** Plots depicting log-transformed scRNA-seq data from B cell lineage cells, isolated from the bone marrow of patient MM238, and normalized either by SCONE, RTAM1 or RTAM2. Each grey dot represents a transcript count tier or expression level for one or more genes in a single cell; cells are ranked by total transcript count (defined by unique molecular identifiers, UMI) in descending order. Larger cells (plasma cells) are to the left, smaller cells (B cells) are to the right. For each cell, the expression intensity of the highest-expressed Ig heavy chain and of the highest expressed Ig light chain (the expressed isotype) is plotted in blue and green respectively.

**b.** and **c**. Plots comparing the expression of Ig heavy chain (**b**) and Ig light chain (**c**) in the plasma cell subpopulation in sample MM238 after data normalization by TPM, SCRAN, SCONE, RTAM1 or RTAM2, showing compression of these expression values into a virtual constant by the quantile method employed by SCONE.

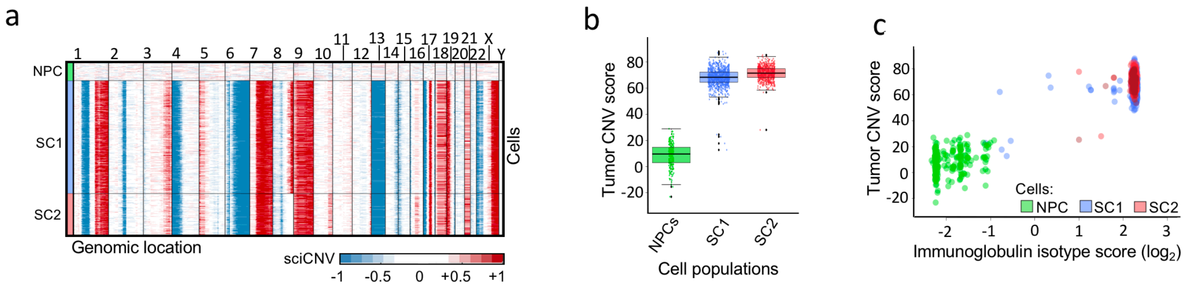


**Figure S22. A Tumor CNV Score separates normal and malignant clone cells.**

**a.** Heatmap showing predicted chromosome copy number gains (red) and losses (blue) in 1724 individual multiple myeloma plasma cells (MMPC), inferred from scRNA-seq using sciCNV. The MMPC are grouped into subclones (SC1, SC2) according to the presence of +8q. The CNVs of 205 normal plasma cells (NPC) are shown as a control. **b.** Identification of malignant cells within scRNA-seq datasets by CNV scores. The MMPC in subclones 1 and 2 (SC1, SC2) are readily distinguished from NPC on the basis of the similarity of their individual sciCNV profiles to the tumor clone mean sciCNV profile, calculated as a 'tumor CNV score'. **c.** Validation of cancer cell identification by Tumor CNV score. The Tumor CNV Scores were plotted again an independent immunoglobulin isotype score, calculated to distinguish cells expressing immunoglobulin of the tumor clone isotype from polyclonal cells expressing other isotypes. Virtually all cells with a high tumor CNV score also expressed immunoglobulin of the tumor isotype, confirming that the tumor CNV score can be used to identify malignant cells. Notably the tumor CNV score can be used to classify cells from all tumors with clonal CNV, unlike immunoglobulin restriction which can only identify tumor cells in lymphoid malignancies.

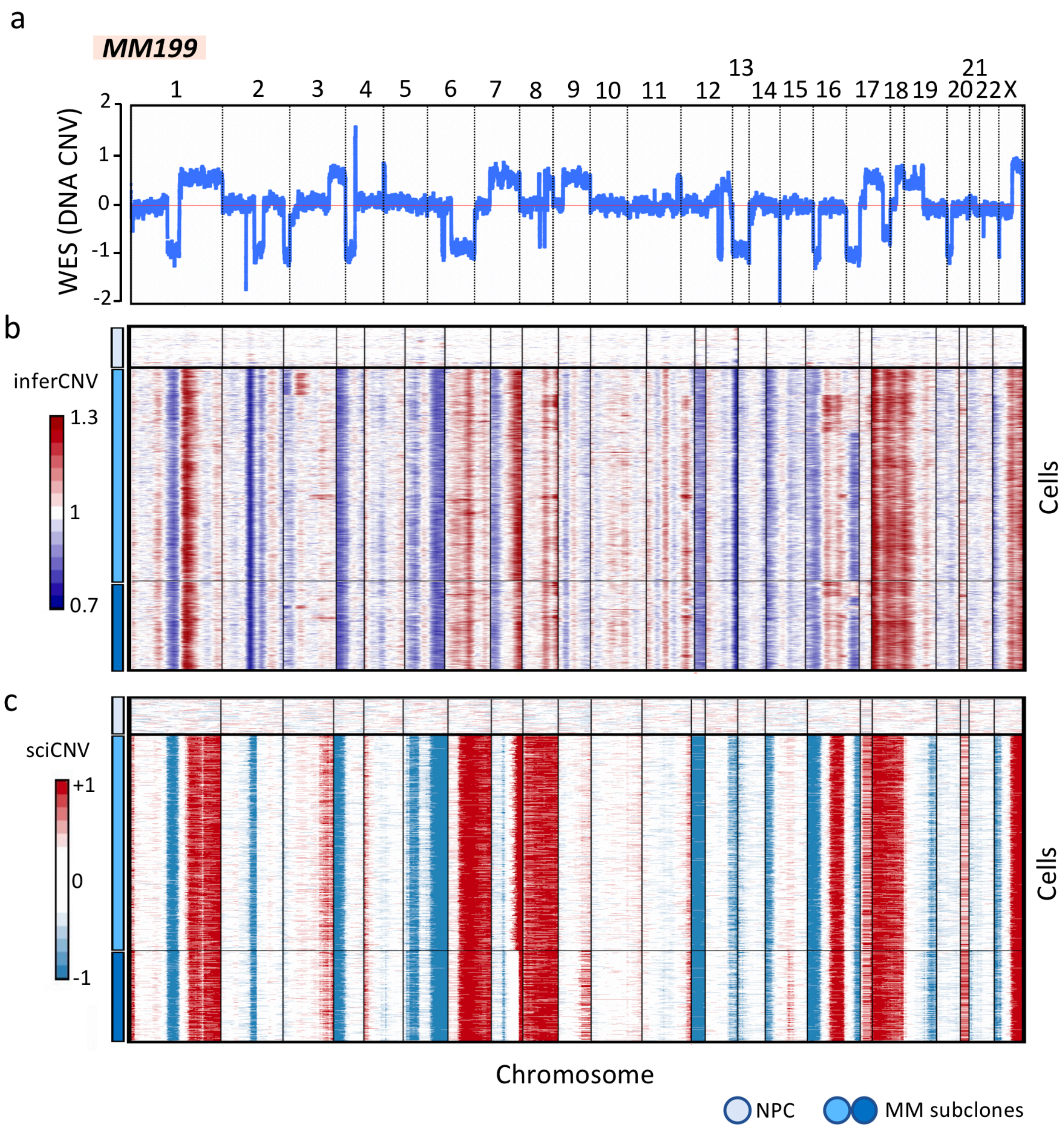


**Figure S23. sciCNV compares favorably with inferCNV in predicting single cell CNV using scRNA-seq – part 1 (sample MM199).**

Single cell CNV profiles, derived by sciCNV or inferCNV for hundreds of individual MM cells using scRNA-seq data, are compared with the tumor bulk CNV profile obtained by DNA-sequencing of pooled tumor cells.

**a.** The tumor bulk CNV profile for sample MM199, generated by DNA whole exome sequencing.

b-c. The CNV profiles of individual plasma cells (n=1724) from MM patient sample MM199, inferred from scRNA-seq using inferCNV (b) or sciCNV (c). The profiles of normal plasma cells (NPCs)(n=205) are shown as controls.

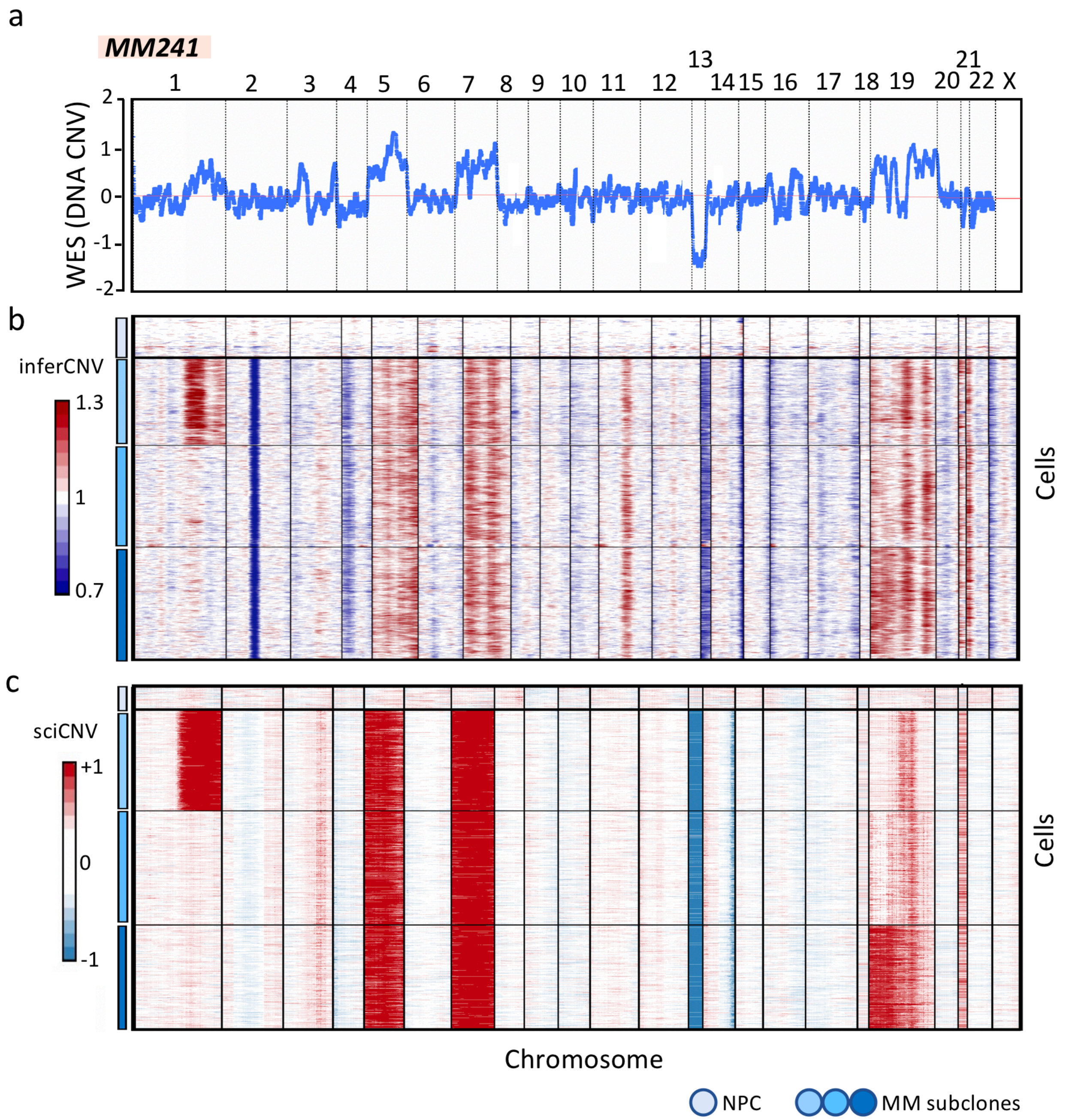


**Figure S24. sciCNV compares favorably with inferCNV in predicting single cell CNV using scRNA-seq – part 2 (sample MM241).**

Single cell CNV profiles, derived by sciCNV or inferCNV for hundreds of individual MM cells using scRNA-seq data, are compared with the tumor bulk CNV profile obtained by DNA-sequencing of pooled tumor cells.

**a.** The tumor bulk CNV profile for sample MM241, generated by DNA whole exome sequencing.

**b-c.** The CNV profiles of individual plasma cells (n=2806) from MM patient sample MM241, inferred from scRNA-seq using inferCNV (b) or sciCNV (c). The profiles of normal plasma cells (NPCs)(n=205) are shown as controls.

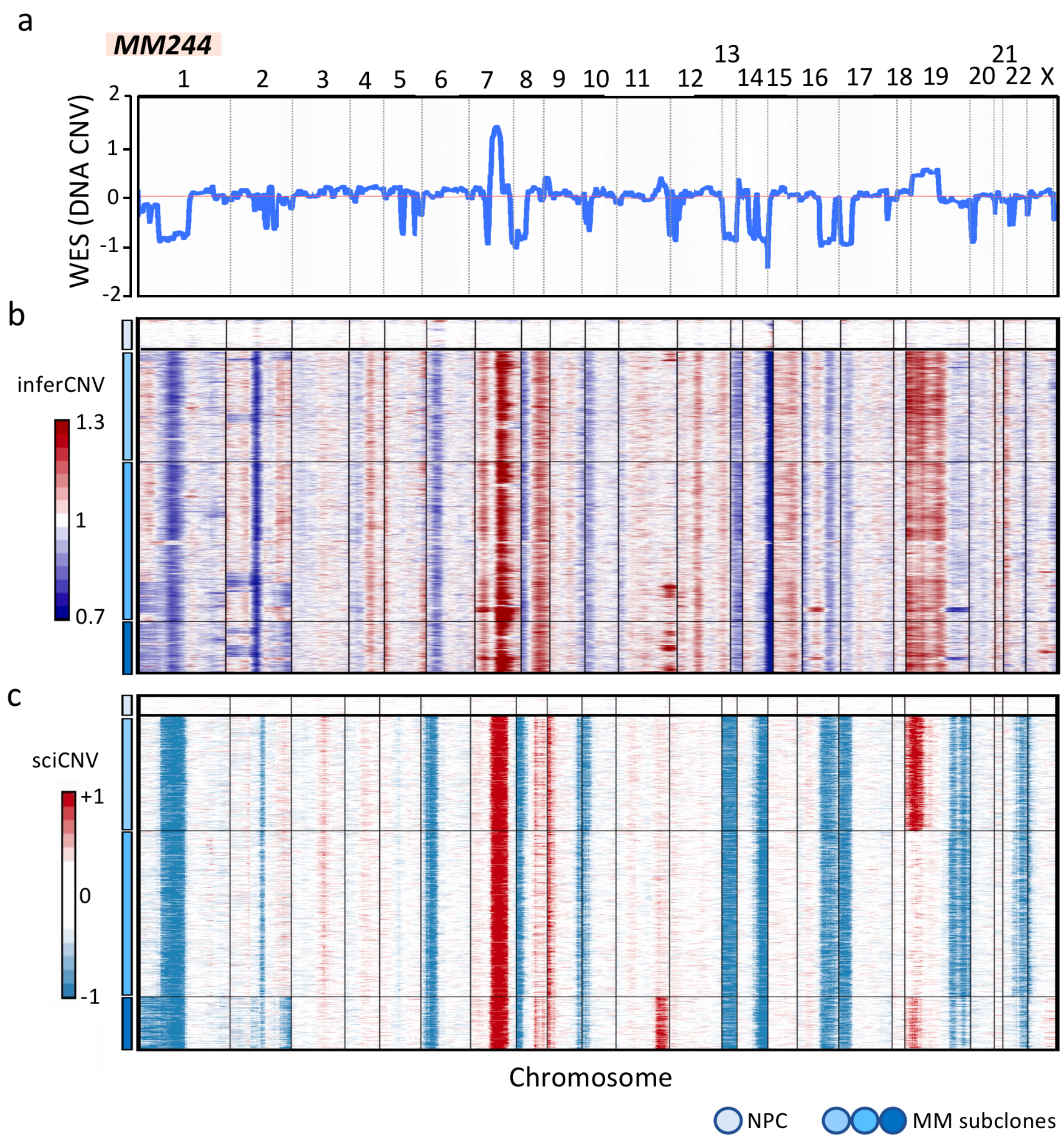


**Figure S25. sciCNV compares favorably with inferCNV in predicting single cell CNV using scRNA-seq – part 3 (sample MM244).**

Single cell CNV profiles, derived by sciCNV or inferCNV for hundreds of individual MM cells using scRNA-seq data, are compared with the tumor bulk CNV profile obtained by DNA-sequencing of pooled tumor cells.

**a.** The tumor bulk CNV profile for sample MM244, generated by DNA whole exome sequencing.

**b-c.** The CNV profiles of individual plasma cells (n=3569) from MM patient sample MM244, inferred from scRNA-seq using inferCNV (b) or sciCNV (c). The profiles of normal plasma cells (NPCs)(n=205) are shown as controls.

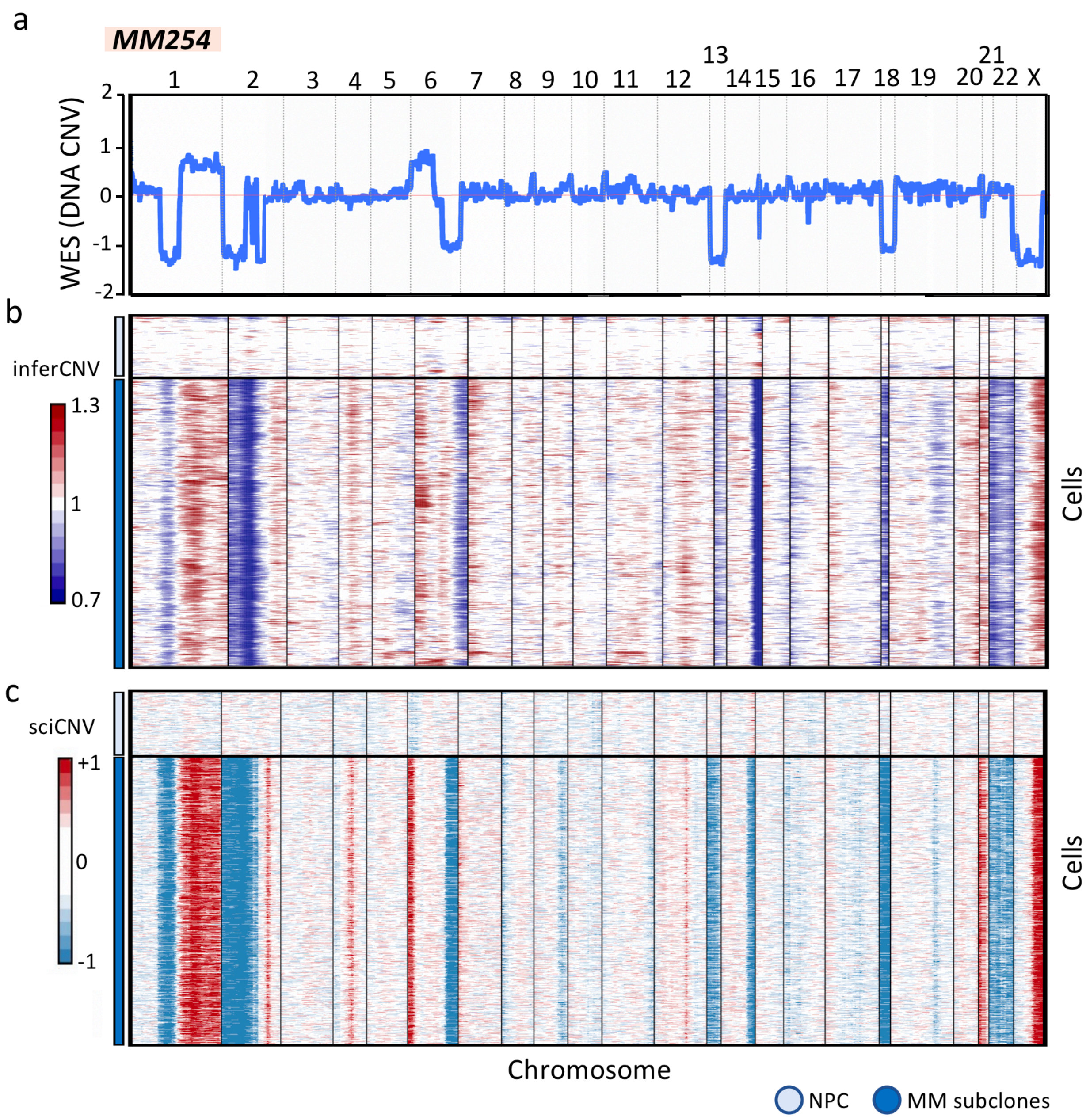


**Figure S26. sciCNV compares favorably with inferCNV in predicting single cell CNV using scRNA-seq – part 4 (sample MM254).**

Single cell CNV profiles, derived by sciCNV or inferCNV for hundreds of individual MM cells using scRNA-seq data, are compared with the tumor bulk CNV profile obtained by DNA-sequencing of pooled tumor cells.

**a.** The tumor bulk CNV profile for sample MM254, generated by DNA whole exome sequencing.

**b-c.** The CNV profiles of individual plasma cells (n=649) from MM patient sample MM254, inferred from scRNA-seq using inferCNV (b) or sciCNV (c). The profiles of normal plasma cells (NPCs)(n=205) are shown as controls.

**MM199**

**GSEA: chromosome position**
Subclone with +8q22-24 (vs no +8q)

| | GS follow link to MSigDB | GS DETAILS | SIZE | ES | NES | NOM p-val | FDR q-val | FWER p-val | RANK AT MAX |
|---|---|---|---|---|---|---|---|---|---|
| 1 | CHR8Q24 | Details ... | 102 | 0.66 | 3.12 | 0.000 | 0.000 | 0.000 | 2066 |
| 2 | CHR8Q22 | Details ... | 42 | 0.61 | 2.44 | 0.000 | 0.000 | 0.000 | 1652 |
| 3 | CHR2P22 | Details ... | 31 | 0.50 | 1.84 | 0.004 | 0.080 | 0.219 | 2576 |
| 4 | CHR6Q21 | Details ... | 42 | 0.44 | 1.75 | 0.006 | 0.131 | 0.423 | 1994 |
| 5 | CHR18Q11 | Details ... | 15 | 0.57 | 1.72 | 0.010 | 0.141 | 0.525 | 2106 |
| 6 | CHR4Q22 | Details ... | 17 | 0.51 | 1.57 | 0.036 | 0.369 | 0.895 | 4067 |
| 7 | CHR17Q25 | Details ... | 117 | 0.31 | 1.52 | 0.013 | 0.469 | 0.963 | 3910 |
| 8 | CHR16Q23 | Details ... | 20 | 0.47 | 1.51 | 0.044 | 0.435 | 0.973 | 1550 |
| 9 | CHR19P12 | Details ... | 24 | 0.44 | 1.50 | 0.049 | 0.404 | 0.979 | 2791 |
| 10 | CHR6Q23 | Details ... | 21 | 0.45 | 1.50 | 0.043 | 0.373 | 0.980 | 282 |

**GSEA: Hallmark gene sets**
Subclone with +8q22-24 (vs no +8q)

| | GS follow link to MSigDB | GS DETAILS | SIZE | ES | NES | NOM p-val | FDR q-val | FWER p-val | RANK AT MAX |
|---|---|---|---|---|---|---|---|---|---|
| 1 | HALLMARK_MYC_TARGETS_V2 | Details ... | 58 | 0.52 | 2.19 | 0.000 | 0.000 | 0.000 | 3260 |
| 2 | HALLMARK_UV_RESPONSE_UP | Details ... | 125 | 0.29 | 1.43 | 0.014 | 0.437 | 0.483 | 2632 |
| 3 | HALLMARK_MYOGENESIS | Details ... | 108 | 0.27 | 1.33 | 0.046 | 0.628 | 0.789 | 3583 |
| 4 | HALLMARK_ESTROGEN_RESPONSE_EARLY | Details ... | 132 | 0.26 | 1.28 | 0.071 | 0.684 | 0.901 | 2789 |
| 5 | HALLMARK_INTERFERON_GAMMA_RESPONSE | Details ... | 175 | 0.24 | 1.25 | 0.075 | 0.680 | 0.942 | 2978 |
| 6 | HALLMARK_MYC_TARGETS_V1 | Details ... | 199 | 0.23 | 1.24 | 0.060 | 0.596 | 0.952 | 2918 |
| 7 | HALLMARK_COMPLEMENT | Details ... | 139 | 0.25 | 1.22 | 0.106 | 0.575 | 0.963 | 2420 |
| 8 | HALLMARK_E2F_TARGETS | Details ... | 198 | 0.23 | 1.20 | 0.105 | 0.592 | 0.978 | 2298 |
| 9 | HALLMARK_G2M_CHECKPOINT | Details ... | 191 | 0.23 | 1.17 | 0.133 | 0.628 | 0.989 | 2286 |
| 10 | HALLMARK_FATTY_ACID_METABOLISM | Details ... | 127 | 0.23 | 1.16 | 0.173 | 0.610 | 0.991 | 2768 |

**GSEA: Curated gene sets**
Subclone with +8q22-24 (vs no +8q)

| | follow | GS DETAILS | SIZE | ES | NES | NOM p-val | FDR q-val | FWER p-val |
|---|---|---|---|---|---|---|---|---|
| 1 | NIKOLSKY_BREAST_CANCER_8Q23_Q24_AMPLICON | Details ... | 108 | 0.66 | 3.13 | 0.000 | 0.000 | 0.000 |
| 2 | CLIMENT_BREAST_CANCER_COPY_NUMBER_UP | Details ... | 15 | 0.76 | 2.30 | 0.000 | 0.003 | 0.006 |
| 3 | REACTOME_PEPTIDE_CHAIN_ELONGATION | Details ... | 83 | 0.50 | 2.24 | 0.000 | 0.005 | 0.014 |
| 4 | KEGG_RIBOSOME | Details ... | 85 | 0.48 | 2.19 | 0.000 | 0.009 | 0.032 |
| 5 | REACTOME_3_UTR_MEDIATED_TRANSLATIONAL_REGULATION | Details ... | 103 | 0.44 | 2.09 | 0.000 | 0.039 | 0.167 |
| 6 | PID_IL3_PATHWAY | Details ... | 24 | 0.56 | 1.94 | 0.000 | 0.184 | 0.649 |
| 7 | NIKOLSKY_BREAST_CANCER_8Q12_Q22_AMPLICON | Details ... | 86 | 0.42 | 1.91 | 0.000 | 0.211 | 0.740 |
| 8 | SUNG_METASTASIS_STROMA_DN | Details ... | 38 | 0.50 | 1.90 | 0.000 | 0.207 | 0.790 |
| 9 | TAKEDA_TARGETS_OF_NUP98_HOXA9_FUSION_3D_DN | Details ... | 15 | 0.63 | 1.88 | 0.004 | 0.228 | 0.852 |
| 10 | BIOCARTA_TEL_PATHWAY | Details ... | 16 | 0.60 | 1.86 | 0.002 | 0.246 | 0.896 |

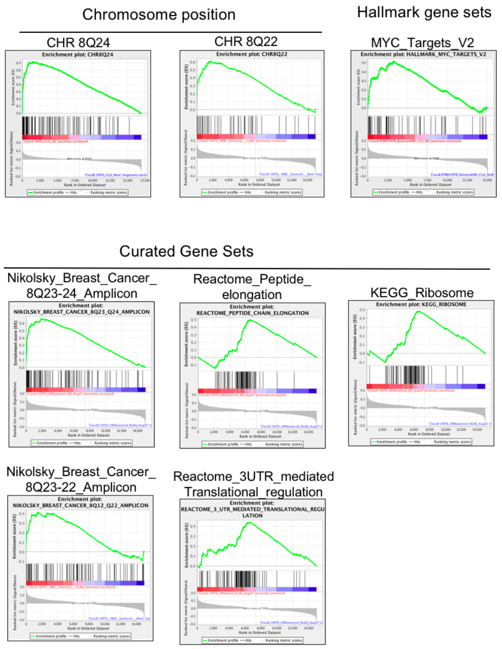


**Figure S27. Summary of gene sets enriched in +8q23-24 cells in MM199**

MM199 cells with and without 8q22-24 gain were examined by GSEA for divergent expression of chromosome position (n=215), hallmark (n=49) and curated (n=3,303) gene sets. For the analysis, the paired subclones in each sample with and without 8q copy number were subsampled to yield subpopulations of cells with matching transcript depth (UMI/cell) distribution. The top 10 enriched gene sets in +8q cells are shown for each sample and each analysis. Significant results (with FWER p<0.05) are highlighted in red and important near significant results (with nominal p<0.05 and FWER>0.05) are shown in yellow. Similar top-ranked enrichment results, from 3303 curated gene sets, between the two samples are joined by a dashed line. Gene set enrichment details are shown in the plots below.

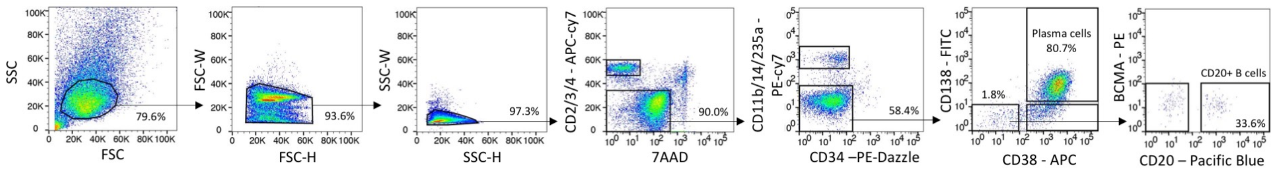


**Figures S28. Sorting strategy for mature plasma cells and Cd20+ B cells.**

Representative flow cytometry plots showing sorting strategy for plasma cells and B cells using lymphoid cell characteristics, doublet exclusion, non-viable cell exclusion, exclusion of cells expressing non B cell-lineage markers, and positive selection for CD38 and CD138 or for CD20.
